# Rare variants contribute disproportionately to quantitative trait variation in yeast

**DOI:** 10.1101/607291

**Authors:** Joshua S Bloom, James Boocock, Sebastian Treusch, Meru J Sadhu, Laura Day, Holly Oates-Barker, Leonid Kruglyak

## Abstract

A detailed understanding of the sources of heritable variation is a central goal of modern genetics. Genome-wide association studies (GWAS) in humans^1^ have implicated tens of thousands of DNA sequence variants in disease risk and quantitative trait variation, but these variants fail to account for the entire heritability of diseases and traits. GWAS have by design focused on common DNA sequence variants; however, recent studies underscore the likely importance of the contribution of rare variants to heritable variation^2^. Further, finding the genes that underlie the GWAS signals remains a major challenge. Here, we use a unique model system to disentangle the contributions of common and rare variants to a large number of quantitative traits. We generated large crosses among 16 diverse yeast strains and identified thousands of quantitative trait loci (QTLs) that explain most of the heritable variation in 38 traits. We combined our results with sequencing data for 1,011 yeast isolates^3^ to decouple variant effect size estimation from allele frequency and showed that rare variants make a disproportionate contribution to trait variation as a consequence of their larger effect sizes. Evolutionary analyses revealed that this contribution is driven by rare variants that arose recently, that such variants are more likely to decrease fitness, and that negative selection has shaped the relationship between variant frequency and effect size. Finally, we leveraged the structure of the crosses to resolve hundreds of QTLs to single genes. These results refine our understanding of trait variation at the population level and suggest that studies of rare variants are a fertile ground for discovery of genetic effects.

## Introduction

How variants with different population frequencies contribute to trait variation is a central question in genetics. Theoretical considerations^4–8^ and previous results in yeast^9^, humans^10–12^, and other species^13^ suggest that rare variants should have larger effect sizes, or, equivalently, that variants implicated in trait variation should be shifted to lower frequencies relative to all variants. Variance partitioning by allele frequency has revealed appreciable contributions of lower-frequency variants to heritability of complex traits in humans, such as prostate cancer^14^ height^2,15^, and body mass index^2^. However, a direct comprehensive comparison of the effects of rare and common variants has been lacking in humans owing to low statistical power to map rare variants and confounding between effect size and allele frequency. Here we report a comprehensive study in yeast designed to overcome these limitations. We built a panel of approximately 14,000 segregants from crosses between 16 diverse yeast strains, mapped thousands of QTLs that account for most of the heritable variation in 38 quantitative traits, and measured the QTL effect sizes. We then estimated the allele frequencies of the underlying variants in a collection of over 1,000 sequenced yeast isolates from around the world^3^. Analysis of these large complementary data sets enabled us to examine the relationship between QTL effect sizes and variant frequency, characterize the genetic architecture of quantitative traits on a population scale, and improve mapping resolution, in many cases to single genes.

## Results

To investigate the genetic basis of quantitative traits in the yeast population, we selected 16 highly diverse *S. cerevisiae* strains that capture much of the known genetic diversity of this species. Specifically, they contain both alleles at 82% of biallelic SNPs and small indels observed at minor allele frequency > 5% in a collection of 1,011 *S. cerevisiae* strains^3^. We sequenced the 16 strains to high coverage in order to obtain a comprehensive set of genetic variants. We constructed a panel of 13,950 individual recombinant haploid yeast segregants by crossing each parental strain to two different strains and collecting an average of 872 progeny per cross (Fig. 1, Supplementary Table 1). We genotyped these segregants by highly multiplexed whole-genome sequencing, with median 2.3-fold coverage per base per individual. Genotypes were called at 298,979 genetic variants, with an average of 71,117 genetic variants segregating in a single cross. We phenotyped each segregant for 38 fitness traits in duplicate by automated growth assays and quantitative imaging (Methods). The resulting genotype-by-phenotype matrix (over half a million phenotypic measurements and 158 billion combinations of genotype and phenotype) formed the basis for all downstream analyses.

**Figure 1.**
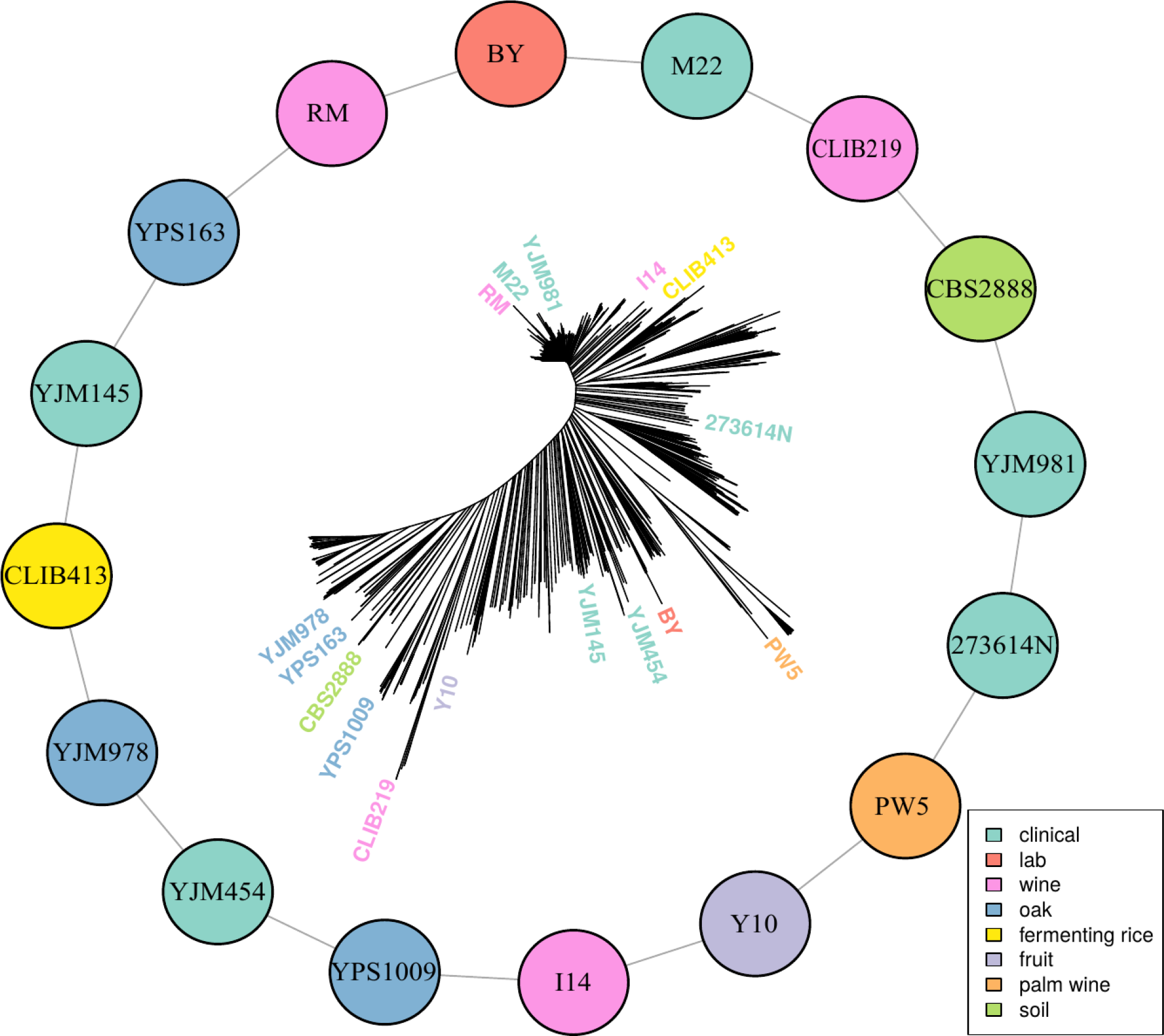
Multiparental cross design with 16 diverse progenitor yeast strains. 16 parental strains were chosen to represent the diversity of the *S. cerevisiae* population, as illustrated by their positions on a neighbor-joining tree based on 1,011 sequenced isolates^3^. These strains were crossed in a single round-robin design, with each strain crossed to two other strains, as depicted by lines connecting the colored circles. Colors indicate the ecological origins of the parental strains.

We used a variance components model^16–18^ to show that, on average, additive genetic effects accounted for just over half of the total phenotypic variance, while pairwise genetic interactions accounted for 8%, approximately 1/6 as much as additive effects (Fig. 2 inset, Supplementary Fig. 1, Supplementary Table 2). We carried out QTL mapping to find the specific loci contributing additively to trait variation. We used a joint mapping approach that leverages information across the entire panel of 13,950 segregants (Methods). We mapped 4,552 QTLs at a false discovery rate (FDR) of 5%, with an average of 120 (range 52-195) QTLs per trait (Supplementary Fig. 2, Supplementary Table 3). The detected QTLs explain a median of 73% of the additive heritability per trait and cross, showing that we can account for most of the genetic contribution to trait variation with specific loci (Fig. 2, Supplementary Table 2). We complemented the joint analysis with QTL mapping within each cross and found a median of 12 QTLs per trait at the same FDR of 5%. The detected loci explained a median of 68% of the additive heritability (Supplementary Table 2). The joint analysis was more powerful, explaining an additional 5% of trait variance and uncovering 458 QTLs not detected within individual crosses. Consistent with the higher statistical power of the joint analysis, these additional QTLs had smaller effect sizes (median of 0.071 SD units vs 0.083 SD units; Wilcoxon rank sum test W=1e6, p=9e-5).

**Figure 2.**
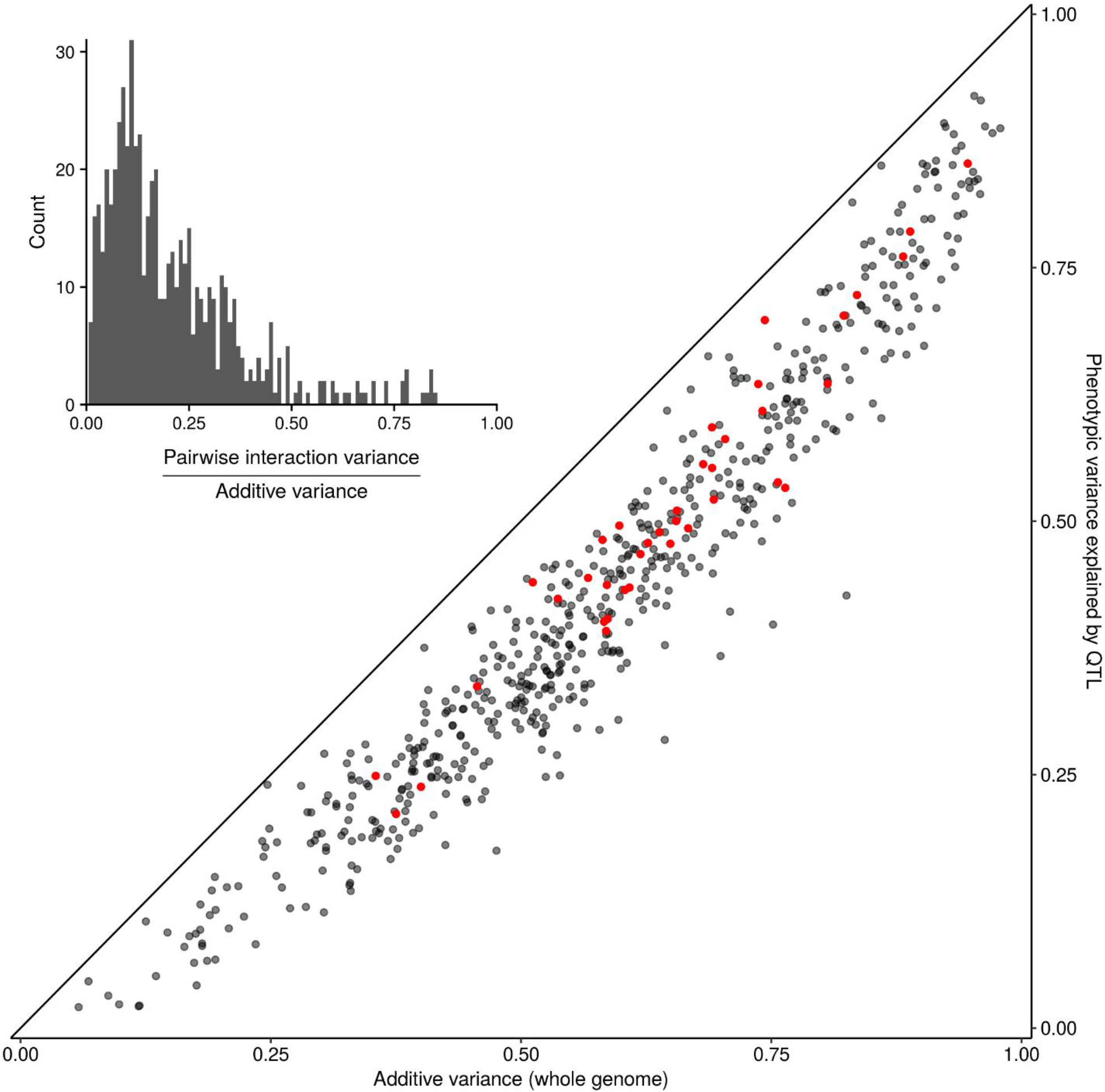
Most heritable variation is explained by detected QTL. Whole-genome estimates of additive genetic variance (X-axis) are plotted against cross-validated estimates of trait variance explained by detected QTLs (Y-axis) for each trait-cross combination. Red points show values for the BYxRM cross. The diagonal line corresponds to 100% of trait variance explained by detected QTL and is shown as a visual guide. (Inset) A histogram of the ratio of non-additive to additive genetic variance for each trait-cross combination estimated by a variance component model.

To investigate the relationship between variant frequency and QTL effects, we focused on biallelic variants observed in our panel whose frequency could be measured in a large collection of 1,011 sequenced yeast strains. Based on their minor allele frequency (MAF) in this collection, we designated variants as rare (MAF < 0.01) or common (MAF > 0.01). By this definition, 27.8% of biallelic variants in our study were rare. For each trait, we computed the relative fraction of variation explained by these two categories of variants in the segregant panel (Methods)^15^. Across all traits, the median contribution of rare variants was 51.7%, despite the fact that they constituted only 27.8% of all variants, and that a rare variant is expected to explain less variance than a common one with the same allelic effect size. These results are consistent with rare variants having larger effect sizes and making a disproportionate contribution to trait variation. Comparing different traits, we saw a wide range of the relative contribution of rare variants, from almost none for growth in the presence of copper sulfate and lithium chloride to over 75% for growth in the presence of cadmium chloride, in low pH, at high temperature, and on minimal medium (Fig. 3a, Supplementary Fig. 3, Supplementary Table 4). The results for copper sulfate and lithium chloride are consistent with GWAS for these traits in the 1,011 sequenced yeast strains—these two traits had the most phenotypic variance explained by detected GWAS loci, which inherently correspond to common variants, with large contributions coming from known common copy-number variation at the *CUP* and *ENA* loci, respectively^3^.

**Figure 3.**
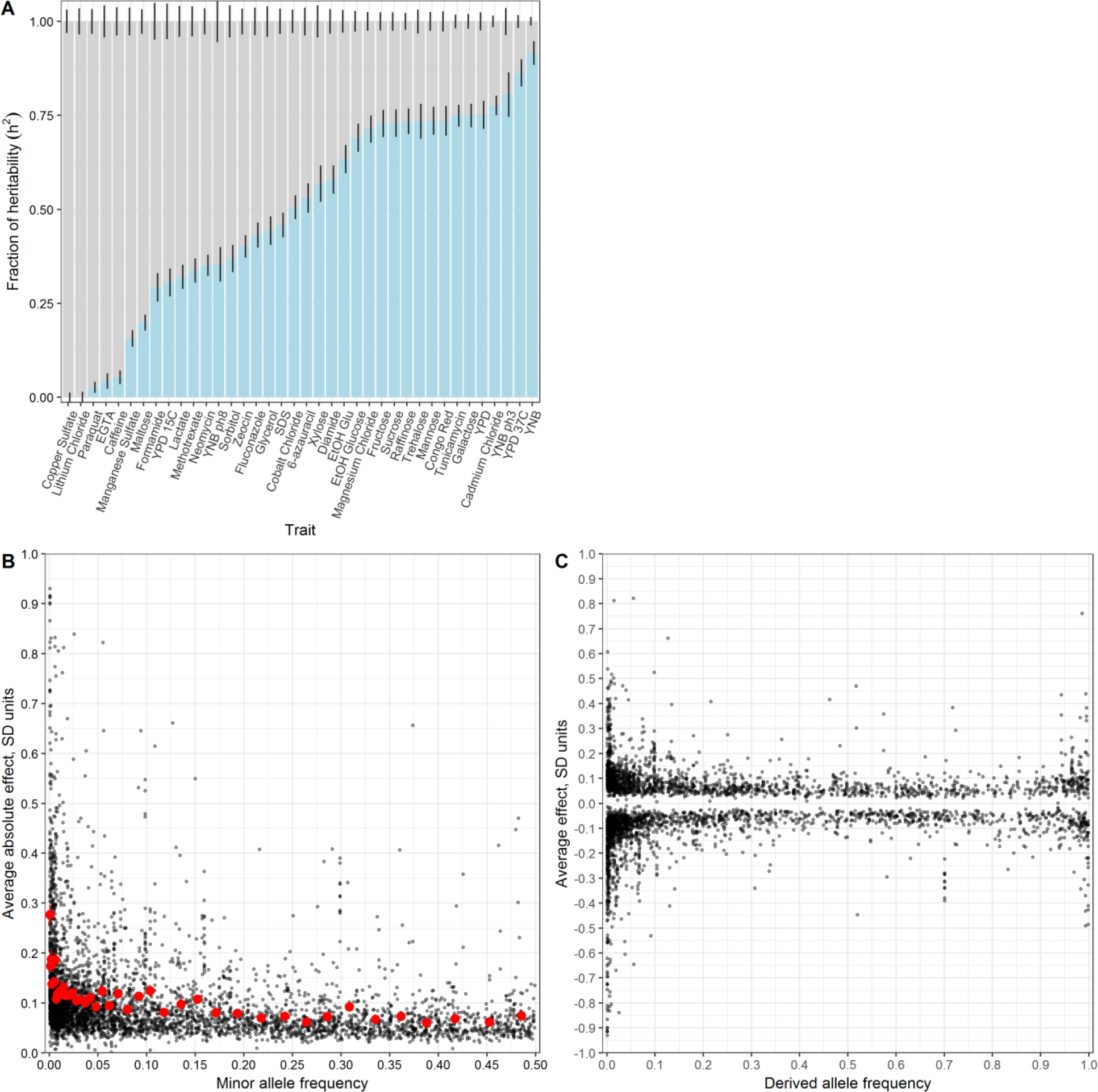
Effect size and contribution to trait variation of rare and common variants. (A) Stacked bar plots of additive genetic variance explained by rare (blue) and common (grey) variants. Error bars show +/− s.e. (B) Minor allele frequency (X-axis) of the lead variant at each QTL^3^ is plotted against QTL effect size (Y-axis). Red points show mean QTL effect size for groups of approximately 100 variants binned by allele frequency. Error bars show +/− s.e.m. (C) Frequency of the derived allele of each QTL lead variant (X-axis), based on comparison with *S. paradoxus*, is plotted against QTL effect size (Y-axis).

In a complementary analysis, we investigated the relationship between the allele frequency of the lead variant at each QTL and the corresponding QTL effect size. Although the lead variant is not necessarily causal, in our study it is likely to be of similar frequency as the causal variant, and a simulation analysis showed that this approach largely preserves the relationship between frequency and effect size (Supplementary Fig. 4). Most QTLs had small effects (64% of QTLs had effects less than 0.1 SD units) and most lead variants were common (78%), consistent with previous linkage and association studies. We observed that QTLs with large effects were highly enriched for rare variants, and conversely, that rare variants were highly enriched for large effect sizes (Fig. 3b, Supplementary Fig. 5). For instance, among QTLs with an absolute effect of at least 0.3 SD units, 145 of the corresponding lead variants were rare and only 90 were common. Rare variants were 6.7 times more likely to have an effect greater than 0.3 SD (Supplementary Table 3, Fisher’s exact test, p<2e-16). Theoretical population genetics models show that for traits under negative selection, variant effect size is expected to be a decreasing function of minor allele frequency^4,5^. We empirically observe this relationship in our data for most of the traits examined, providing evidence that they have evolved under negative selection in the yeast population (Supplementary Fig. 6).

**Figure 4.**
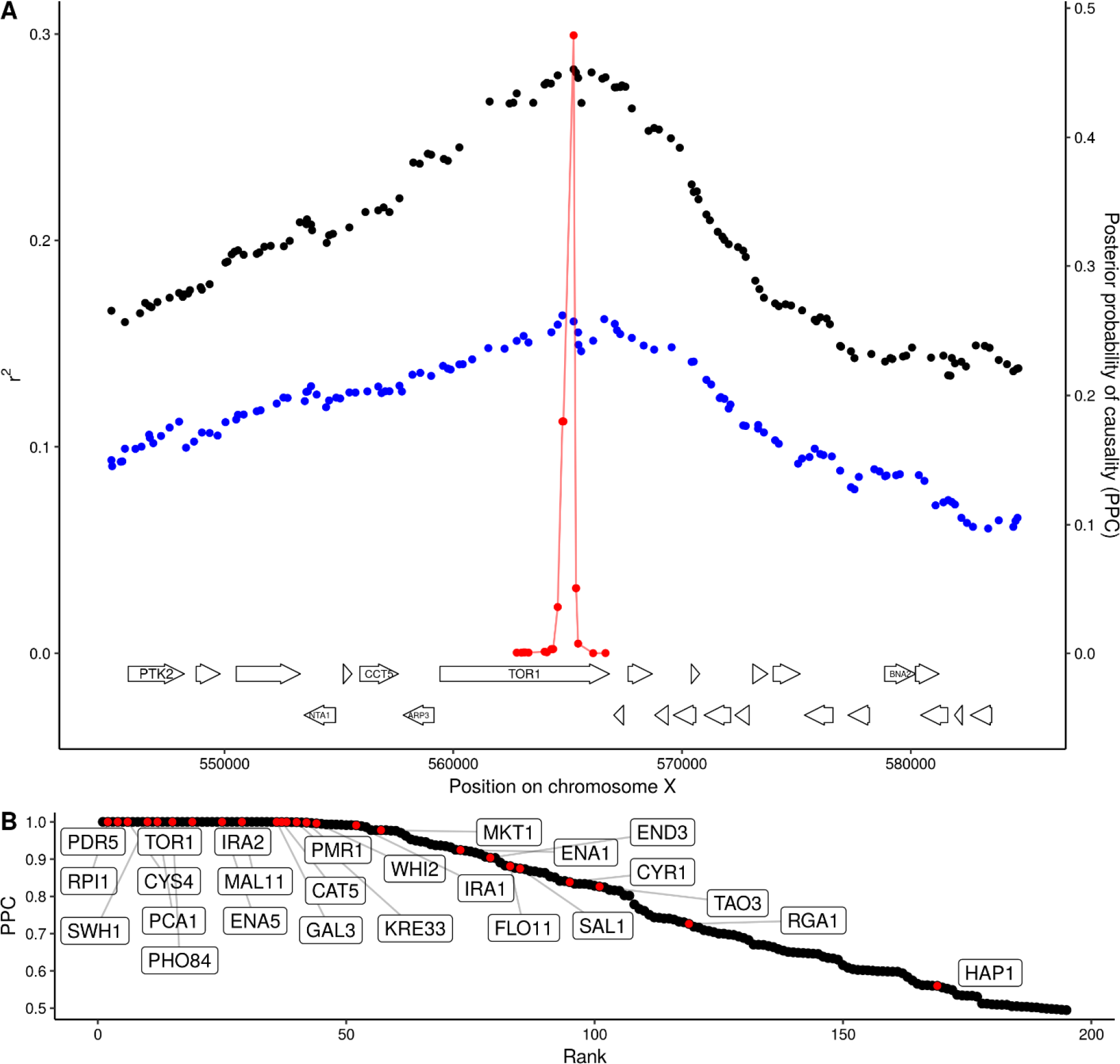
QTL fine-mapping at gene-level resolution. (A) Statistical fine-mapping of a QTL for growth in the presence of caffeine. Genetic mapping signal, shown as the coefficient of determination between genotype and phenotype (Y-axis, left), is plotted against genome position (X-axis) for crosses between 273614N and YJM981 (black) and YJM981 and CBS2888 (blue). The posterior probability of causality (PPC), plotted in red (Y-axis, right), localizes the QTL to a portion of the gene TOR1. (B) PPC is shown as black dots for 195 genes identified as causal at an FDR of 20%, sorted by PPC. Genes containing natural variants that have been experimentally validated as causal for trait variation in prior studies^21–25^ are shown in red and labelled with gene names.

The existence of a close sister species of *S. cerevisiae*—*S. paradoxus*—allowed us to distinguish rare variants by their ancestral state. Variants that share the major allele with *S. paradoxus* are more likely to have arisen in the *S. cerevisiae* population recently than those that share the minor allele with *S. paradoxus*. We classified low-frequency variants as recent or ancient according to whether their major or minor allele was shared with *S. paradoxus*, respectively. Recently arising deleterious alleles have had less time to be purged by negative selection, and therefore recent variants are expected to have stronger effects on gene function, and hence manifest as QTLs with larger effects. Consistent with the expectation above, we observed that recent variants were 1.8 times more likely than ancient variants to have an effect size greater than 0.1 SD units (Fisher’s exact test p=9e-5) (Fig. 3c). We further examined the direction of QTL effects and found that recent variants were 1.5 times more likely to decrease fitness (Fisher’s exact test p=8e-3). Strikingly, no ancient variant decreased fitness by more than 0.5 SD units, whereas 41 recent variants did (Fisher’s exact test p=7e-3).

An understanding of trait variation at the level of molecular mechanisms requires narrowing QTLs to the underlying causal genes. Such fine-mapping is a challenge because genetic linkage causes variants across an extended region to show mapping signals of similar strength. Statistical fine-mapping aims to address this challenge by estimating the probability that each variant within a QTL region is causal based on the precise pattern of genotype-phenotype correlations^19–21^. Our crossing design enables us to obtain higher resolution for QTLs observed in two crosses that share a parent strain by looking for consistent inheritance patterns in both. Specifically, we focused on QTLs with effects greater than 0.14 SD units and used a Bayesian framework^20^ to compute the posterior probability that each variant is causal (Fig. 4a). We then aggregated these probabilities to obtain causality scores for each gene in a QTL. With this approach, we resolved 427 QTLs to single causal genes at an FDR of 20%. Because some QTLs have pleiotropic effects on multiple traits, this gene set contains 195 unique genes, greatly expanding the repertoire of causal genes in yeast. We searched the literature and found that 26 of the 195 genes identified here are supported by previous experimental evidence as causal for yeast trait variation^21–25^ (Fig. 4b, Supplementary Table 5). At a more stringent FDR of 5%, we found 105 unique causal genes, which included 24 of the 26 genes with experimental evidence.

Causal genes were highly enriched for GO terms related to the plasma membrane (45 of 522, 16.5 expected, q=.1.8e-7), metal ion transport (13 of 83, 2.6 expected, q=0.0009), and positive regulation of nitrogen compound biosynthesis (28 of 393, 12.5 expected, q=0.0076) (Supplementary Table 5). Strikingly, 5 of the 6 genes involved in cAMP biosynthesis were identified as causal (*IRA1*, *IRA2*, *BCY1*, *CYR1*, and *RAS1;* 0.19 expected, q=0.0002). Additional genes in the RAS/cAMP signaling pathway were also identified as causal, including *GPR1*, which is involved in glucose sensing, *SRV2*, which binds adenylate cyclase, and *RHO3*, which encodes a RAS-like GTPase. In yeast, the RAS/cAMP pathway regulates cell cycle progression, metabolism, and stress resistance^26^. Variation in many of these genes influenced growth on alternative carbon sources. We hypothesize that the yeast population contains abundant functional variation in genes that regulate the switch from glucose to alternative carbon sources through the RAS/cAMP pathway.

## Discussion

We previously used a cross between lab (BY) and vineyard strains (RM) of yeast to show that the majority of heritable phenotypic differences arise from additive genetic effects, and we were able to detect, at genome-wide significance, specific loci that together account for the majority of quantitative trait variation^18,27^. It has been argued that the BY lab reference strain S288c used in those and many other yeast studies is genetically and phenotypically atypical compared to other yeast isolates^28^. Our results here, obtained from crosses among 16 diverse strains, generalize these findings to the *S. cerevisiae* population and show that S288c is not exceptional from the standpoint of genetic variation and quantitative traits. We discovered over 4,500 quantitative trait loci (QTLs) that influence yeast growth in a wide variety of conditions. These loci likely capture the majority of common variants that segregate in *S. cerevisiae* and have appreciable phenotypic effects on growth. We were able to localize approximately 8% of the QTLs to single genes based on genetic mapping information alone. Interestingly, these genes cluster in specific functional categories and pathways, suggesting that different strains of *S. cerevisiae* may have evolved different strategies for nutrient sensing and response as a function of specializing in particular environmental niches^29^. In addition to the findings described here, we anticipate that our data set will be a useful resource for further dissecting the genetic basis of trait variation at the gene and variant level, and for evaluating statistical methods aimed at inferring causal genes and variants. In particular, the set of loci and genes identified here provides an ideal starting point for massively parallel editing experiments that directly test the phenotypic consequences of sequence variants^30^.

By combining our results with deep population sequencing in yeast^3^ we were able to examine the contributions of variants in different frequency classes to trait variation. We observed a broad range of genetic architectures across the traits studied here, with variation in some traits dominated by common variants, while variation in others is mostly explained by rare variants. Overall, rare variants made a disproportionate contribution to trait variation as a consequence of their larger effect sizes. A complementary mapping approach in an overlapping set of yeast isolates also revealed enrichment of rare variants with larger effect sizes (Fournier and Schacherer, personal communication). These results are consistent with the finding from GWAS that common variants have small effects, as well as with linkage studies that find rare variants with large effect sizes. Our study design also revealed a substantial component of genetic variation—variants with low allele frequency and small effect size—that has been refractory to discovery in humans because both GWAS and linkage studies lack statistical power to detect this class of variants. Recent work in humans has suggested that rare variants account for a substantial fraction of heritability of complex traits and diseases^2^. Our study presents a more direct and fine-grained view of this component of trait variation and implies that larger sample sizes and more complete genotype information will be needed for more comprehensive studies in other systems.

## Acknowledgements

We thank Bogdan Pasaniuc, Frank W Albert, Olga T Schubert, Liangke Gou, Tzitziki Lemus Vergara, Matthieu Delcourt, Longhua Guo, and Eyal Ben-David for helpful manuscript feedback and edits. We thank Semyon Kruglyak, Erich Jaeger, and Illumina for performing synthetic long-read sequencing of the parental yeast strains. This work was supported by funding from the Howard Hughes Medical Institute (to LK) and NIH grant R01GM102308 (to LK).

## Author contributions

JSB performed the experiments with assistance from ST, LD, and HOB. MS provided helpful discussions. JSB analyzed the data with assistance from JB. LK supervised the project. JSB and LK wrote the manuscript.

## References

1. Visscher, P. M. et al. 10 Years of GWAS Discovery: Biology, Function, and Translation. Am. J. Hum. Genet. 101, 5–22 (2017).

2. Wainschtein, P. et al. Recovery of trait heritability from whole genome sequence data. bioRxiv 588020 (2019). doi:10.1101/588020

3. Peter, J. et al. Genome evolution across 1,011 Saccharomyces cerevisiae isolates. Nature 556, 339–344 (2018).

4. Pritchard, J. K. Are Rare Variants Responsible for Susceptibility to Complex Diseases? Am J Hum Genet 69, 124–137 (2001).

5. Eyre-Walker, A. Genetic architecture of a complex trait and its implications for fitness and genome-wide association studies. PNAS 107, 1752–1756 (2010).

6. Gibson, G. Rare and Common Variants: Twenty arguments. Nat Rev Genet 13, 135–145 (2012).

7. Zuk, O. et al. Searching for missing heritability: designing rare variant association studies. Proc. Natl. Acad. Sci. U.S.A. 111, E455–464 (2014).

8. Robinson, M. R., Wray, N. R. & Visscher, P. M. Explaining additional genetic variation in complex traits. Trends Genet 30, 124–132 (2014).

9. Ehrenreich, I. M. et al. Genetic Architecture of Highly Complex Chemical Resistance Traits across Four Yeast Strains. PLOS Genetics 8, e1002570 (2012).

10. Ganna, A. et al. Quantifying the Impact of Rare and Ultra-rare Coding Variation across the Phenotypic Spectrum. Am. J. Hum. Genet. 102, 1204–1211 (2018).

11. Lek, M. et al. Analysis of protein-coding genetic variation in 60,706 humans. Nature 536, 285–291 (2016).

12. Marouli, E. et al. Rare and low-frequency coding variants alter human adult height. Nature 542, 186–190 (2017).

13. Wallace, J. G. et al. Association Mapping across Numerous Traits Reveals Patterns of Functional Variation in Maize. PLOS Genetics 10, e1004845 (2014).

14. Mancuso, N. et al. The contribution of rare variation to prostate cancer heritability. Nat. Genet. 48, 30–35 (2016).

15. Yang, J. et al. Genetic variance estimation with imputed variants finds negligible missing heritability for human height and body mass index. Nat. Genet. 47, 1114–1120 (2015).

16. Yang, J. et al. Common SNPs explain a large proportion of the heritability for human height. Nature Genetics 42, 565–569 (2010).

17. Campos, G. de los, Sorensen, D. & Gianola, D. Genomic Heritability: What Is It? PLOS Genetics 11, e1005048 (2015).

18. Bloom, J. S. et al. Genetic interactions contribute less than additive effects to quantitative trait variation in yeast. Nature Communications 6, ncomms9712 (2015).

19. Pasaniuc, B. & Price, A. L. Dissecting the genetics of complex traits using summary association statistics. Nat Rev Genet 18, 117–127 (2017).

20. Farh, K. K.-H. et al. Genetic and epigenetic fine mapping of causal autoimmune disease variants. Nature 518, 337–343 (2015).

21. Treusch, S., Albert, F. W., Bloom, J. S., Kotenko, I. E. & Kruglyak, L. Genetic Mapping of MAPK-Mediated Complex Traits Across S. cerevisiae. PLOS Genetics 11, e1004913 (2015).

22. Fay, J. C. The molecular basis of phenotypic variation in yeast. Curr. Opin. Genet. Dev. 23, 672–677 (2013).

23. Wang, X. & Kruglyak, L. Genetic Basis of Haloperidol Resistance in Saccharomyces cerevisiae Is Complex and Dose Dependent. PLOS Genetics 10, e1004894 (2014).

24. Jerison, E. R. et al. Genetic variation in adaptability and pleiotropy in budding yeast. Elife 6, (2017).

25. Sadhu, M. J., Bloom, J. S., Day, L. & Kruglyak, L. CRISPR-directed mitotic recombination enables genetic mapping without crosses. Science 352, 1113–1116 (2016).

26. Tisi, R., Belotti, F. & Martegani, E. Yeast as a Model for Ras Signalling. in Ras Signaling: Methods and Protocols (eds. Trabalzini, L. & Retta, S. F.) 359–390 (Humana Press, 2014). doi:10.1007/978-1-62703-791-4_23

27. Bloom, J. S., Ehrenreich, I. M., Loo, W. T., Lite, T.-L. V. & Kruglyak, L. Finding the sources of missing heritability in a yeast cross. Nature 494, 234–237 (2013).

28. Warringer, J. et al. Trait Variation in Yeast Is Defined by Population History. PLOS Genetics 7, e1002111 (2011).

29. Chantranupong, L., Wolfson, R. L. & Sabatini, D. M. Nutrient Sensing Mechanisms Across Evolution. Cell 161, 67–83 (2015).

30. Shendure, J. & Fields, S. Massively Parallel Genetics. Genetics 203, 617–619 (2016).

