## Supplementary Materials for "Rare variants contribute disproportionately to quantitative trait variation in yeast"

### Materials and Methods

Unless otherwise specified, all computational analyses were performed in R (v3.4.4). Analysis code and processing scripts are available at https://github.com/joshsbloom/yeast-16-parents. The version numbers of R packages used are listed in this repository.

#### Short-read and synthetic long read sequencing of parental strains

Parental genotypes were obtained by deep (> 100X) paired-end sequencing of the 16 parental strains. A VCF file containing SNPs and small indels was generated for the parents using bwa (v0.7.1)[1] to align to the sacCer3 reference[2], Picard (v2.12.2)[3] to remove PCR duplicates, and the GATK HaplotypeCaller (v3.8)[4] with expected sample ploidy set to 1. A separate pipeline was developed to leverage additional synthetic long-reads (Illumina/Moleculo) to identify larger structural variants in the parents. Briefly, synthetic long-read assemblies were filtered to only include scaffolds greater than 10kb. Scaffolds were corrected with our short-read data using Pilon[5]. CNVs were discovered using custom scripts modified from scripts originally used to generate calls for testing LUMPY[6]. CNVs were genotyped in all parents using the approach presented in SVTyper[7]. Scripts associated with the CNV detection pipeline are available at https://github.com/theboocock/long_read_cnv.

#### Construction of haploid segregant panels

Segregants for the BYxRM cross and YPS163xYJM145 cross were obtained by sporulation of the hybrid diploid parents for 5-7 days in SPO++ sporulation medium (http://dunham.gs.washington.edu/sporulationdissection.htm) and tetrad dissection using the MSM 400 dissection microscope (Singer Instrument Company Ltd.). Four-spore tetrads were retained. For BYxRM one segregant was randomly chosen per tetrad[8]. For YPX163xYJM145 all segregants from ~250 tetrads were used. For all other crosses, the hybrid diploids were were pre-grown in YPD with either G418 or cloNat, depending on which fluorescent magic marker plasmid they contained[9]. Then they were sporulated in SPO++ and either cloNat or G418 for 5-7 days. A random spore prep was used to isolate haploid progeny (https://openwetware.org/wiki/McClean:Random_Spore_Prep), modified to exclude the use of glass beads for spore separation. Cells were plated on selective media, grown for two days, and colony fluorescence was visualized. Green fluorescent colonies and red fluorescent colonies corresponding to MATa and MATα haploid progeny were picked to deep-well 96-well plates and then split into frozen stocks.

#### Preparation of whole-genome sequencing libraries for segregants

Yeast were grown in 1 ml of yeast peptone dextrose in 2-ml deep-well 96-well plates (Thermo Scientific). Plates were sealed with Breathe-Easy gas-permeable membranes (Sigma-Aldrich). Yeasts were grown without shaking for 2 days in a 30 °C incubator. Cell walls were digested with Zymolase, and DNA was extracted using either the 96-well DNeasy Blood & Tissue kits

(Qiagen) for the BYxRM and YPS163xYJM145 segregants, or 96-well E-Z 96 Tissue DNA kit, following the bacterial protocol (Omega) for all other segregants. DNA concentrations were determined using the Quant-iT dsDNA High-Sensitivity DNA quantification kit (Invitrogen) and the Bio-Tek Synergy 2 plate reader. DNA was diluted to 0.22 ng per µl using a Biomek FX liquind handing robot (Beckamn Coulter). For each segregant, 5 µl of 0.22 ng per µl of DNA was added to 4 µl of 5X Nextera HWM buffer (Illumina), 6 µl of water and 5 µl of 1/35 diluted Nextera enzyme. The transposition reaction was performed for 5 min at 55 °C. Directly after the tagmentation reaction and without additional sample purification, Illumina sequencing adaptors and custom indices were added by PCR. 10 µl of tagmented DNA was combined with 0.5 µl each of 10 µM index primers (one of N701-N712 plus one of 96 custom indices), 5 µl of 10X Ex Taq buffer, 0.375 µl Ex Taq polymerase (Takara), 4 µl of 2.5 mM dNTPs and 29.625 µl of water, and amplified with 20 cycles of PCR. Up to 1152-plex libraries were run on a Hiseq 2500 with single end 150 bp reads, except BYxRM[8] and YPS163xYJM145 which were sequenced with 100 bp reads.

**Segregant Genotype Calling**

Fastq files for were demultiplexed using fastq-multx (v1.3.1)[10] and aligned to the SacCer3 version of the reference genome using bwa. Adapter sequences were trimmed from reads and Phred33 quality scores were computed with Trimmomatic (v0.32)[11]. PCR duplicates were removed using Picard and then merged into one CRAM file per cross using Picard. VCF files were generated for each cross using the GATK haplotype caller[4] and genotypes were called at known variant sites between the parental strains. Additional custom provided R code was used to remove regions with strong mapping bias toward the reference genome[12], filter poor quality markers, and remove segregants with too many crossovers, likely corresponding to diploid contaminants. Missing segregant genotype information was imputed using a hidden Markov model (HMM) implemented in R/QTL[13]. Structural variants identified in the parent VCF files were considered missing information in the segregants and the HMM was used to impute genotypes at those sites.

**Phenotyping by endpoint colony growth**

Segregants were arrayed to 384-well liquid plates in duplicate with different plate positions across the duplicates. Segregants were grown in YPD for approximately 48 hours without shaking and then pinned to agar plates using a BM-5 colony arraying robot (S&P Robotics). Plates were incubated for 48 hours and end-point growth was quantified by automated plate imaging using the colony arraying robot. Colony radii were calculated using functions in the EBImage R package[14], and endpoint growth measurements were filtered and normalized for plate effects as described previously[8,15]. In addition, a manual filtering step was used to filter out aberrant colonies arising from technical artefacts, such as from wet spots on the agar plates at the time colonies were pinned. Unless otherwise specified, the average value across replicates was used per segregant for all downstream analyses.

#### Within-cross QTL mapping

QTL were mapped using a forward stepwise regression procedure that controls the FDR[16] for each trait and cross. We tested for linkage at each marker along the genome by calculating $r^2$, where r is the Pearson correlation coefficient between segregant genotypes at the marker and segregant phenotypes. 10,000 permutations of phenotype to strain assignment were performed and this statistic was calculated across the genome for each of the permutations. For each of the permutations, the maximum statistic was recorded to generate an empirical null distribution of the maximum statistic[17]. A p-value was calculated as the probability the observed maximum statistic comes from the empirical null distribution of maximum statistics. If the observed statistic was greater than all of the empirical null statistics the p-value was recorded as 1e-4. The p-value was added to a set of p-values ($p_1, \ldots p_k$), and the entire procedure was repeated (including permutations) with the previously identified marker(s) included as regression covariates. A 'FowardStop', FDR-controlling statistic[16] was calculated as $-\frac{1}{k}\sum_{i=1}^{k}\log(1-p_i)$. We continued to add selected markers to a multiple regression model as long as the 'ForwardStop' statistic was less than or equal to 5%.

We note that we chose to use this procedure rather than procedures we have used in the past[8,12,15] because it is simple, does not require exchangeability of statistics across different traits, gives very similar results as previous methods, and we verified through simulations (not shown) that it controls FDR for forward stepwise model selection under different QTL architectures.

QTL peak positions were re-localized by, one QTL at a time, searching for the marker position that maximizes the likelihood of the data on the chromosome in a model with all other detected QTL peak markers as covariates.

#### Cross-validation procedures to estimate heritability explained by QTL

The amount of additive variance explained by detected QTLs was estimated using cross-validation. For the within-cross analysis, segregants were randomly split into 10 sets. Each set of segregants was left out of the procedure one at a time (held-out set). The within-cross QTL mapping procedure was performed for all the other sets (training set). For the QTL markers detected in this training set and with effects estimated in the training set, the amount of variance explained by the joint model of the set of significant QTL markers was estimated in the held out set. For the joint analysis described below, we performed a similar procedure, splitting the segregants within each cross into 10 sets, leaving one of the sets from each cross out (held-out set) identifying QTL jointly across the other sets (training set) and estimating their effects in each cross (training set) and then estimating the variance explained in the held-out set.

#### Within-cross analysis to estimate additive and pairwise genetic interaction variance

To estimate the fraction of phenotypic variance attributable to additive genetic effects for each cross and trait we fit the model $y=a+e$, where $y$ contains the segregant phenotype values and is standardized to have mean 0 and variance 1. Here, *a* are the additive genetic effects and the residual error is denoted as *e*. The distributions of these effects are assumed to be multivariate normal with mean zero and variance-covariance as follows:

$$a \sim N(0, \sigma_A^2 \boldsymbol{A}) \;\; and \;\; e \sim N(0, \sigma_{EV}^2 \boldsymbol{I})$$

Here, $\boldsymbol{A}$ is the additive relatedness matrix, the fraction of genome shared between pairs of segregants and was calculated as $\boldsymbol{MM'}/n$ where $\boldsymbol{M}$ is the *n* x *m* matrix of standardized marker genotypes, *n* is the number of segregants and *m* is the number of markers.

We also fit an expanded model to estimate the relative contribution of additive vs non-additive (pairwise epistatic) effects. This model was parameterized as:

$$y = \beta X + Zq + Za + Zf + Zg + Zi + Zp + e$$

The distributions of these effects are assumed to be multivariate normal with mean zero and variance-covariance as follows:

$$q \sim N\left(0, \sigma^2_{A_{QTL}} \boldsymbol{A_{QTL}}\right), \qquad a \sim N(0, \sigma_A^2 \boldsymbol{A}), \qquad f \sim N\left(0, \sigma^2_{A_{QTL}*A_{QTL}} \boldsymbol{A_{QTL}} \circ \boldsymbol{A_{QTL}}\right),$$

$$g \sim N\left(0, \sigma^2_{A_{QTL}*A} \boldsymbol{A_{QTL}} \circ \boldsymbol{A}\right),\; i \sim N(0, \sigma^2_{A*A} \boldsymbol{A} \circ \boldsymbol{A}),\; p \sim \mathrm{N}(0, \sigma_R^2 \boldsymbol{I_n}), \text{ and } \mathrm{e} \sim \mathrm{N}(0, \sigma_{EV}^2 \boldsymbol{I_m})$$

where *y* is a vector of length *L* that contains phenotypes for n segregants including replicate measurements such that *L* = n x [number of replicates]. *β* is a vector of estimated fixed effect coefficients. $\boldsymbol{X}$ is a matrix of fixed effects (here *β* is the overall mean, and *X* is a $1_L$ vector of ones unless otherwise specified). $\boldsymbol{Z}$ is an *L* x n incidence matrix that maps *L* total measures to *n* total segregants. In order, the random effect terms correspond to the effects of detected QTL, effects from the whole genome, epistatic interactions between detected QTL, epistatic interactions between additive QTL and the genome, epistatic interactions between all pairs of markers across the genome, and residual repeatability, following very similar methods and syntax as described previously[15]. The mixed model was fit with the regress R package[18] using restricted maximum likelihood estimation (REML). Standard errors of variance component estimates were calculated as the square root of the diagonal of the Fisher information matrix from the iteration at convergence of the Newton-Raphson algorithm. These procedures were used for all other mixed model analyses described below. For the analysis that compared the fraction non-additive to additive variation we calculated $\frac{\sigma^2_{A_{QTL}*A_{QTL}} + \sigma^2_{A_{QTL}*A} + \sigma^2_{A*A}}{\sigma^2_{A_{QTL}} + \sigma^2_A}$.

**Allele-frequency lookup in 1,011 yeast isolate population**

We used bcftools isec[19] to intersect our VCF containing sequence variant information on the 16 parental strains with the 1,011 yeast isolate VCF generated by Peter et al.[20], and vcftools[21] to further filter only biallelic variants. This subset of 259,647 biallelic markers was used for variance components analysis and joint QTL mapping across the panel. Allele frequencies in the larger panel of 1,011 yeast isolates were extracted from the provided VCF[20]. Derived allele frequencies were calculated by using nucmer[22] to perform whole genome alignment between the sacCer3 reference assembly and the CBS432 assembly of *S. paradoxus*. Variants were identified using delta-filter and show-snps commands provided in nucmer. Biallelic variants in our panel were classified as ancient if the variant matches the *S. paradoxus* sequence and recent if not. The unfolded allele frequency was calculated as the frequency of the recent variant. We could determine ancestral status for approximately 80% of the biallelic variants. To improve power for enrichment tests we used derived allele frequency <5% and >95%  as cutoffs when comparing effect sizes and signs of effects between derived and ancestral variants.

**Genotype recoding for joint analyses**

We coded the biallelic markers for which we had allele frequency data from the larger yeast isolate panel as -1 for matching the reference strain, or 1 if not matching the reference. If a variant does not segregate in a particular cross it was coded as 0 in that cross.

**Mixed model analysis with allele-frequency partitioning**

We fit the following mixed model model per trait (jointly across the different crosses):

$$y = \beta \boldsymbol{X} + r + c + e$$

The distributions of these effects are assumed to be multivariate normal with mean zero and variance-covariance as follows:

$$r \sim N\left(0, \sigma_R^2 \boldsymbol{A}_{\boldsymbol{maf<1\%}}\right), \qquad c \sim N\left(0, \sigma_C^2 \boldsymbol{A}_{\boldsymbol{maf>=1\%}}\right), \qquad and\ e \sim N(0, \sigma_{EV}^2 \boldsymbol{I}_{\boldsymbol{m}})$$

where *y* is a vector of length 13,950 that contains phenotypes for segregants concatenated across the different crosses. β is a vector of estimated fixed effects of each cross. **X** is an incidence matrix mapping segregants to crosses. Here, the two relatedness matrices $A_{maf<1\%}$ and $A_{maf>=1\%}$ were calculated separately for all markers with MAF<1% and MAF>=1% respectively in the larger panel of 1,011 yeast isolates. Per marker, the genotype values were scaled to have mean 0 and variance 1, for each of the segregants from crosses in which that marker segregates. After the scaling procedure, for any cross in which the marker is fixed, genotype was coded as 0. Then, with **M** being the n segregants by m markers matrix corresponding to the standardized genotypes for that subset of markers, we calculated the relatedness matrix as a Gower's centered matrix[23–25] $\frac{\boldsymbol{MM\prime}}{\frac{tr(\boldsymbol{MM\prime})}{n}}$ which has the property that the average diagonal coefficient equals 1.

We used the same logic to construct additional covariance matrices when more finely binning variants by allele frequency in the external panel (7 allele-frequency bins model). Bins were chosen to contain approximately equal numbers of variants. We also fit the 7 allele-frequency bins model using only variants that were private to each each parent (variants that only segregate in a pair of crosses). In this last model, the allele-frequency of variants used for the analysis are all approximately the same across the panel. Therefore, this last model does not make the assumption that the variance of variants effects is inversely proportional to their frequencies in the mapping panel[26].

The procedure for fitting these models was the same as described above in the section 'within-cross variance component analysis'.

**Accounting for large effect QTL and polygenic background for all chromosomes except the chromosome of interest for joint QTL mapping**

For each chromosome of interest and for each trait and cross and trait we calculated $y_c = C\mathbf{Q} + a_L + s_c$ where $y_c$ is the vector of trait values for a given trait and cross, $\mathbf{Q}$ is a matrix of QTL genotypes at peak markers from the within-cross mapping described above, with FDR< 5% that are not located on the chromosome of interest, $C$ is a vector of estimated QTL effects from the section 'within-cross QTL mapping', $a_L$ is the additive genetic variance from all chromosomes excluding the chromosome of interest. $a_L$ comes from the REML-based BLUP estimate of the effect all other chromosomes, including the fixed effects of detected QTL on the other. The goal of this step was to obtain the residual trait values $s_c$ that can be used to scan for QTLs on a chromosome of interest and corrects for mapped genetic sources of variation that do not arise from the chromosome of interest[27].

**Joint QTL mapping**

Under the assumption that a causal biallelic variant has a consistent additive effect in all the crosses in which it segregates, we implemented a model to identify such variants jointly across our entire segregant panel[28,29]. This procedure increases statistical power. For example, for variants that are private to one of the 16 parental strains, this procedure will approximately double the sample size for estimating the variant effects. For variants that are shared between multiple parents, sample size for estimating variant effects will further increase.

For each trait and each chromosome, and then for each marker on that chromosome, we calculated a t-statistic as $\frac{r}{\sqrt{\frac{1-r^2}{n-2}}}$. Here r is the Pearson correlation between the recoded segregant genotypes across the panel, and the vector s, which corresponds to the values of $s_c$ described in the previous section concatenated across the different crosses. The number of informative segregants, n, differs for each biallelic variant, and corresponds to the sum of the sample sizes for each cross in which the variant segregates. P-values were calculated that factor in the different number of informative segregants, n, in the calculation of the degrees of freedom using built-in R functions. The -log10(p) was recorded. This statistic was calculated for each marker on the chromosome. 1,000 permutations of phenotype to strain assignment were performed, but these permutations were performed with phenotype values within each cross (we did not permute values between crosses) and this statistic was calculated across the genome for each of the permutations. For each of the permutations, the maximum statistic was recorded to generate an empirical null distribution of the maximum statistic[17]. A new corrected p-value was calculated as the probability the observed maximum statistic comes from the empirical null distribution of maximum statistics. If the observed maximum statistic was greater than all of the empirical null maximum statistics the p-value was recorded as 1e-3.  The p-value was added to a set of p-values ($p_1$, … $p_k$), and the entire procedure was repeated (including permutations) with the previously identified marker(s) included as regression covariates. A 'FowardStop', FDR-controlling statistic[16] was calculated as described above. We continued to add selected markers to a multiple regression model as long as the 'ForwardStop' statistic was less than or equal to 5%.

Allele frequencies of the lead variants were looked up in the 1,011 isolate panel. For each trait and cross, trait values were standardized and variant effects for all lead variants with FDR<5% from the joint QTL mapping procedure that segregate in that cross were estimated using a multiple regression model.

**Statistical fine-mapping to identify causal genes**

We implemented the probabilistic identification of causal snps (PICS) procedure, a Bayesian approach to estimate the probability that a variant is causal. A very thorough description of the method, including details about the logic and implementation, is present in Fahr et al[30]. We aggregated these probabilities within genes to estimate the probability that a gene contains the causal variant. We noted the position of the observed QTL peak (called the 'lead' variant in the GWAS literature), and its effect size for all QTL that explained more than 2% of phenotypic variance from the within-cross mapping (equivalent to 0.1414 SD units). We assumed that the prior probabilities of a variant being causal, or being identified as a lead variant are equal. For this analysis, we only used variants that fall within a 50kb window centered around the detected QTL peak. For each variant within this window, we simulated the observed QTL effect size on the background of noise, 500 times. Here, noise was estimated as the residual error of the within-cross QTL model for that trait and cross. Each of the simulations was generated by a different permutation of the assignment of the residual error to segregant. We then repeated our mapping procedure for the simulated data and calculated the fraction of simulations where the observed QTL peak from our trait mappings was the lead variant given the simulated causal variant. This posterior probability was estimated for each of the variants within the 50kb window, and then normalized so that the sum of all the probabilities in the window is 1. This generated a variant-level probability of causality for each variant within the window for that trait and cross.

Next, we identified overlapping QTL. Overlapping QTL were defined as the QTL coming from neighboring crosses that shared a parent, have 1.5 LOD drop confidence intervals that overlap, and have QTL effect directions that are consistent between the neighboring crosses. For these overlapping QTL we calculated the product of the causality probabilities (described above) for each variant shared between the two crosses (and segregating in both crosses) and then normalized these probabilities so that they sum to 1. To calculate the probability that a gene was causal, we summed these probabilities for all variants that fell within each gene. Here a gene was defined as all variants that fell within the defined open reading frame as well as variants that fell halfway between the start and stop of the adjacent open reading frames. We calculated a FDR by sorting the observed posterior probabilities of causality per gene from highest to lowest, calculating a posterior error probability as 1 minus the posterior probability of causality, and calculating the cumulative mean of these probabilities[31–33].

We note that the causal gene statistic is an estimate of the posterior probability that a gene is causal assuming that one causal variant in the defined window is responsible for generating a signal in two crosses that share a parent strain, that we estimate the effects of causal variants in both crosses without error, and that genotypes are called without error.

**Gene ontology enrichment analyses**

We tested for GO enrichments using the R package topGO[34], using the Fisher test for enrichment ant the ‘classic’ scoring method that does not adjust the enrichments for significance of child GO terms.

### Supplementary Figures

#### Supplementary Figure 1.

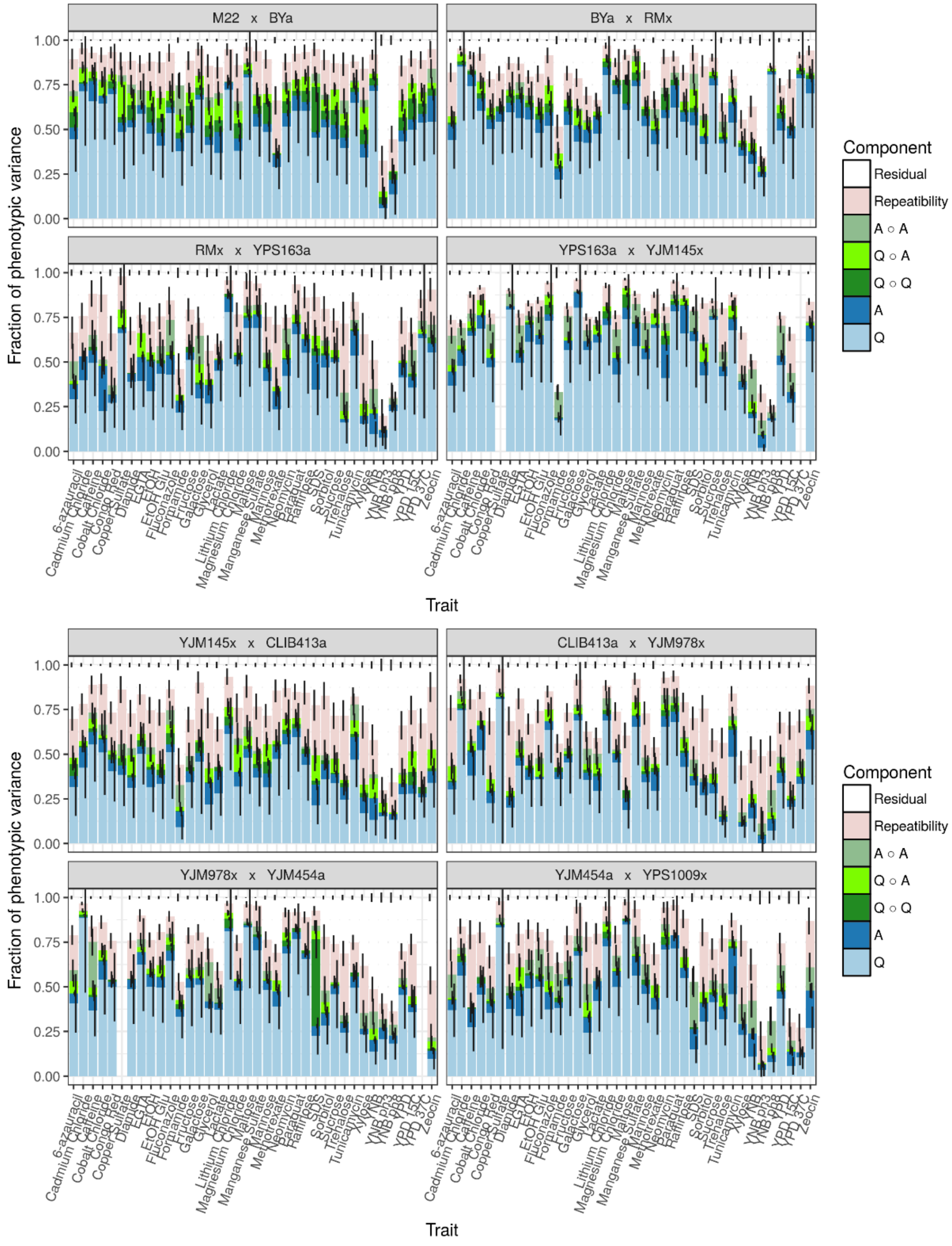

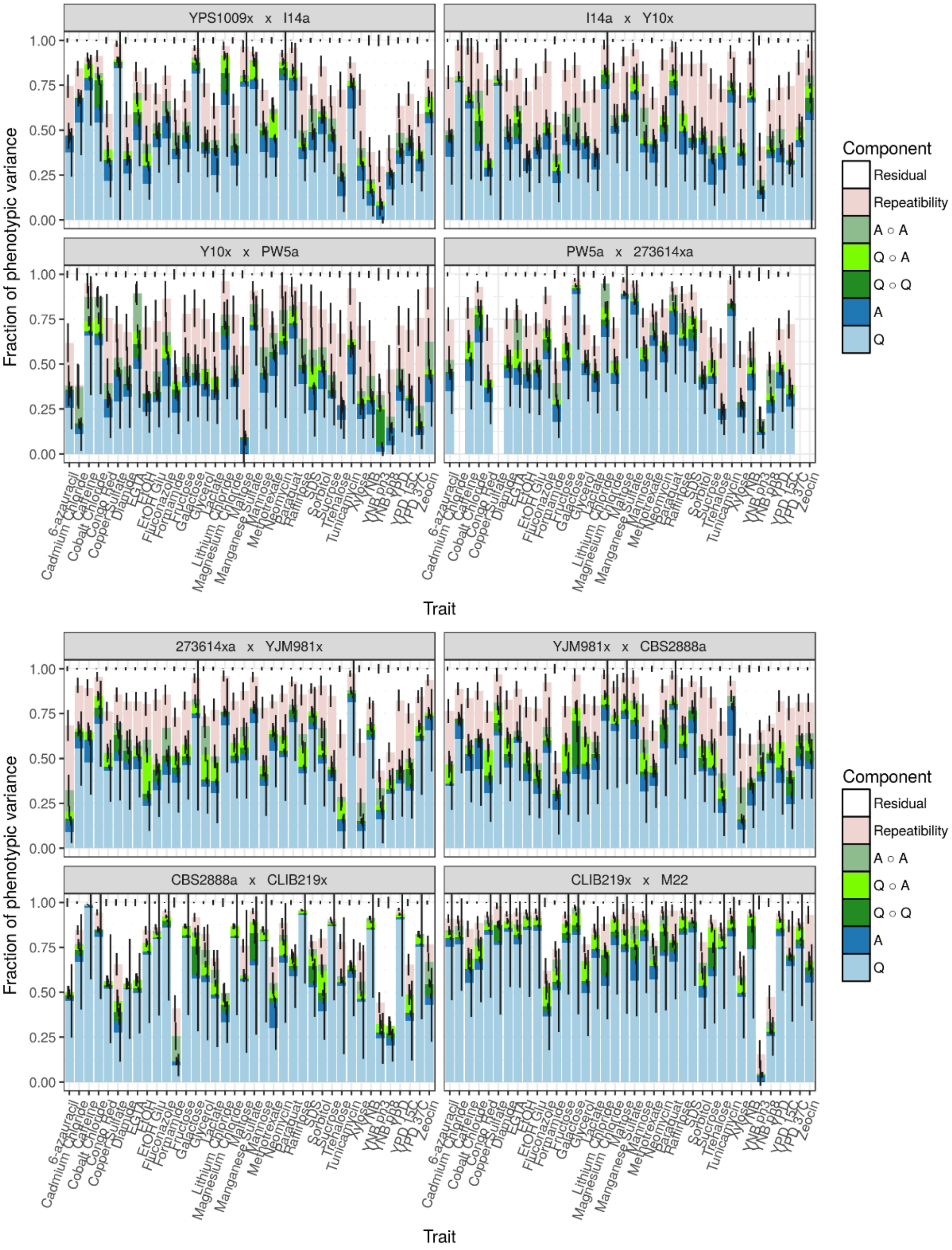


Supplementary Figure 1. For each trait and cross a stacked barplot is shown representing the results from a multiple variance component model. Phenotypic covariance was modeled as the sum of QTL effects (light blue, denoted as Q), additive genome effects (dark blue, denoted as A), interactions between additive QTLs (dark green, denoted as Q∘Q), interactions between additive QTLs and the rest of the genome (light green, denoted as Q∘A ), interactions between all loci in the genome (sea green, denoted as A∘A), residual effect of strain (pink), and residual error (white).

Supplementary Figure 2.

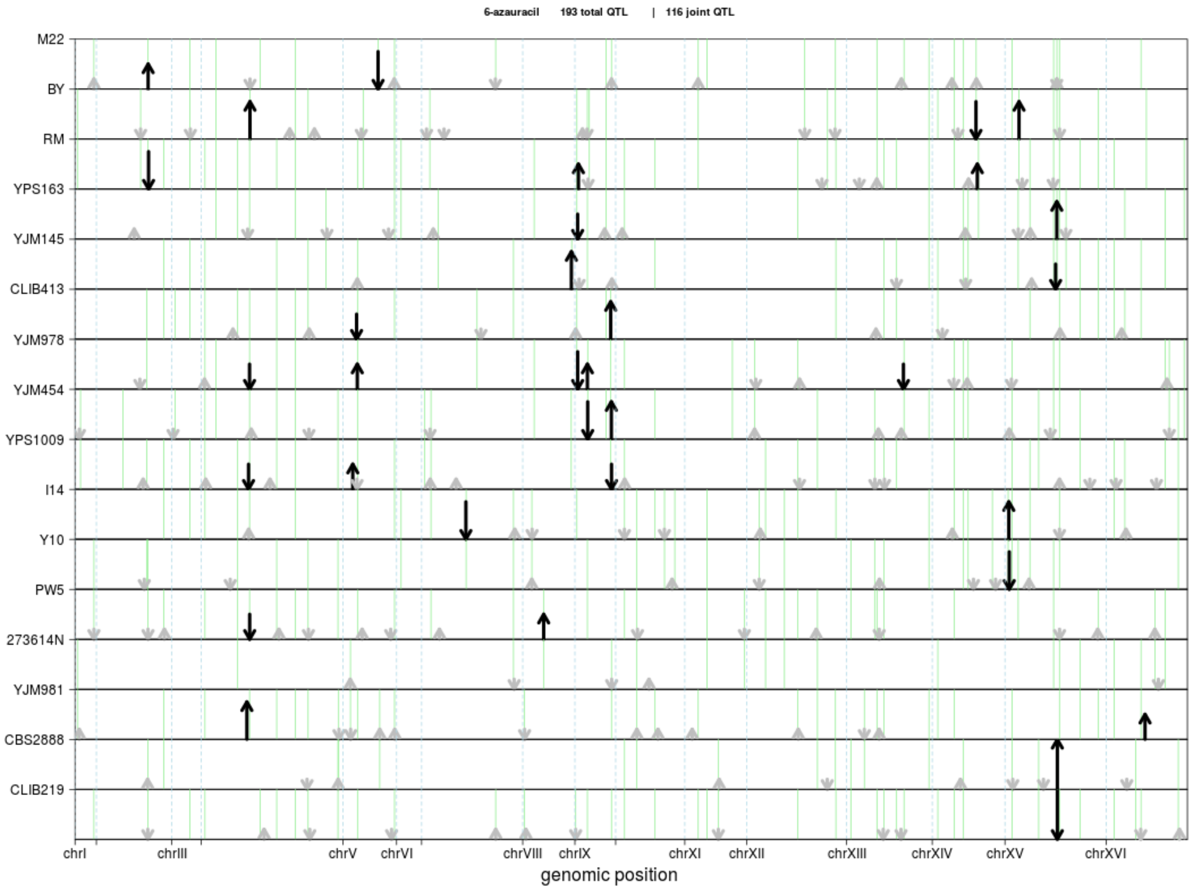

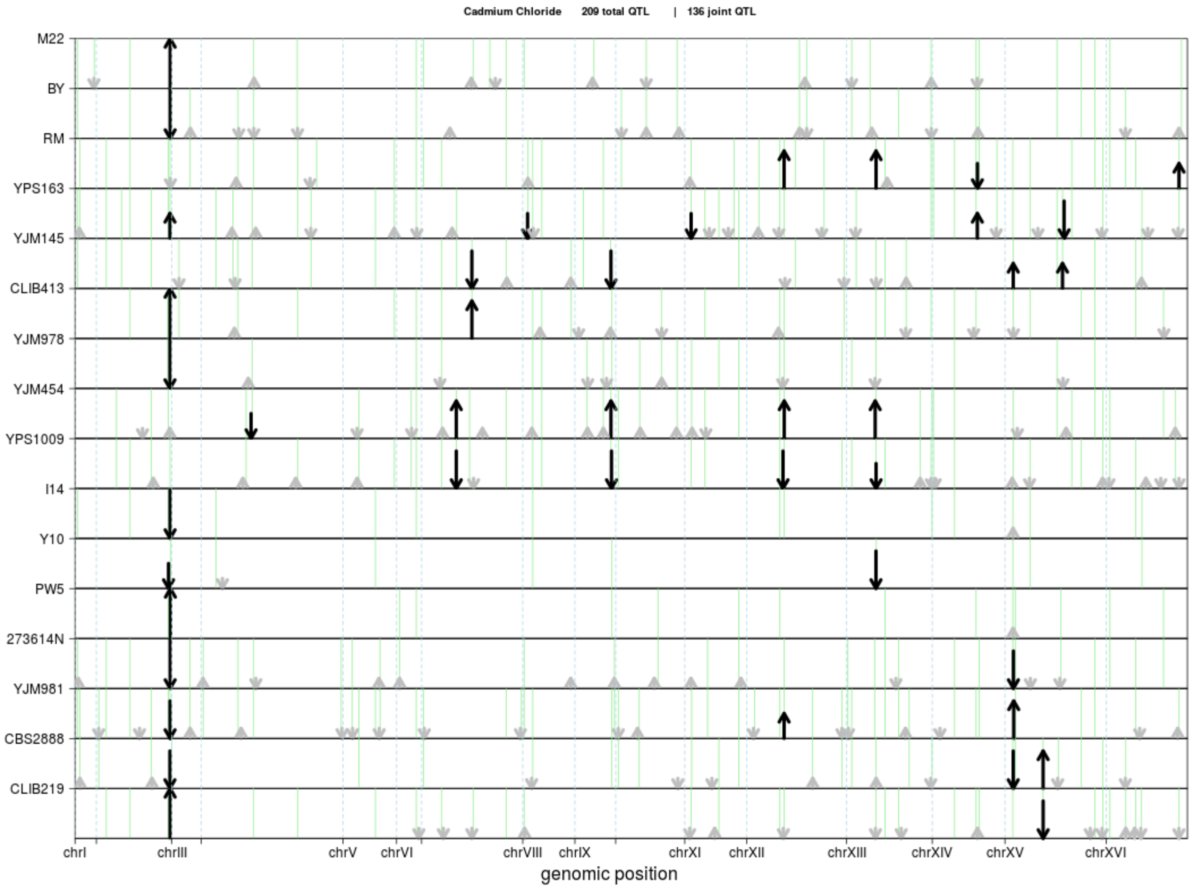

Cadmium Chloride 209 total QTL | 136 joint QTL
M22
BY
RM
YPS163
YJM145
CLIB413
YJM978
YJM454
YPS1009
I14
Y10
PW5
273614N
YJM981
CBS2888
CLIB219
chrI
chrIII
chrV
chrVI
chrVIII
chrIX
chrXI
chrXII
chrXIII
chrXIV
chrXV
chrXVI
genomic position


295

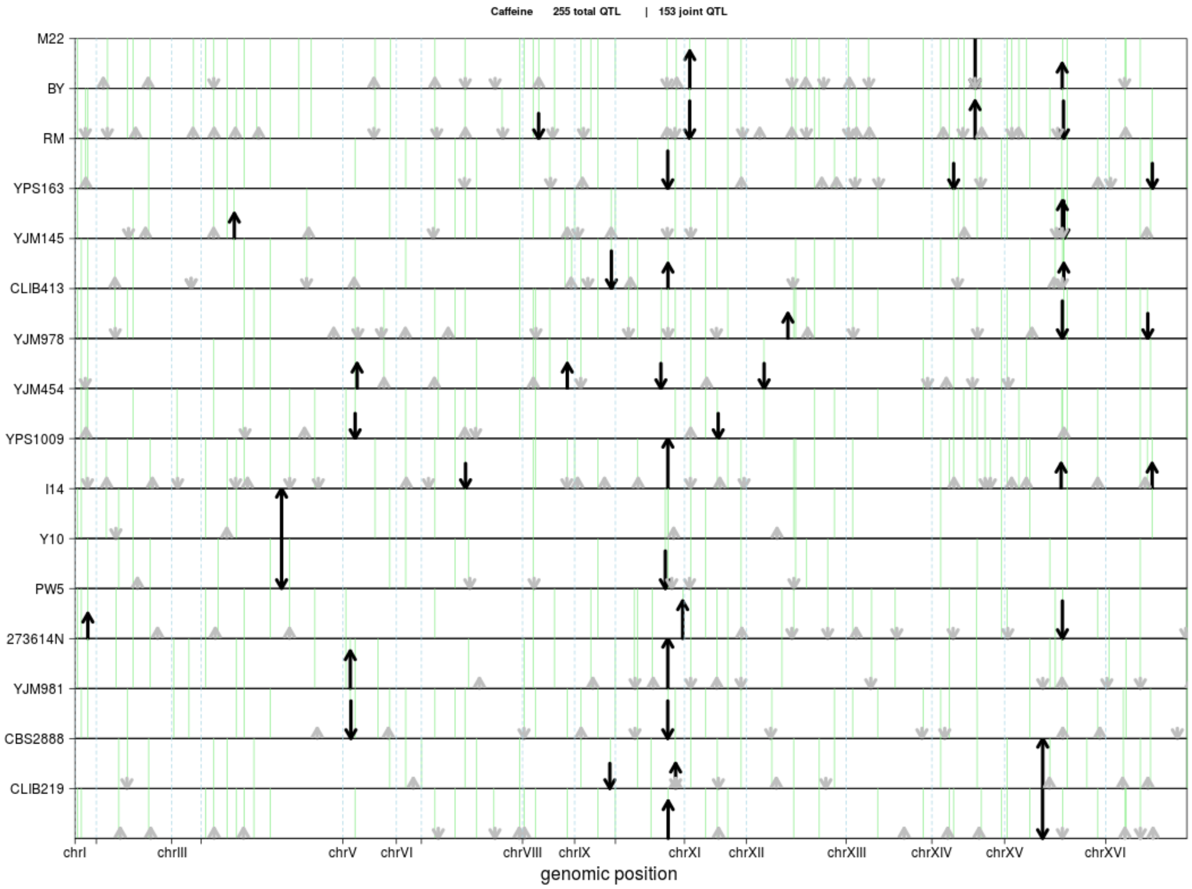

Caffeine 255 total QTL | 153 joint QTL
M22
BY
RM
YPS163
YJM145
CLIB413
YJM978
YJM454
YPS1009
I14
Y10
PW5
273614N
YJM981
CBS2888
CLIB219
chrI
chrIII
chrV
chrVI
chrVIII
chrIX
chrXI
chrXII
chrXIII
chrXIV
chrXV
chrXVI
genomic position


296

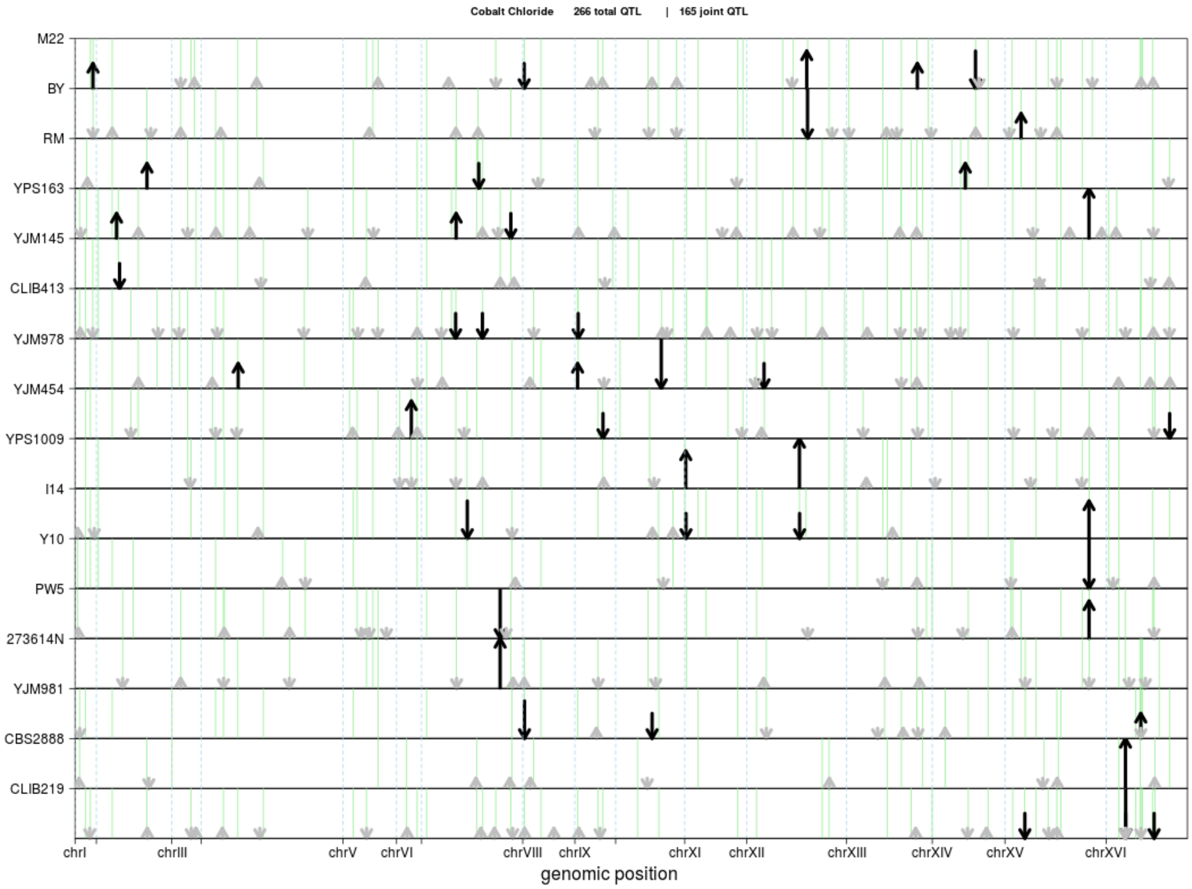

Cobalt Chloride 266 total QTL | 165 joint QTL
M22
BY
RM
YPS163
YJM145
CLIB413
YJM978
YJM454
YPS1009
I14
Y10
PW5
273614N
YJM981
CBS2888
CLIB219
chrI
chrIII
chrV
chrVI
chrVIII
chrIX
chrXI
chrXII
chrXIII
chrXIV
chrXV
chrXVI
genomic position


297

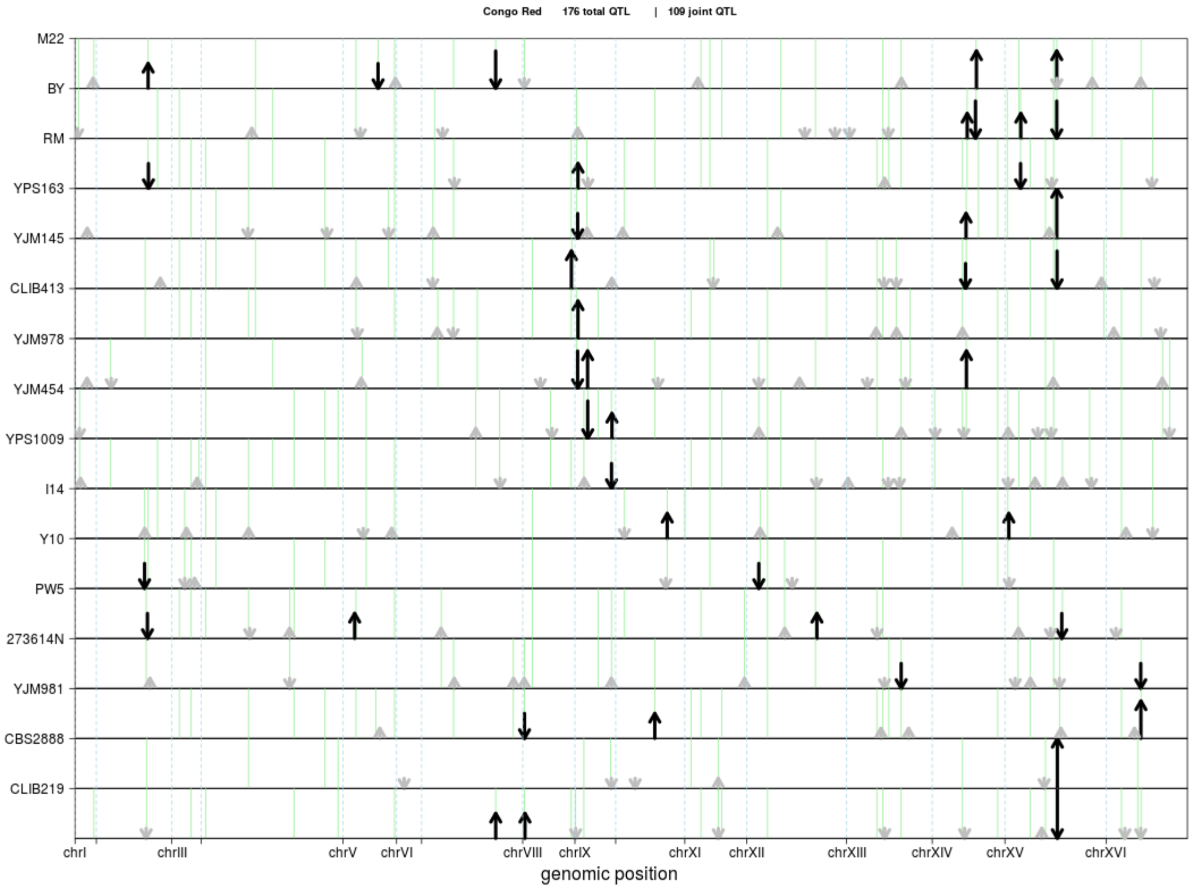

Congo Red 176 total QTL | 109 joint QTL
M22
BY
RM
YPS163
YJM145
CLIB413
YJM978
YJM454
YPS1009
I14
Y10
PW5
273614N
YJM981
CBS2888
CLIB219
chrI
chrIII
chrV
chrVI
chrVIII
chrIX
chrXI
chrXII
chrXIII
chrXIV
chrXV
chrXVI
genomic position


298

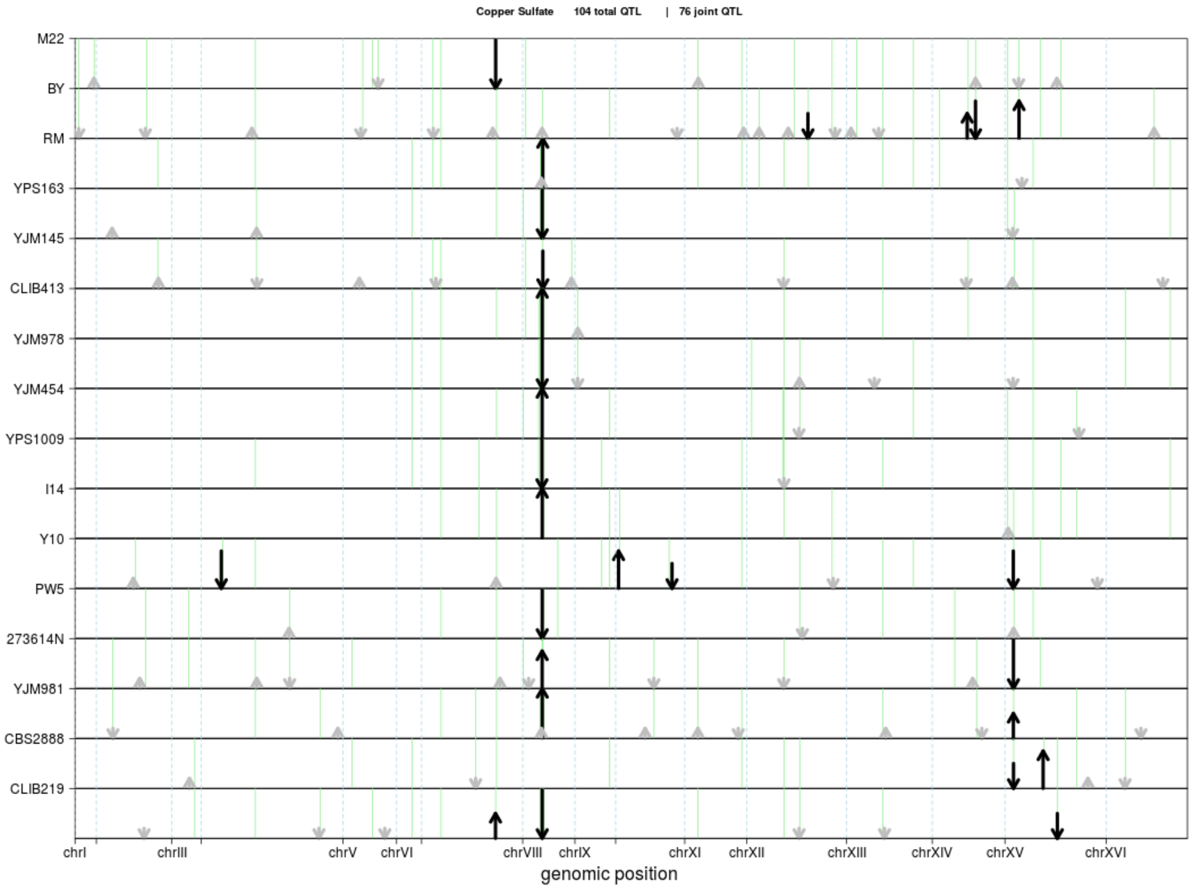

Copper Sulfate 104 total QTL | 76 joint QTL
M22
BY
RM
YPS163
YJM145
CLIB413
YJM978
YJM454
YPS1009
I14
Y10
PW5
273614N
YJM981
CBS2888
CLIB219
chrI
chrIII
chrV
chrVI
chrVIII
chrIX
chrXI
chrXII
chrXIII
chrXIV
chrXV
chrXVI
genomic position


299

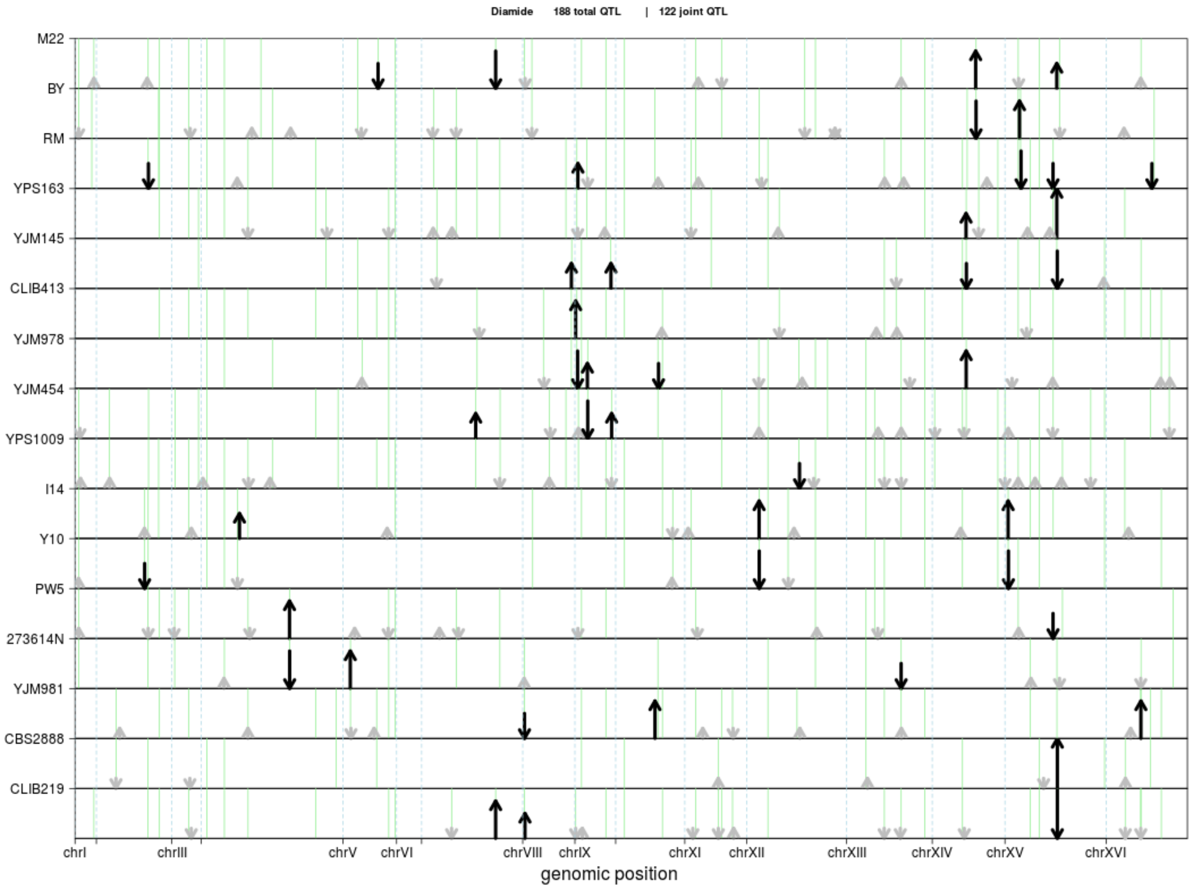

Diamide 188 total QTL | 122 joint QTL
M22
BY
RM
YPS163
YJM145
CLIB413
YJM978
YJM454
YPS1009
I14
Y10
PW5
273614N
YJM981
CBS2888
CLIB219
chrI
chrIII
chrV
chrVI
chrVIII
chrIX
chrXI
chrXII
chrXIII
chrXIV
chrXV
chrXVI
genomic position

301

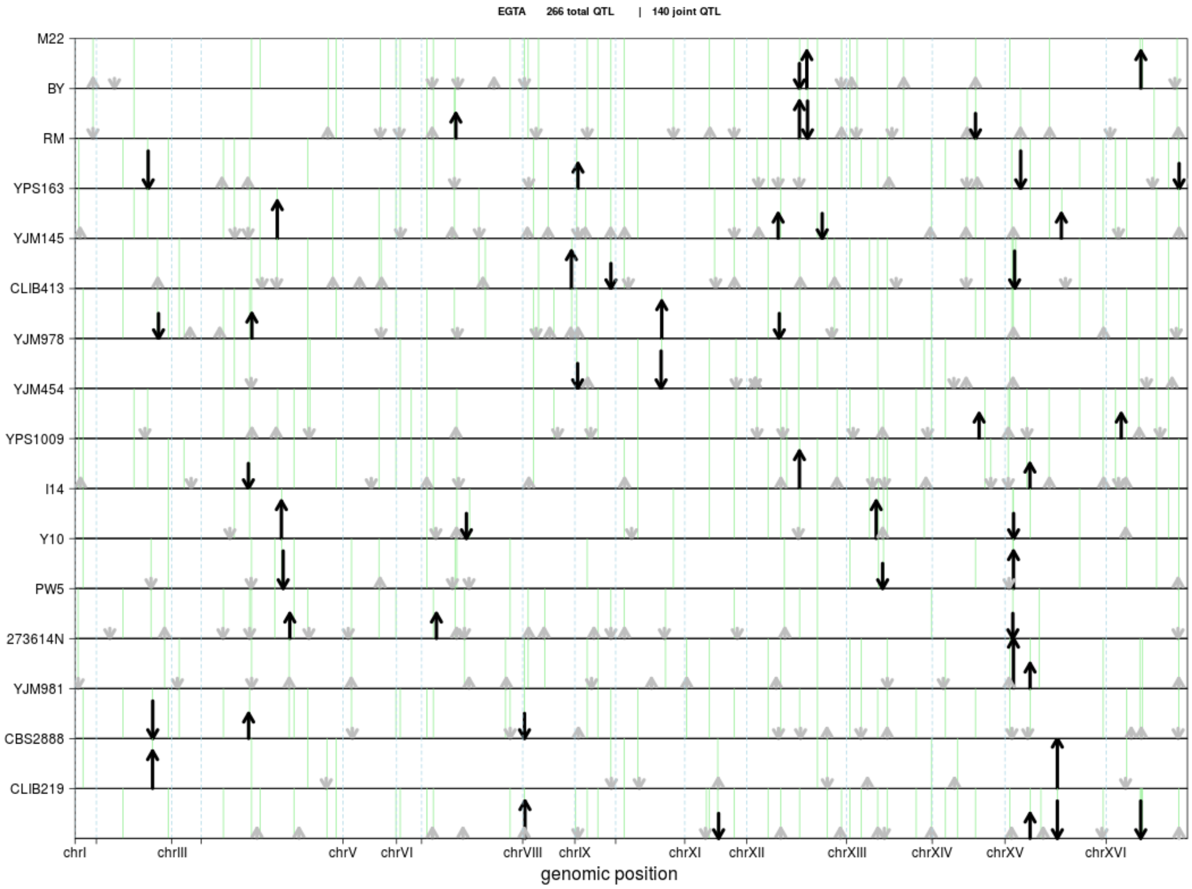
EGTA 266 total QTL | 140 joint QTL
M22
BY
RM
YPS163
YJM145
CLIB413
YJM978
YJM454
YPS1009
I14
Y10
PW5
273614N
YJM981
CBS2888
CLIB219
chrI
chrIII
chrV
chrVI
chrVIII
chrIX
chrXI
chrXII
chrXIII
chrXIV
chrXV
chrXVI
genomic position

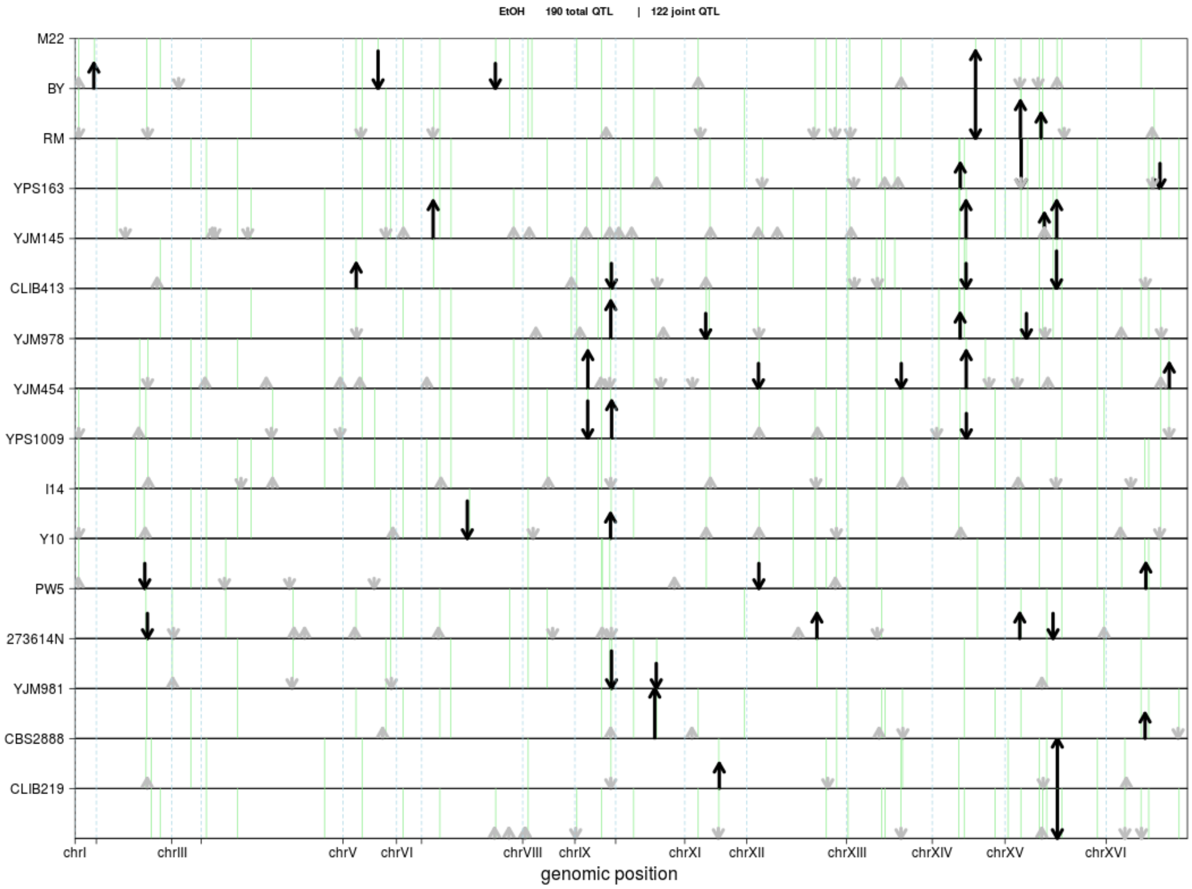
EtOH 190 total QTL | 122 joint QTL
M22
BY
RM
YPS163
YJM145
CLIB413
YJM978
YJM454
YPS1009
I14
Y10
PW5
273614N
YJM981
CBS2888
CLIB219
chrI
chrIII
chrV
chrVI
chrVIII
chrIX
chrXI
chrXII
chrXIII
chrXIV
chrXV
chrXVI
genomic position

302

303

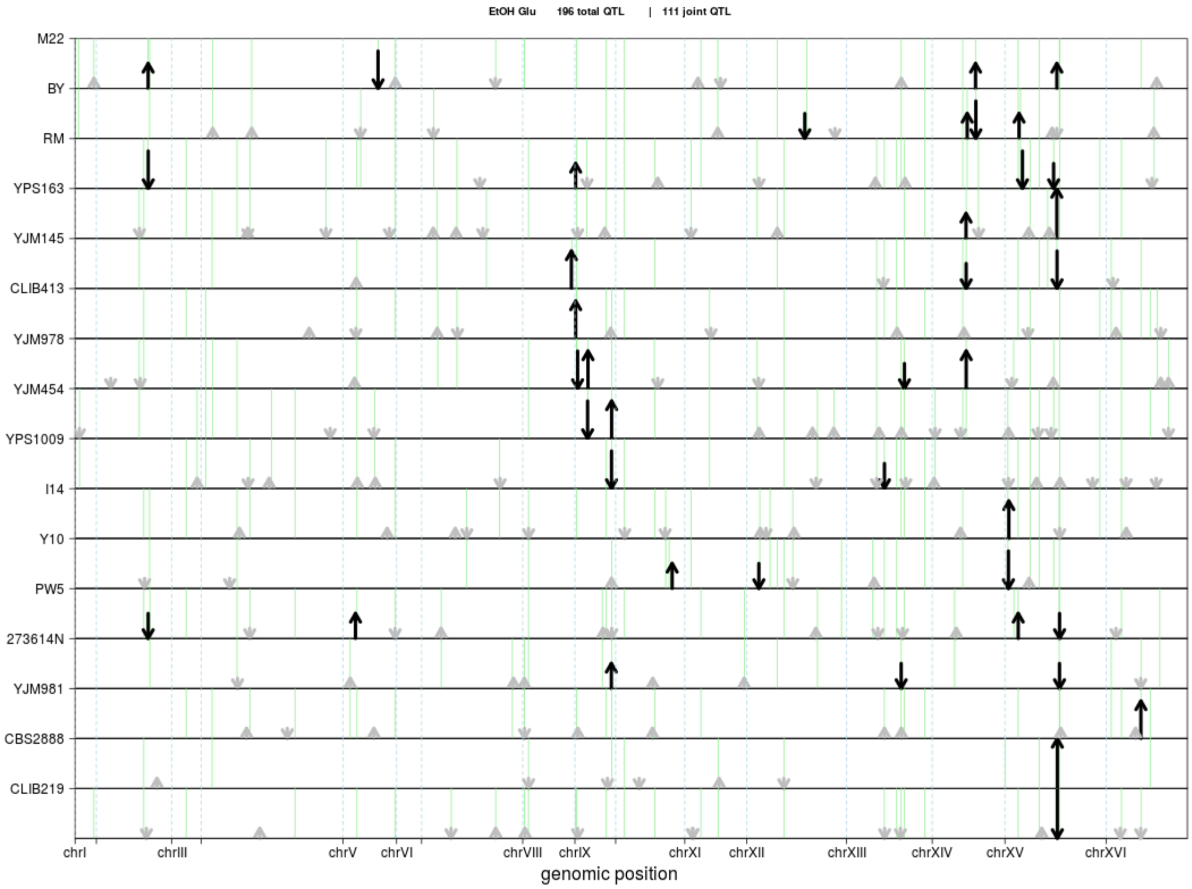
EtOH Glu 196 total QTL | 111 joint QTL
M22
BY
RM
YPS163
YJM145
CLIB413
YJM978
YJM454
YPS1009
I14
Y10
PW5
273614N
YJM981
CBS2888
CLIB219
chrI
chrIII
chrV
chrVI
chrVIII
chrIX
chrXI
chrXII
chrXIII
chrXIV
chrXV
chrXVI
genomic position

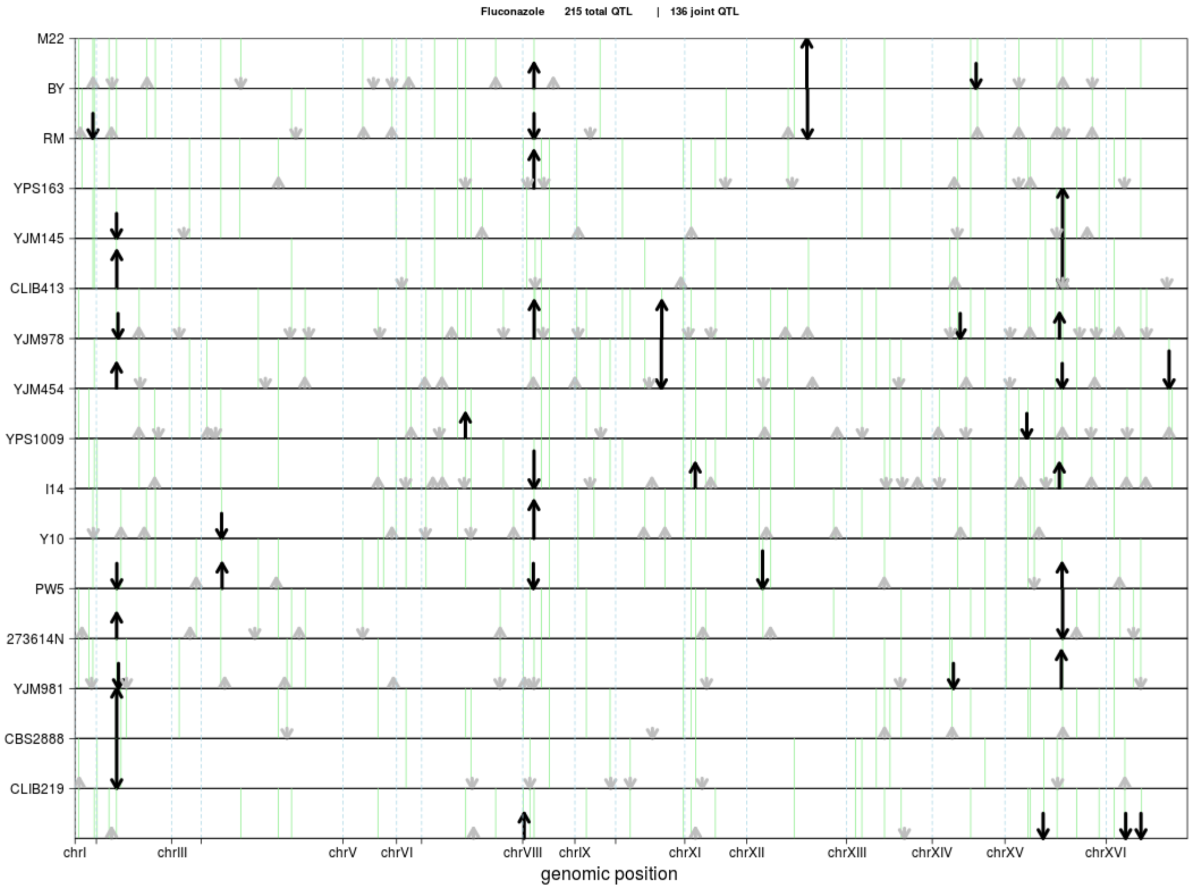
Fluconazole 215 total QTL | 136 joint QTL
M22
BY
RM
YPS163
YJM145
CLIB413
YJM978
YJM454
YPS1009
I14
Y10
PW5
273614N
YJM981
CBS2888
CLIB219
chrI
chrIII
chrV
chrVI
chrVIII
chrIX
chrXI
chrXII
chrXIII
chrXIV
chrXV
chrXVI
genomic position

304

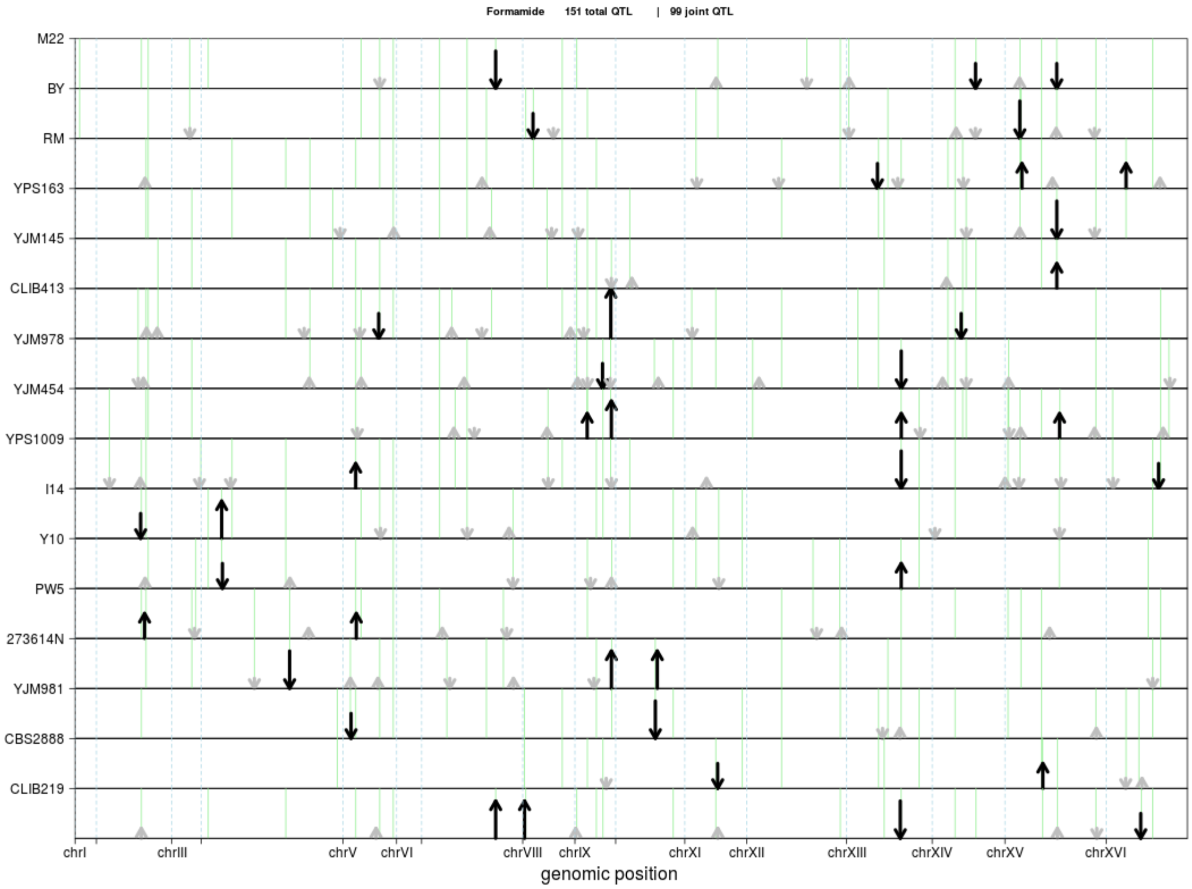
Formamide 151 total QTL | 99 joint QTL
M22
BY
RM
YPS163
YJM145
CLIB413
YJM978
YJM454
YPS1009
I14
Y10
PW5
273614N
YJM981
CBS2888
CLIB219
chrI
chrIII
chrV
chrVI
chrVIII
chrIX
chrXI
chrXII
chrXIII
chrXIV
chrXV
chrXVI
genomic position

305

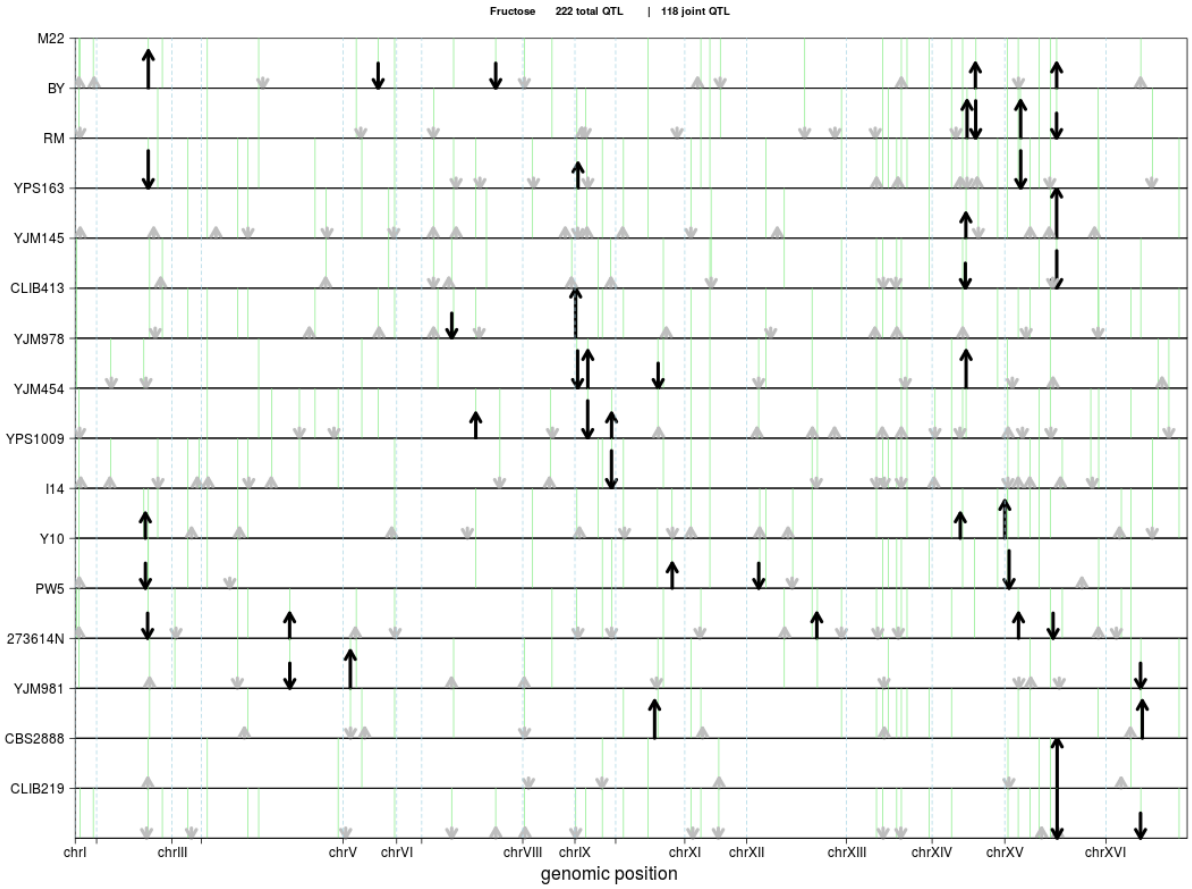

Fructose 222 total QTL | 118 joint QTL
M22
BY
RM
YPS163
YJM145
CLIB413
YJM978
YJM454
YPS1009
I14
Y10
PW5
273614N
YJM981
CBS2888
CLIB219
chrI
chrIII
chrV
chrVI
chrVIII
chrIX
chrXI
chrXII
chrXIII
chrXIV
chrXV
chrXVI
genomic position


306

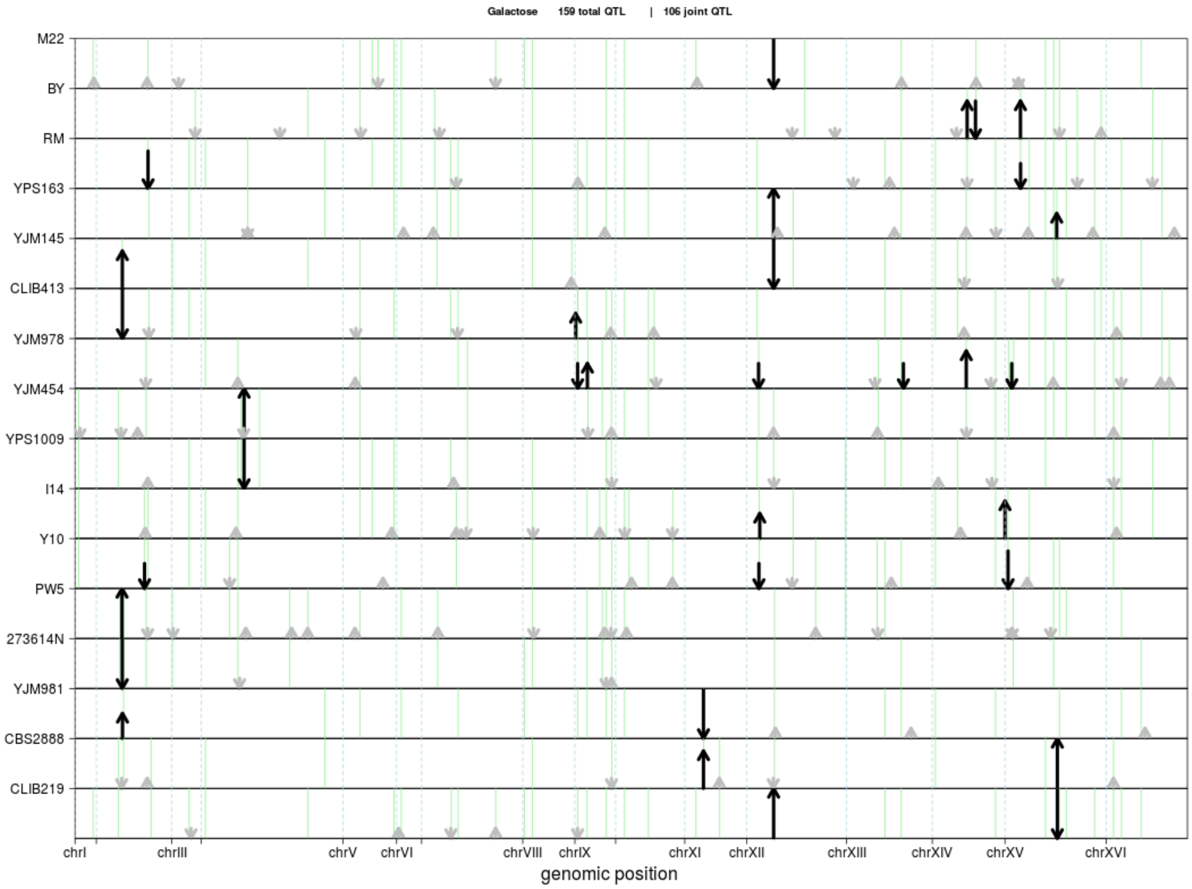

Galactose 159 total QTL | 106 joint QTL
M22
BY
RM
YPS163
YJM145
CLIB413
YJM978
YJM454
YPS1009
I14
Y10
PW5
273614N
YJM981
CBS2888
CLIB219
chrI
chrIII
chrV
chrVI
chrVIII
chrIX
chrXI
chrXII
chrXIII
chrXIV
chrXV
chrXVI
genomic position


307

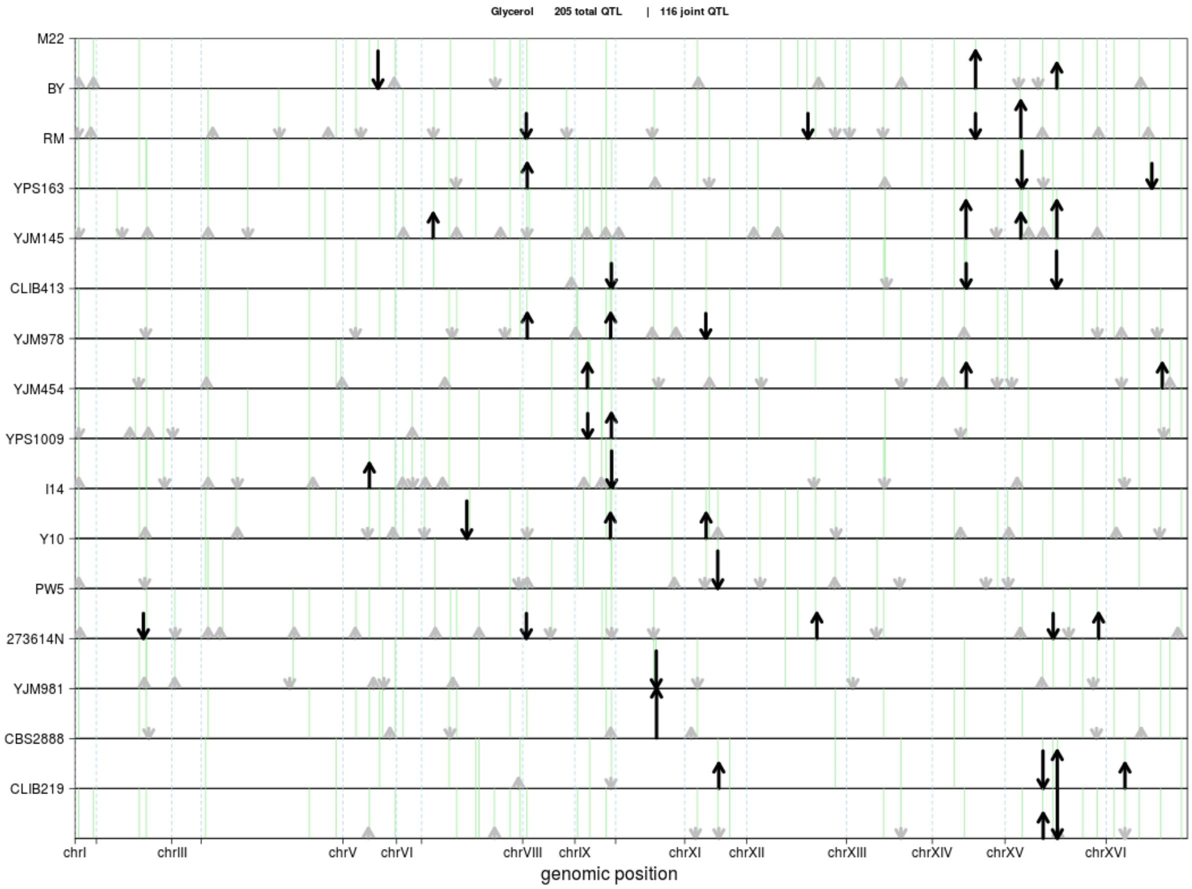

Glycerol 205 total QTL | 116 joint QTL
M22
BY
RM
YPS163
YJM145
CLIB413
YJM978
YJM454
YPS1009
I14
Y10
PW5
273614N
YJM981
CBS2888
CLIB219
chrI
chrIII
chrV
chrVI
chrVIII
chrIX
chrXI
chrXII
chrXIII
chrXIV
chrXV
chrXVI
genomic position


308

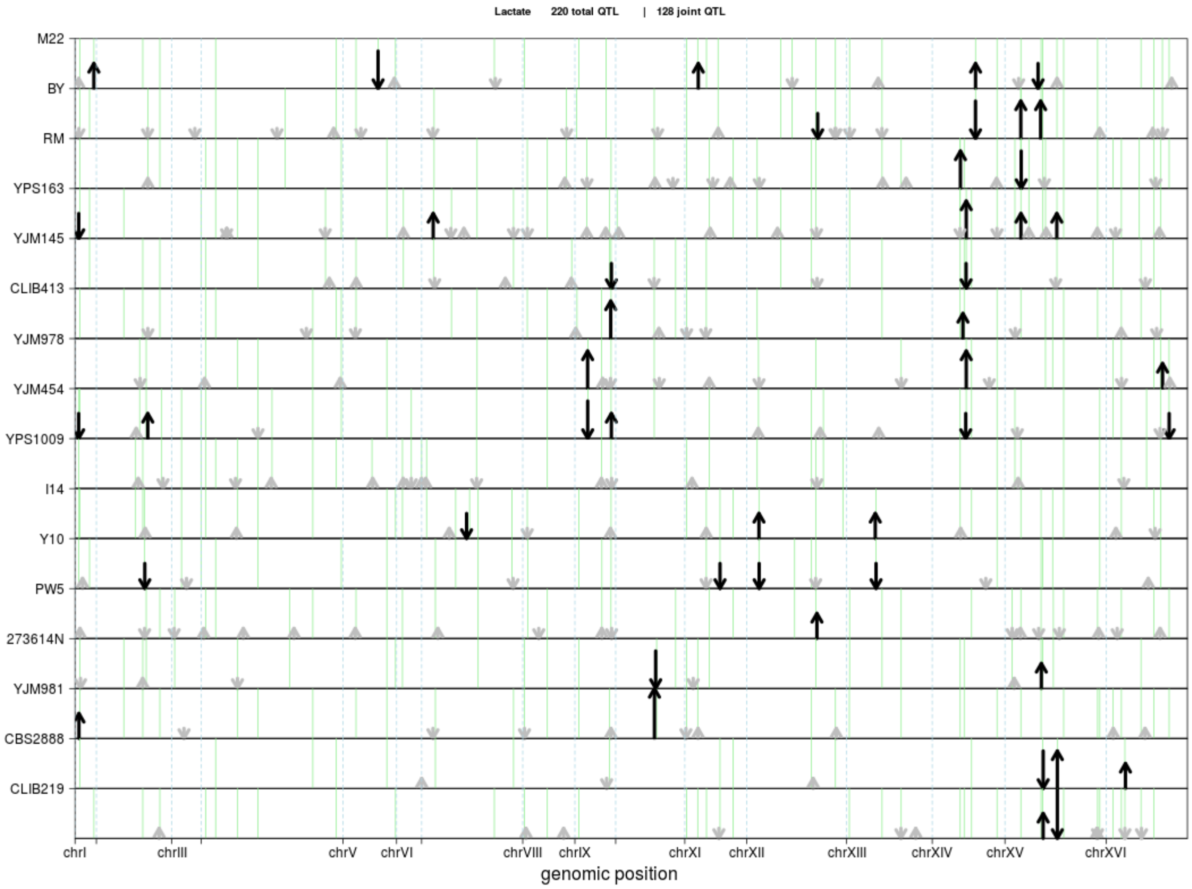

Lactate 220 total QTL | 128 joint QTL
M22
BY
RM
YPS163
YJM145
CLIB413
YJM978
YJM454
YPS1009
I14
Y10
PW5
273614N
YJM981
CBS2888
CLIB219
chrI
chrIII
chrV
chrVI
chrVIII
chrIX
chrXI
chrXII
chrXIII
chrXIV
chrXV
chrXVI
genomic position


309

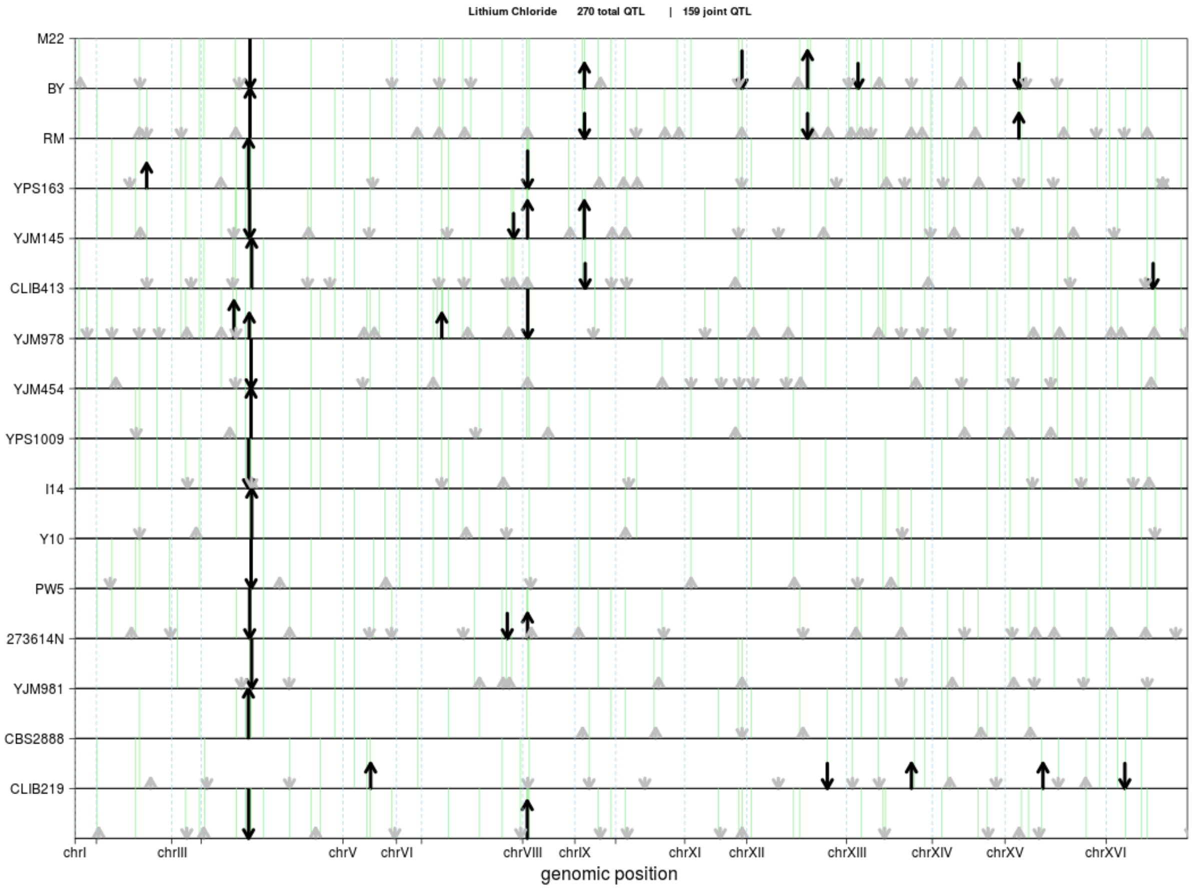

Lithium Chloride 270 total QTL | 159 joint QTL
M22
BY
RM
YPS163
YJM145
CLIB413
YJM978
YJM454
YPS1009
I14
Y10
PW5
273614N
YJM981
CBS2888
CLIB219
chrI
chrIII
chrV
chrVI
chrVIII
chrIX
chrXI
chrXII
chrXIII
chrXIV
chrXV
chrXVI
genomic position


310

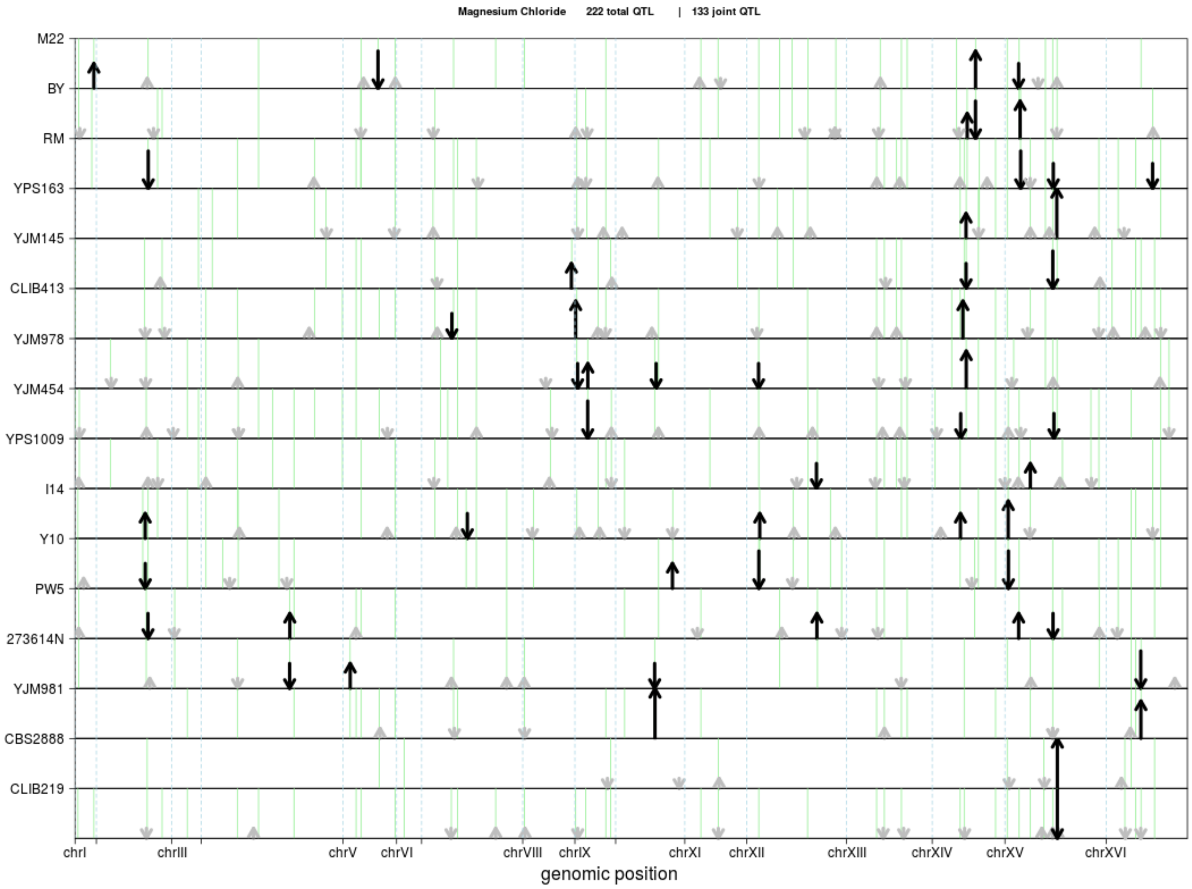

Magnesium Chloride 222 total QTL | 133 joint QTL
M22
BY
RM
YPS163
YJM145
CLIB413
YJM978
YJM454
YPS1009
I14
Y10
PW5
273614N
YJM981
CBS2888
CLIB219
chrI
chrIII
chrV
chrVI
chrVIII
chrIX
chrXI
chrXII
chrXIII
chrXIV
chrXV
chrXVI
genomic position


311

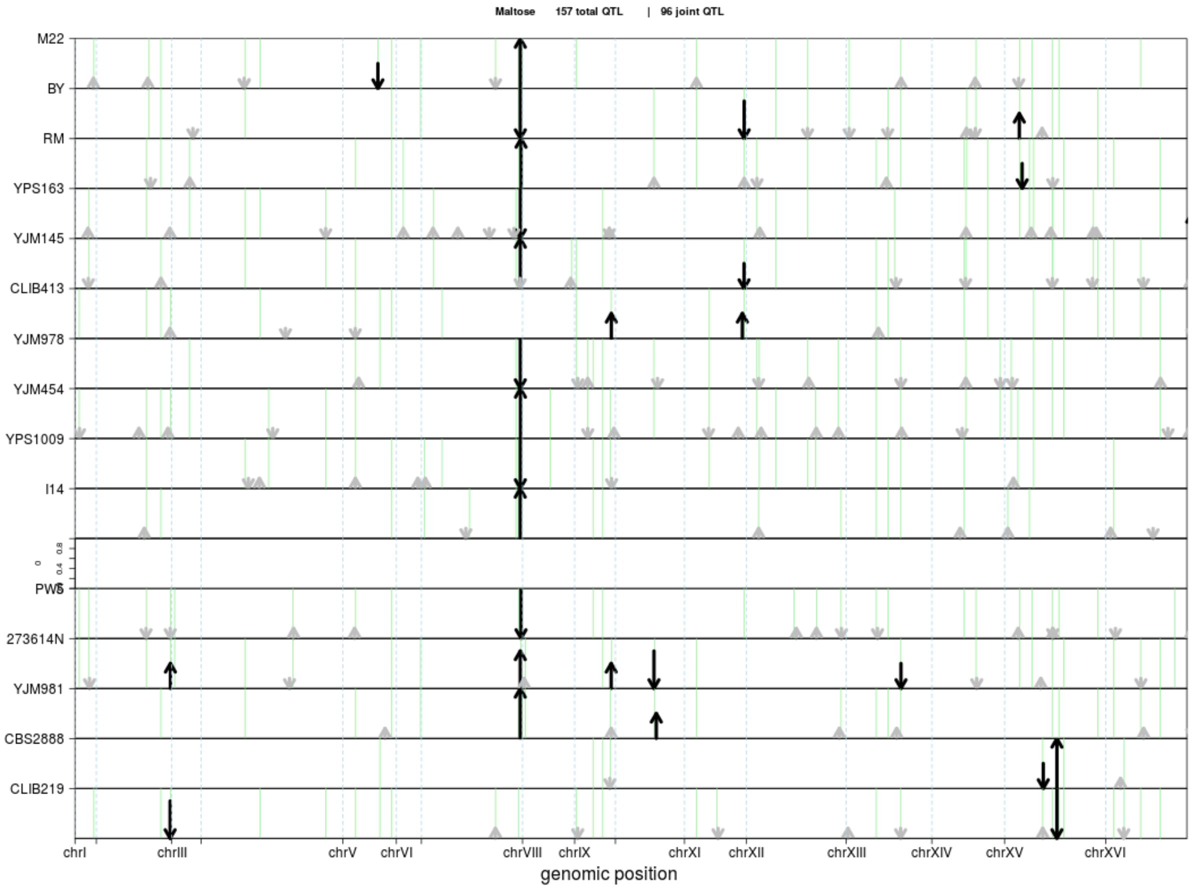

Maltose 157 total QTL | 96 joint QTL
M22
BY
RM
YPS163
YJM145
CLIB413
YJM978
YJM454
YPS1009
I14
PW5
273614N
YJM981
CBS2888
CLIB219
chrI
chrIII
chrV
chrVI
chrVIII
chrIX
chrXI
chrXII
chrXIII
chrXIV
chrXV
chrXVI
genomic position


312

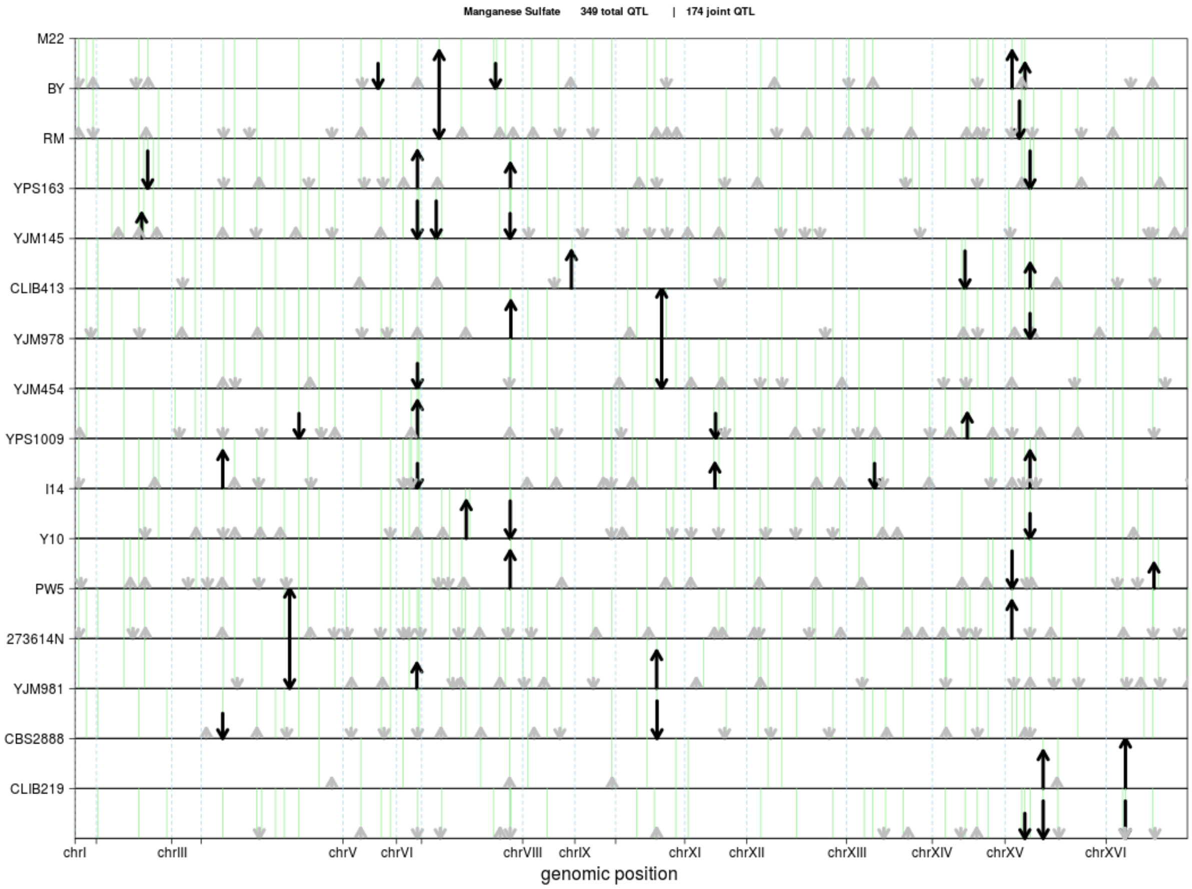
Manganese Sulfate 349 total QTL | 174 joint QTL
M22
BY
RM
YPS163
YJM145
CLIB413
YJM978
YJM454
YPS1009
I14
Y10
PW5
273614N
YJM981
CBS2888
CLIB219
chrI
chrIII
chrV
chrVI
chrVIII
chrIX
chrXI
chrXII
chrXIII
chrXIV
chrXV
chrXVI
genomic position

314

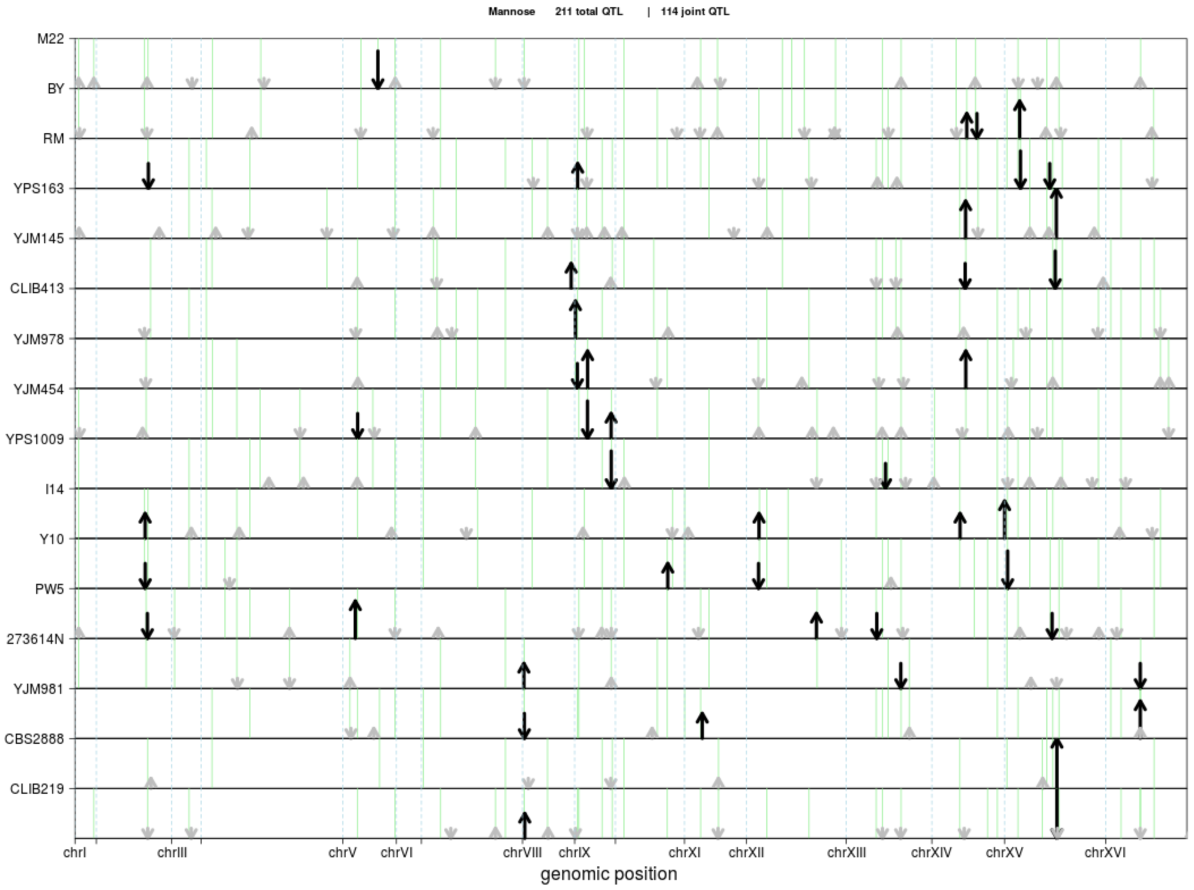
Mannose 211 total QTL | 114 joint QTL
M22
BY
RM
YPS163
YJM145
CLIB413
YJM978
YJM454
YPS1009
I14
Y10
PW5
273614N
YJM981
CBS2888
CLIB219
chrI
chrIII
chrV
chrVI
chrVIII
chrIX
chrXI
chrXII
chrXIII
chrXIV
chrXV
chrXVI
genomic position

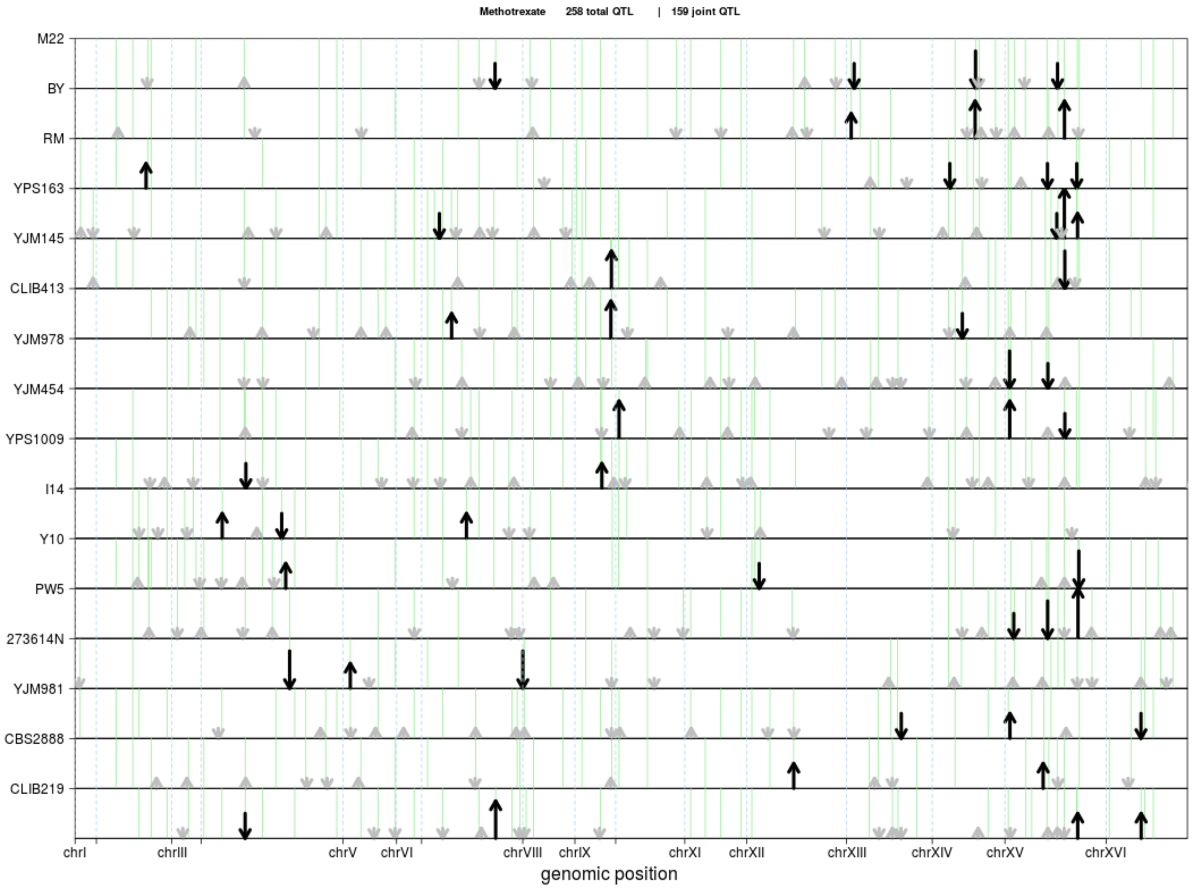

Methotrexate 258 total QTL | 159 joint QTL
M22
BY
RM
YPS163
YJM145
CLIB413
YJM978
YJM454
YPS1009
I14
Y10
PW5
273614N
YJM981
CBS2888
CLIB219
chrI
chrIII
chrV
chrVI
chrVIII
chrIX
chrXI
chrXII
chrXIII
chrXIV
chrXV
chrXVI
genomic position


315

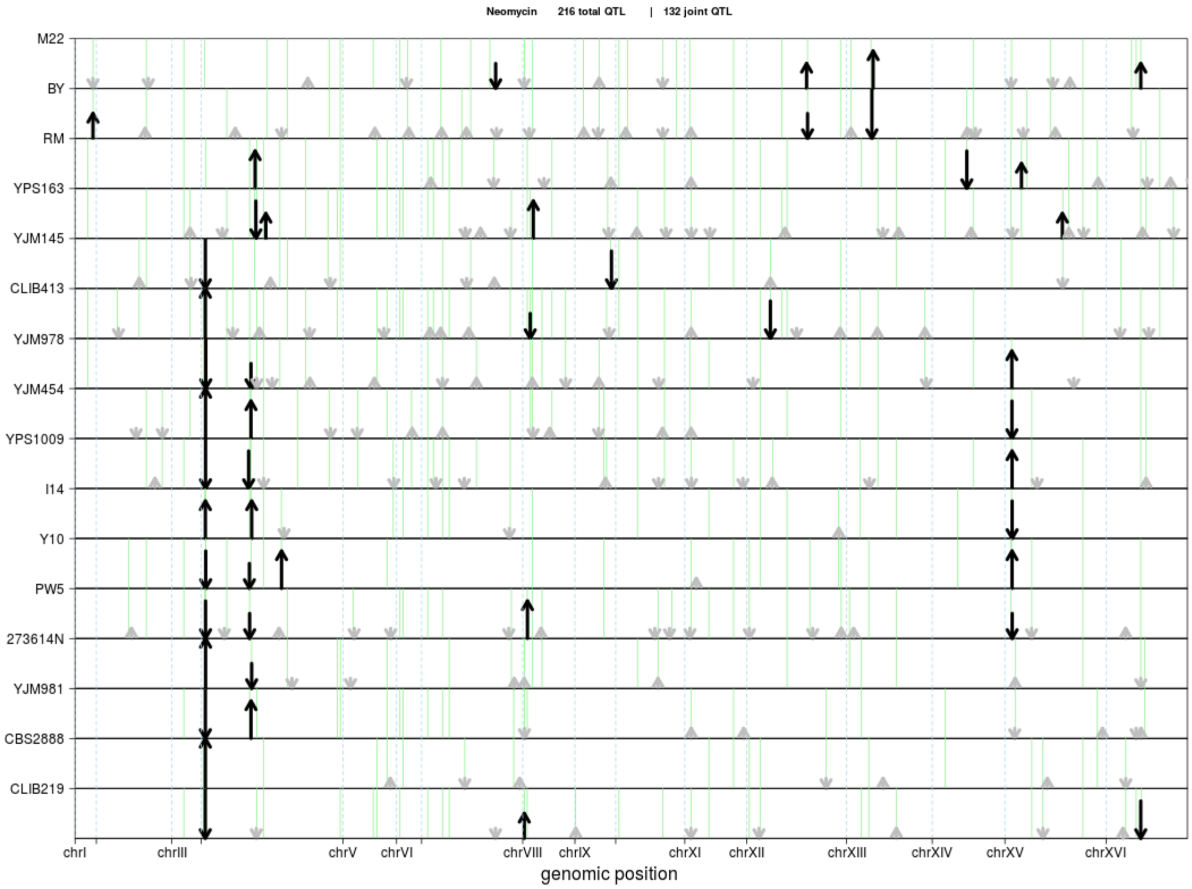

Neomycin 216 total QTL | 132 joint QTL
M22
BY
RM
YPS163
YJM145
CLIB413
YJM978
YJM454
YPS1009
I14
Y10
PW5
273614N
YJM981
CBS2888
CLIB219
chrI
chrIII
chrV
chrVI
chrVIII
chrIX
chrXI
chrXII
chrXIII
chrXIV
chrXV
chrXVI
genomic position

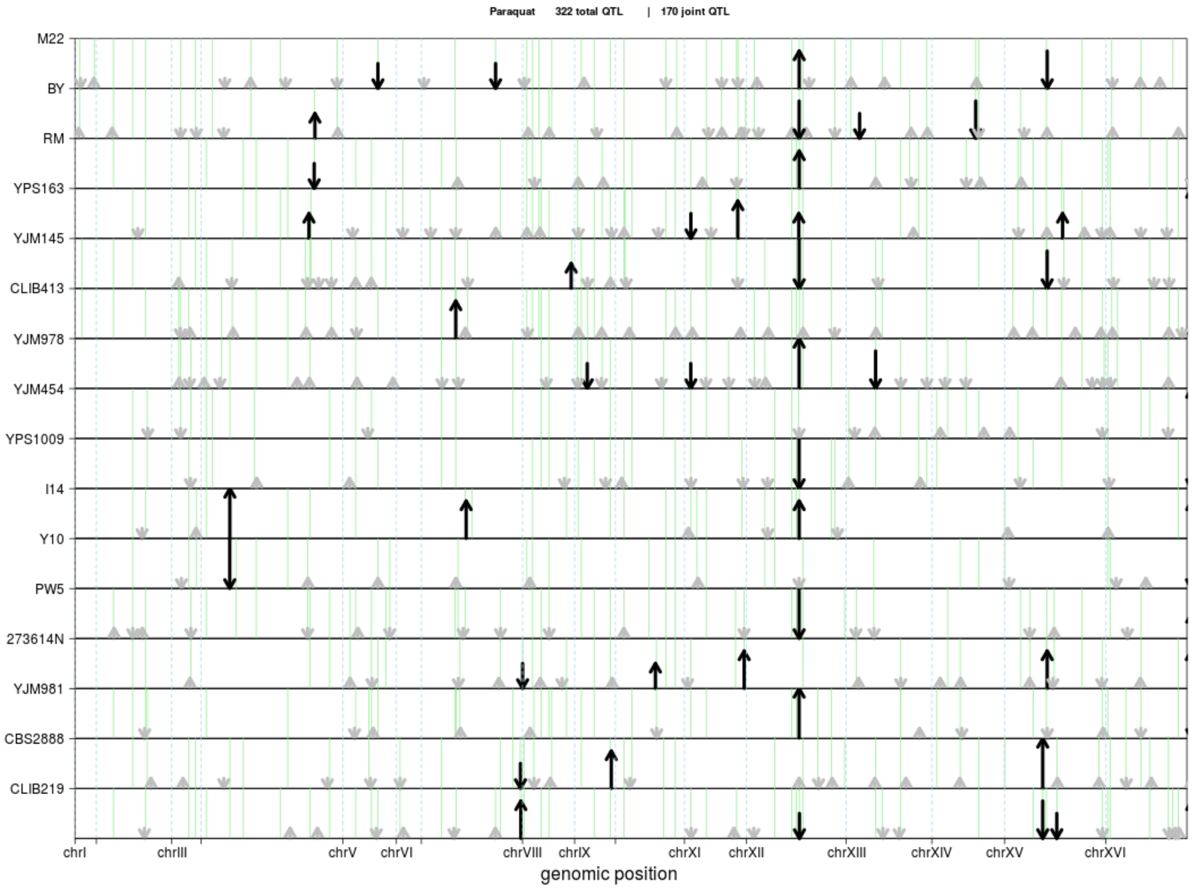

Paraquat 322 total QTL | 170 joint QTL
M22
BY
RM
YPS163
YJM145
CLIB413
YJM978
YJM454
YPS1009
I14
Y10
PW5
273614N
YJM981
CBS2888
CLIB219
chrI
chrIII
chrV
chrVI
chrVIII
chrIX
chrXI
chrXII
chrXIII
chrXIV
chrXV
chrXVI
genomic position


317

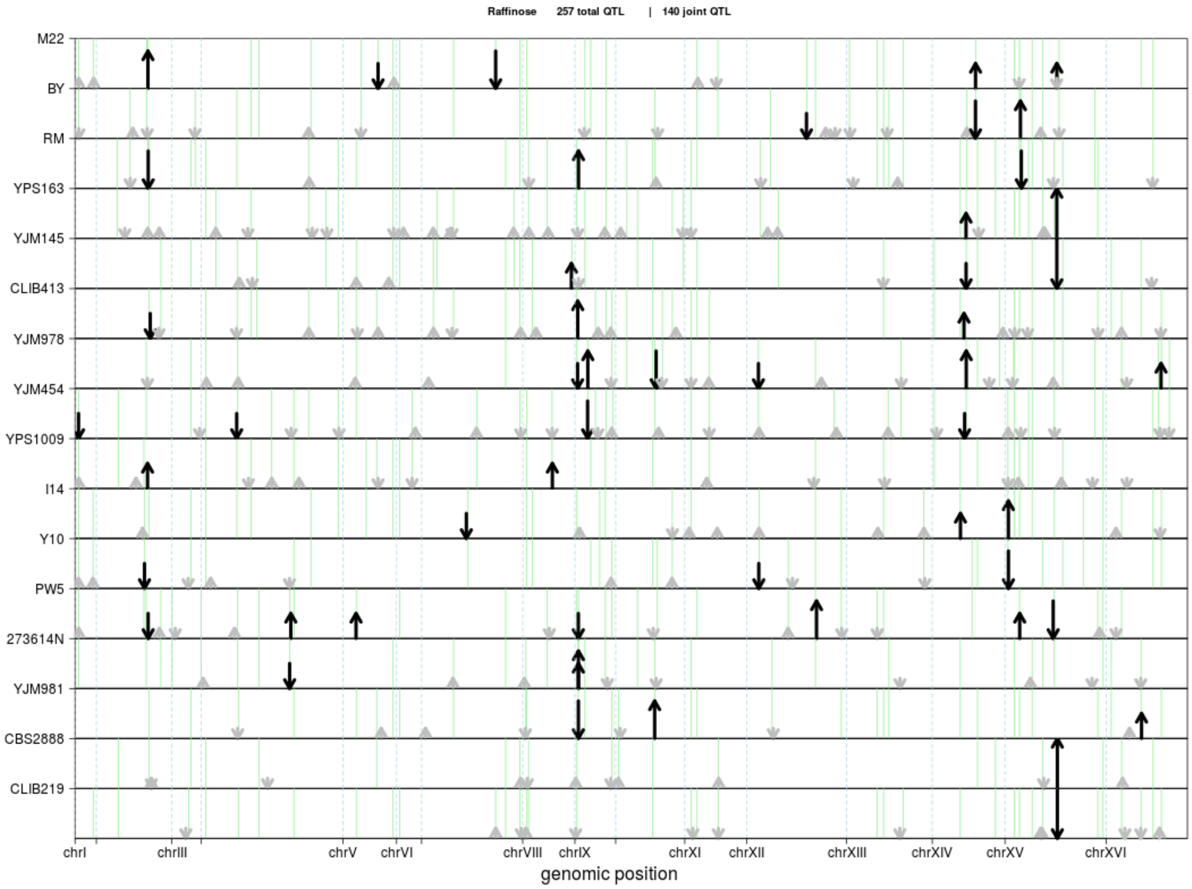
Raffinose 257 total QTL | 140 joint QTL
M22
BY
RM
YPS163
YJM145
CLIB413
YJM978
YJM454
YPS1009
I14
Y10
PW5
273614N
YJM981
CBS2888
CLIB219
chrI
chrIII
chrV
chrVI
chrVIII
chrIX
chrXI
chrXII
chrXIII
chrXIV
chrXV
chrXVI
genomic position

318

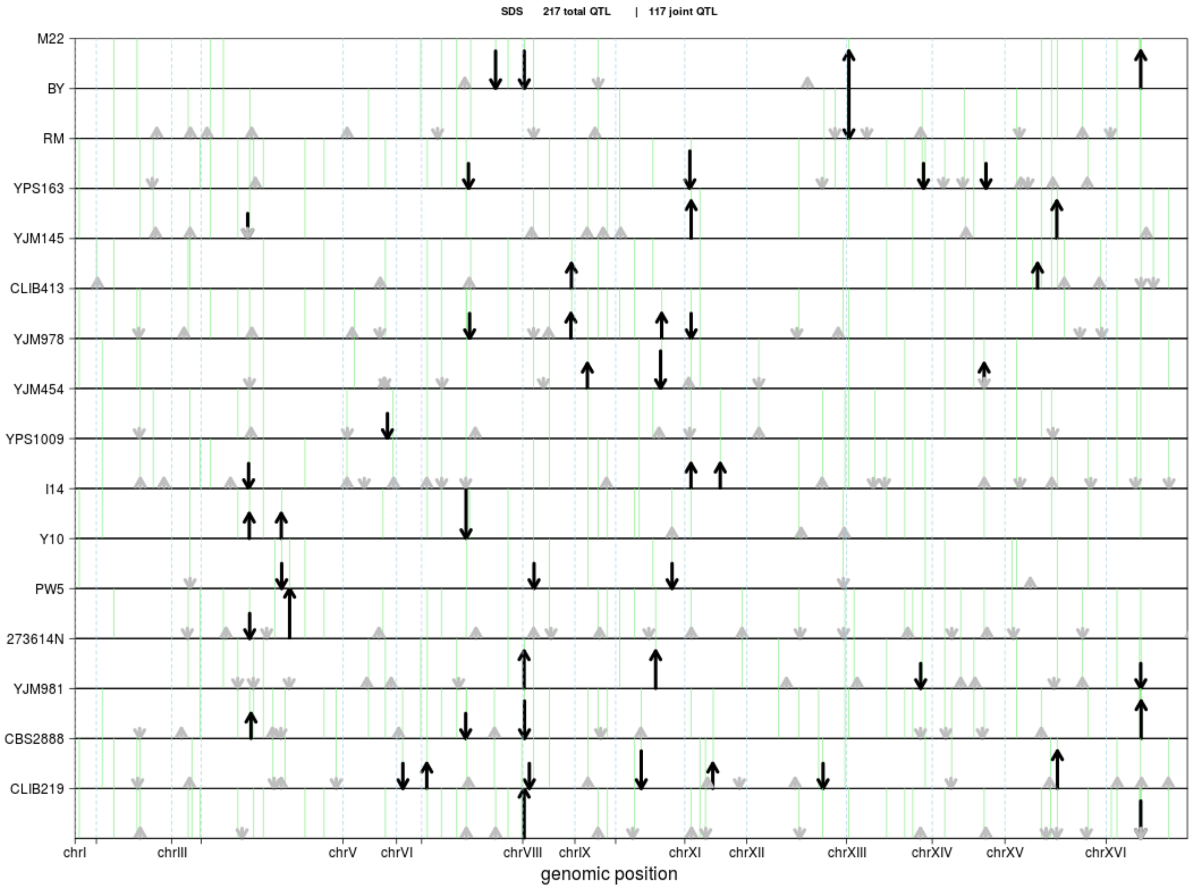
SDS 217 total QTL | 117 joint QTL
M22
BY
RM
YPS163
YJM145
CLIB413
YJM978
YJM454
YPS1009
I14
Y10
PW5
273614N
YJM981
CBS2888
CLIB219
chrI
chrIII
chrV
chrVI
chrVIII
chrIX
chrXI
chrXII
chrXIII
chrXIV
chrXV
chrXVI
genomic position

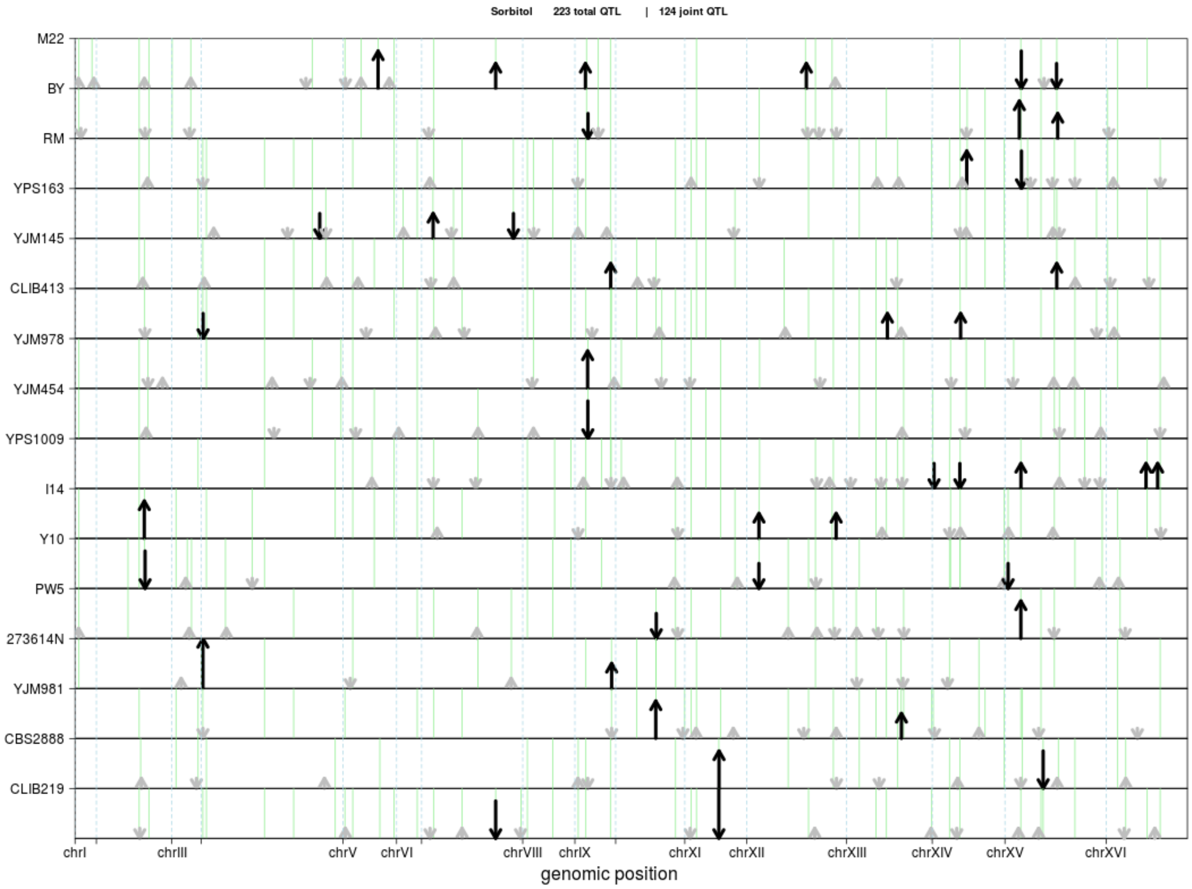

Sorbitol 223 total QTL | 124 joint QTL
M22
BY
RM
YPS163
YJM145
CLIB413
YJM978
YJM454
YPS1009
I14
Y10
PW5
273614N
YJM981
CBS2888
CLIB219
chrI
chrIII
chrV
chrVI
chrVIII
chrIX
chrXI
chrXII
chrXIII
chrXIV
chrXV
chrXVI
genomic position


320

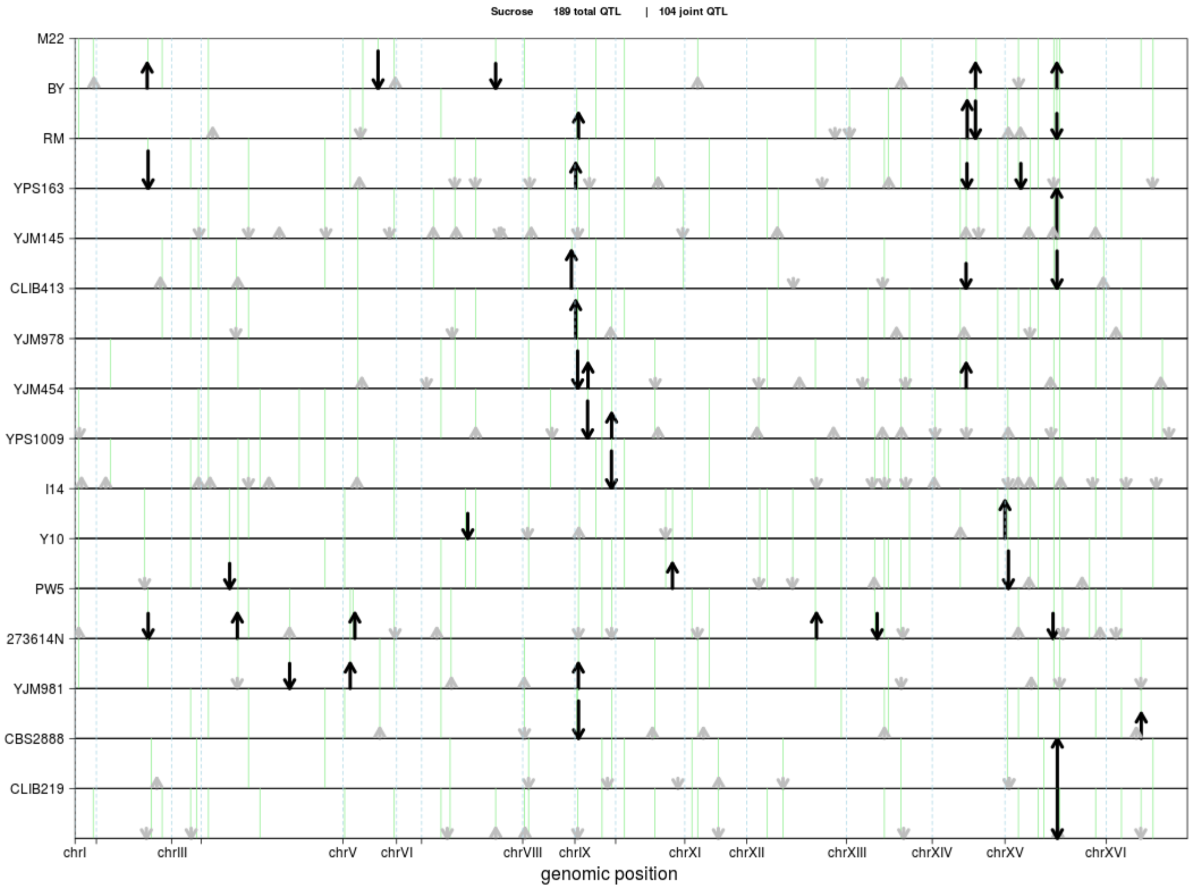

Sucrose 189 total QTL | 104 joint QTL
M22
BY
RM
YPS163
YJM145
CLIB413
YJM978
YJM454
YPS1009
I14
Y10
PW5
273614N
YJM981
CBS2888
CLIB219
chrI
chrIII
chrV
chrVI
chrVIII
chrIX
chrXI
chrXII
chrXIII
chrXIV
chrXV
chrXVI
genomic position

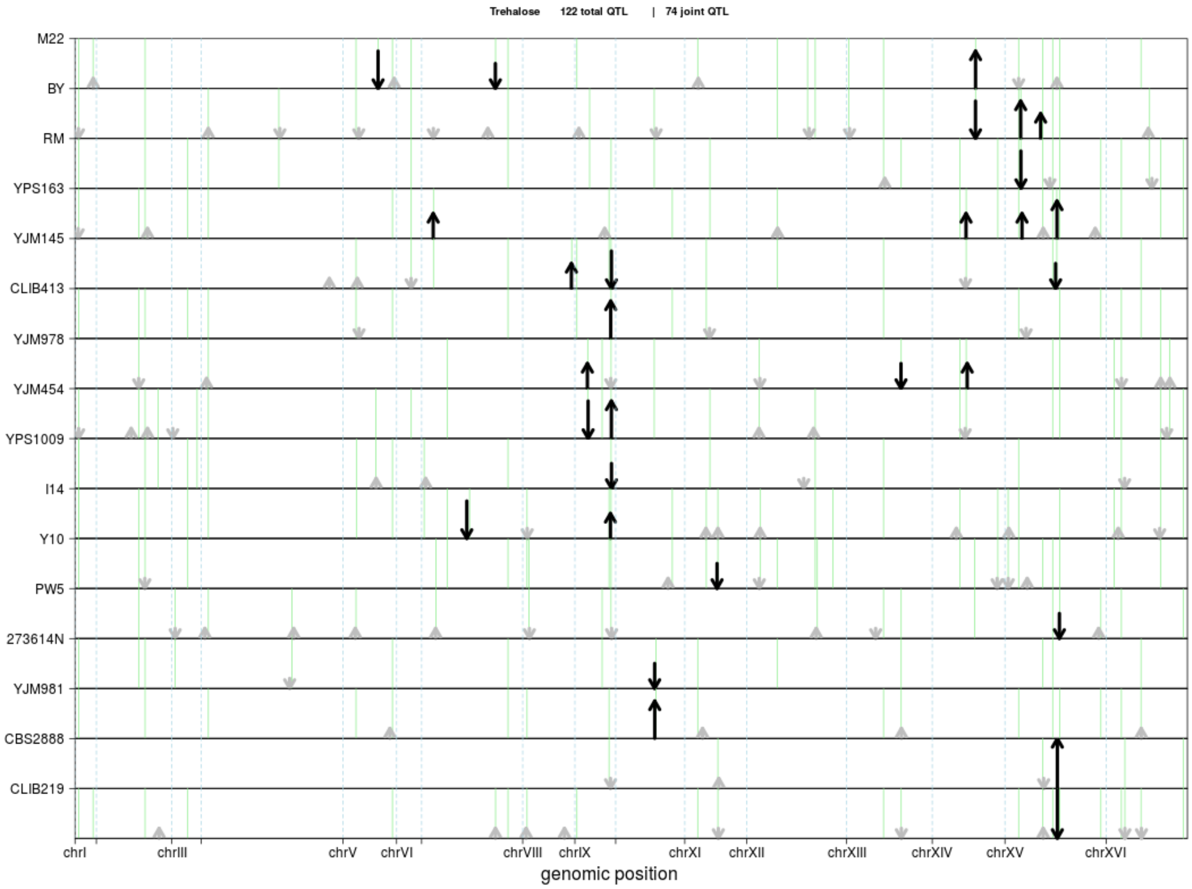
Trehalose 122 total QTL | 74 joint QTL
M22
BY
RM
YPS163
YJM145
CLIB413
YJM978
YJM454
YPS1009
I14
Y10
PW5
273614N
YJM981
CBS2888
CLIB219
chrI
chrIII
chrV
chrVI
chrVIII
chrIX
chrXI
chrXII
chrXIII
chrXIV
chrXV
chrXVI
genomic position

322

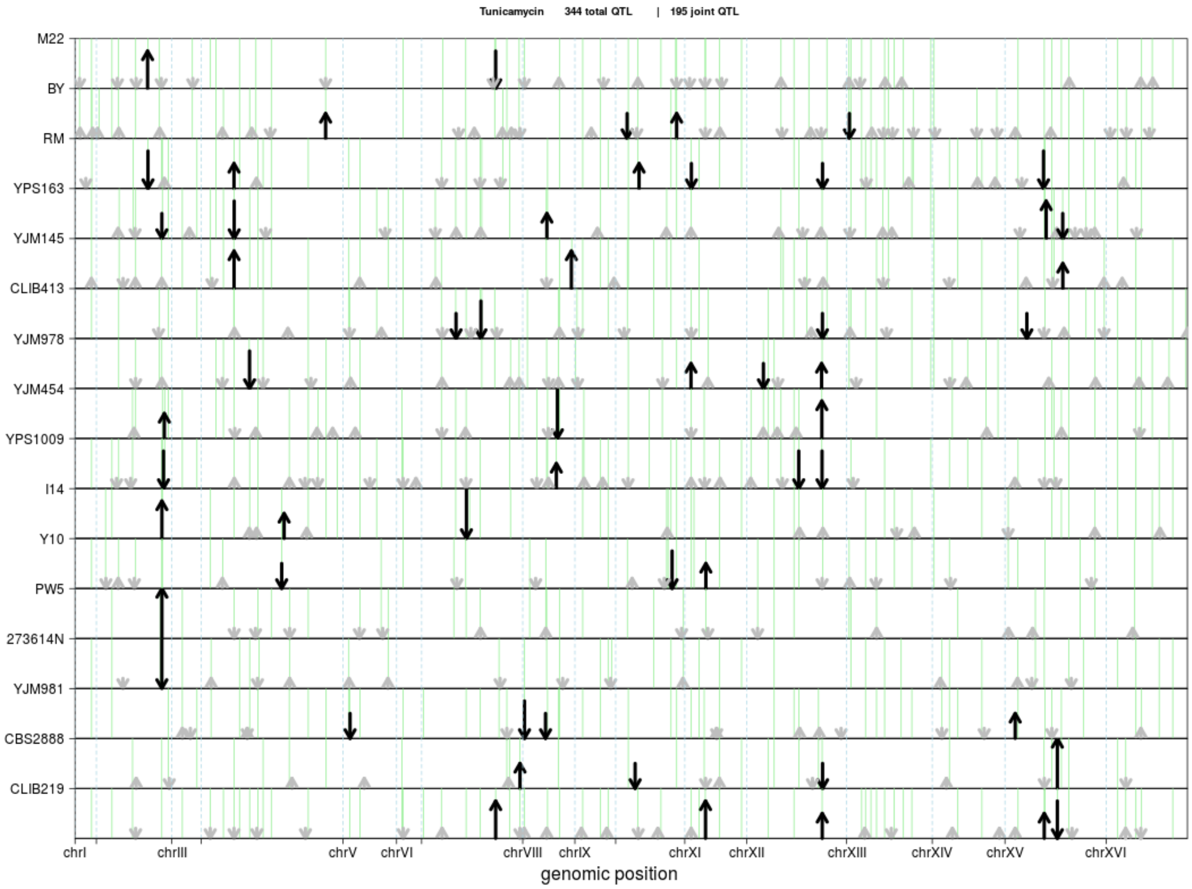

Tunicamycin 344 total QTL | 195 joint QTL
M22
BY
RM
YPS163
YJM145
CLIB413
YJM978
YJM454
YPS1009
I14
Y10
PW5
273614N
YJM981
CBS2888
CLIB219
chrI
chrIII
chrV
chrVI
chrVIII
chrIX
chrXI
chrXII
chrXIII
chrXIV
chrXV
chrXVI
genomic position


323

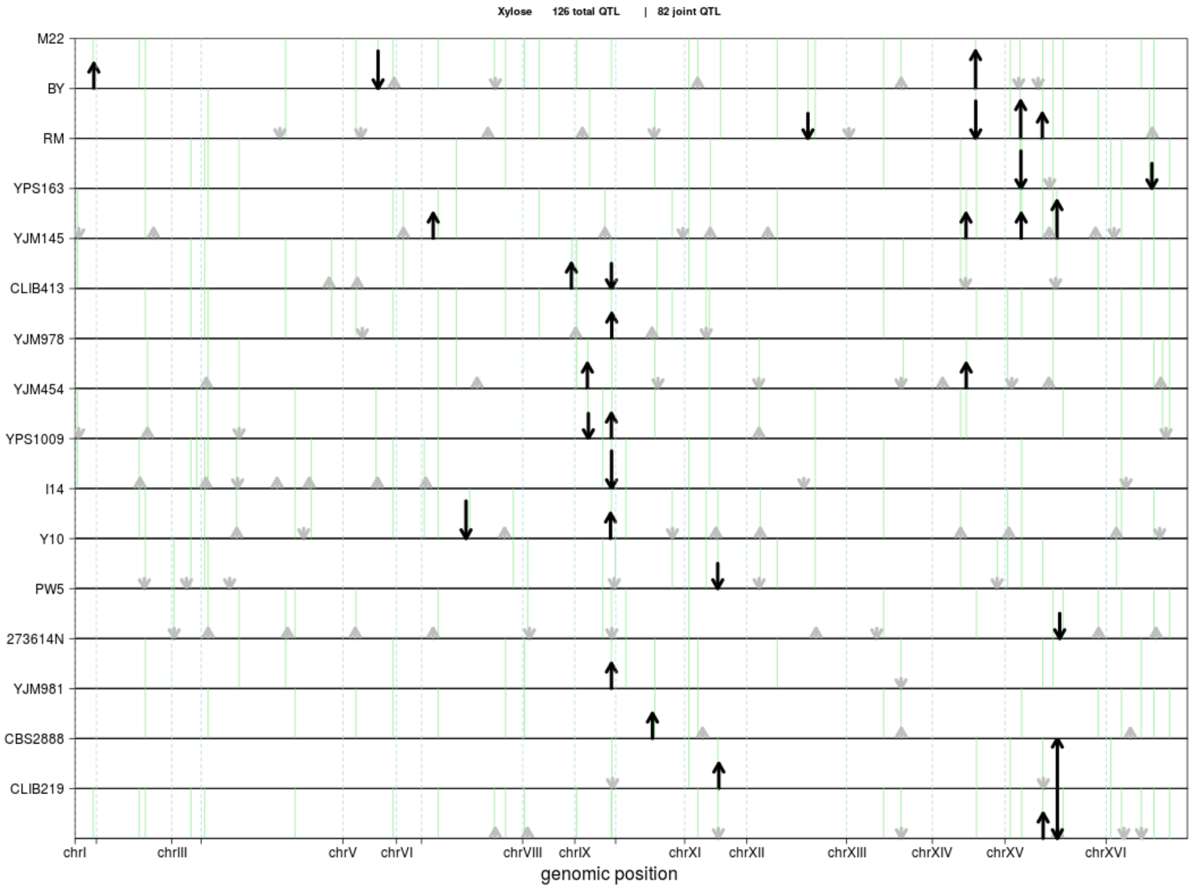

Xylose 126 total QTL | 82 joint QTL
M22
BY
RM
YPS163
YJM145
CLIB413
YJM978
YJM454
YPS1009
I14
Y10
PW5
273614N
YJM981
CBS2888
CLIB219
chrI
chrIII
chrV
chrVI
chrVIII
chrIX
chrXI
chrXII
chrXIII
chrXIV
chrXV
chrXVI
genomic position


324

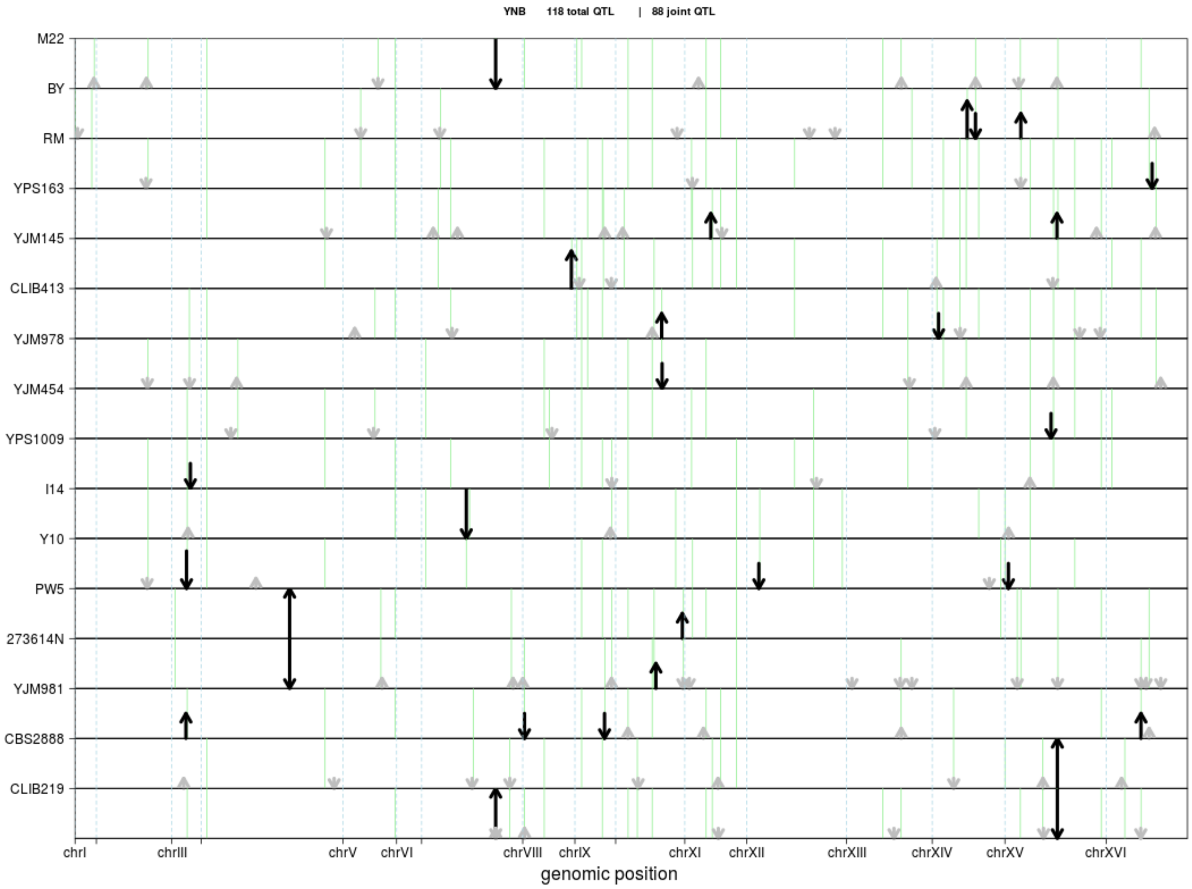
YNB 118 total QTL | 88 joint QTL
M22
BY
RM
YPS163
YJM145
CLIB413
YJM978
YJM454
YPS1009
I14
Y10
PW5
273614N
YJM981
CBS2888
CLIB219
chrI
chrIII
chrV
chrVI
chrVIII
chrIX
chrXI
chrXII
chrXIII
chrXIV
chrXV
chrXVI
genomic position

325

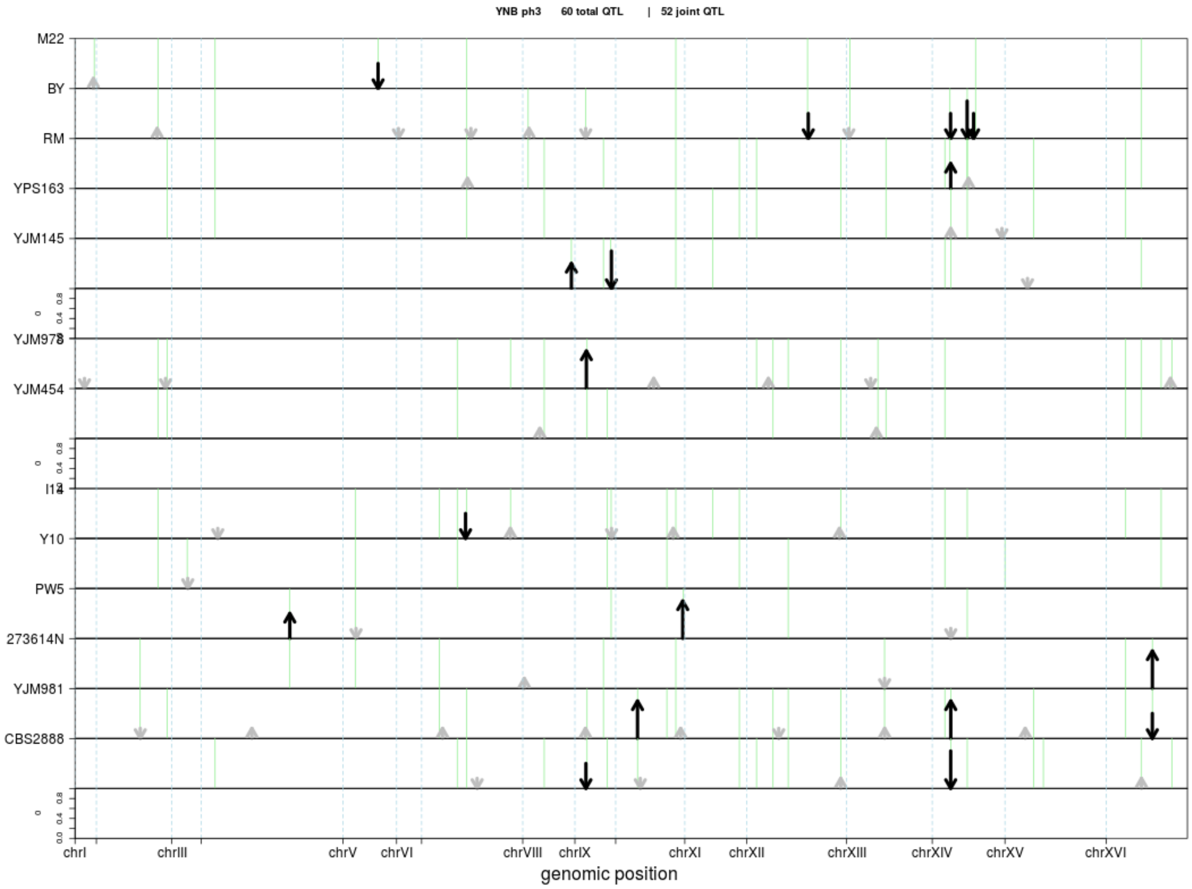

YNB ph3 60 total QTL | 52 joint QTL
M22
BY
RM
YPS163
YJM145
YJM978
YJM454
I14
Y10
PW5
273614N
YJM981
CBS2888
chrI
chrIII
chrV
chrVI
chrVIII
chrIX
chrXI
chrXII
chrXIII
chrXIV
chrXV
chrXVI
genomic position

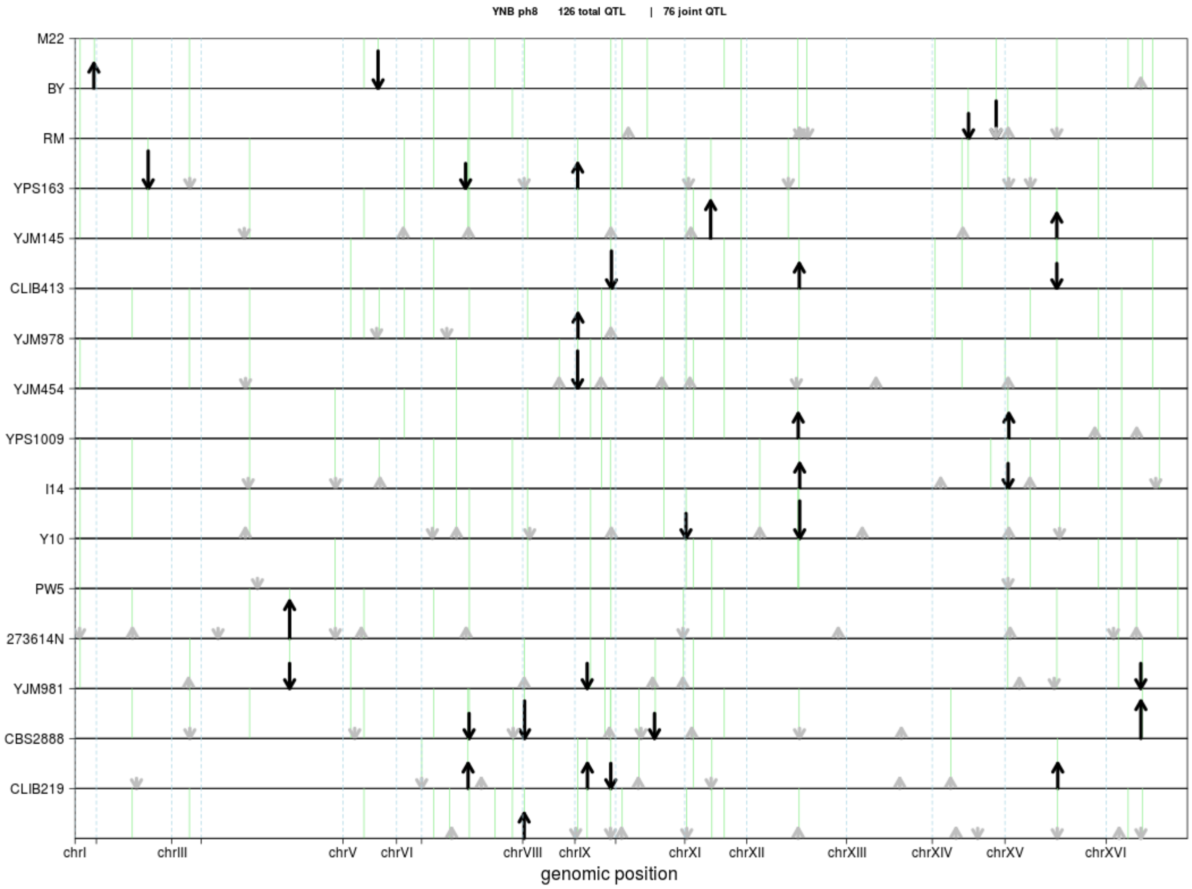
YNB ph8 126 total QTL | 76 joint QTL
M22
BY
RM
YPS163
YJM145
CLIB413
YJM978
YJM454
YPS1009
I14
Y10
PW5
273614N
YJM981
CBS2888
CLIB219
chrI
chrIII
chrV
chrVI
chrVIII
chrIX
chrXI
chrXII
chrXIII
chrXIV
chrXV
chrXVI
genomic position

327

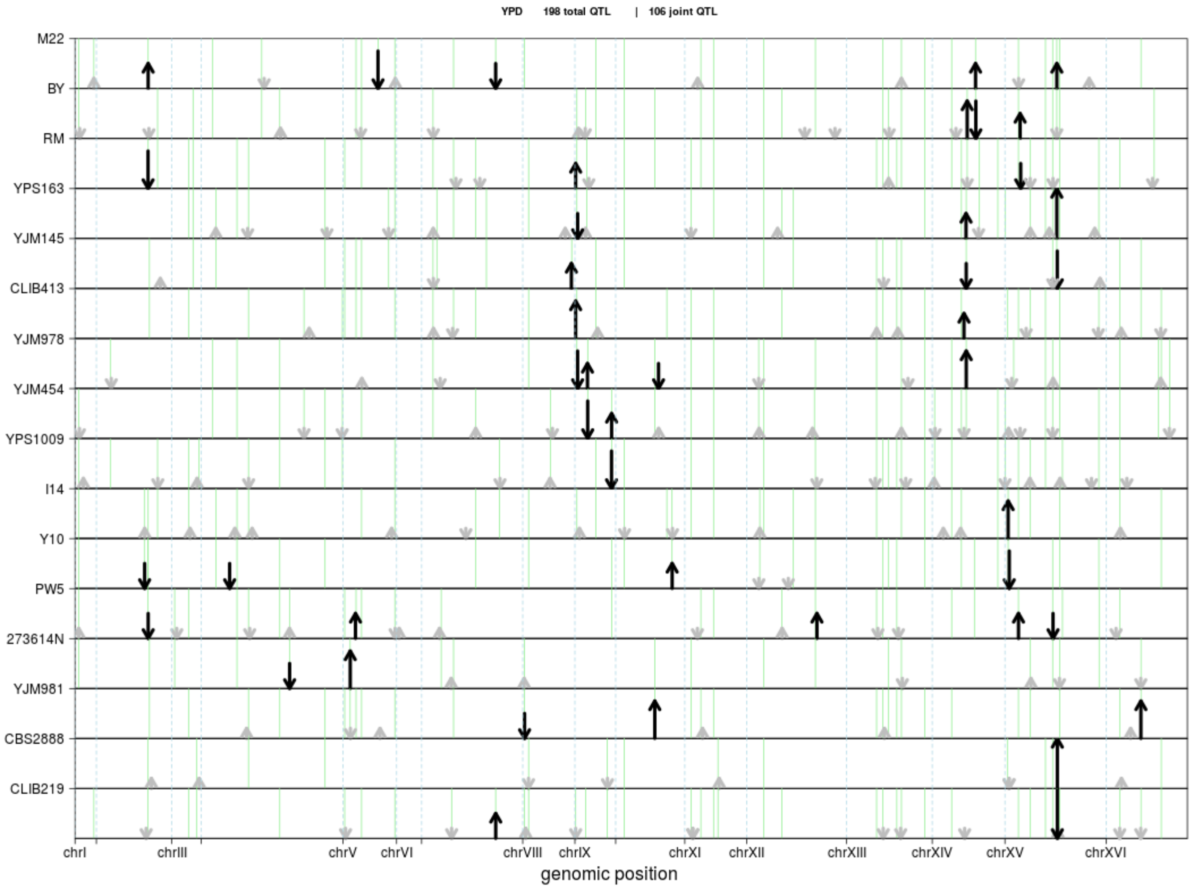
YPD 198 total QTL | 106 joint QTL
M22
BY
RM
YPS163
YJM145
CLIB413
YJM978
YJM454
YPS1009
I14
Y10
PW5
273614N
YJM981
CBS2888
CLIB219
chrI
chrIII
chrV
chrVI
chrVIII
chrIX
chrXI
chrXII
chrXIII
chrXIV
chrXV
chrXVI
genomic position

328

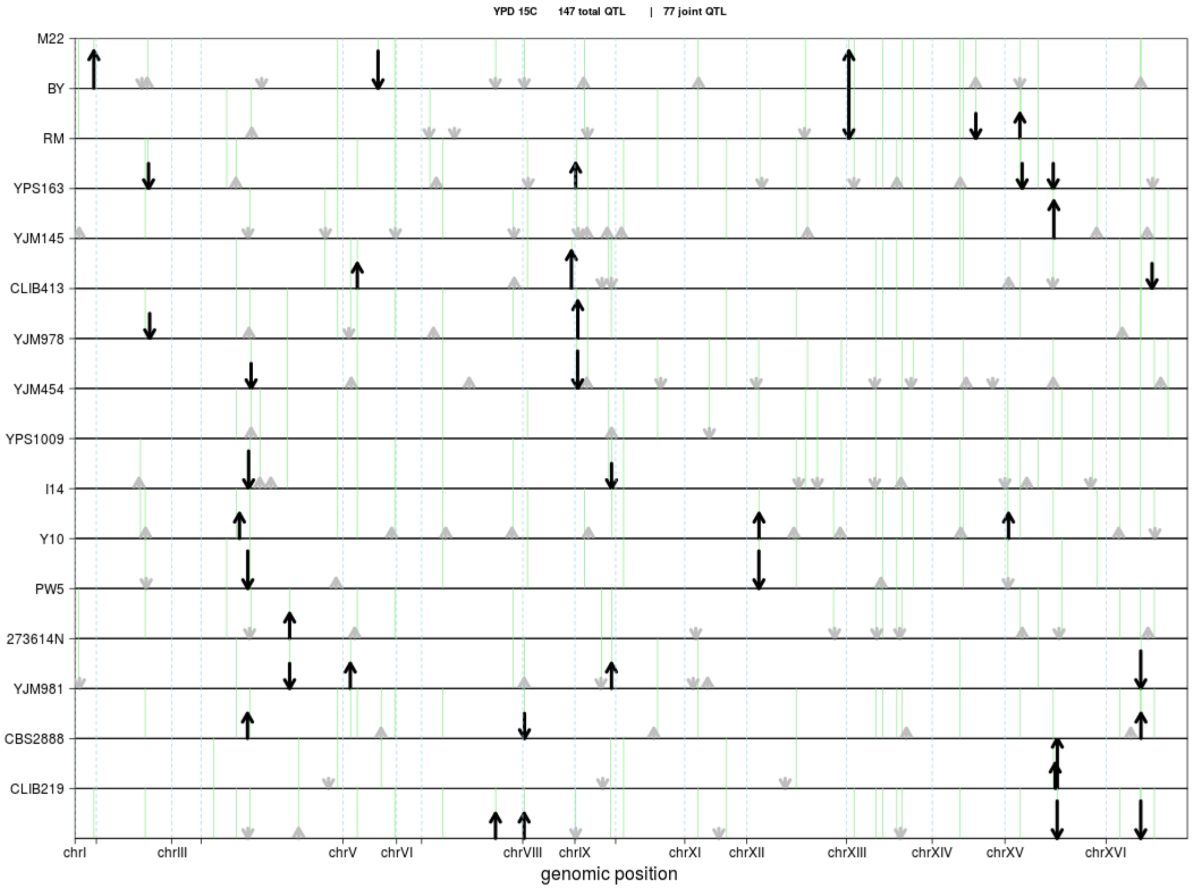
YPD 15C 147 total QTL | 77 joint QTL
M22
BY
RM
YPS163
YJM145
CLIB413
YJM978
YJM454
YPS1009
I14
Y10
PW5
273614N
YJM981
CBS2888
CLIB219
chrI
chrIII
chrV
chrVI
chrVIII
chrIX
chrXI
chrXII
chrXIII
chrXIV
chrXV
chrXVI
genomic position

329

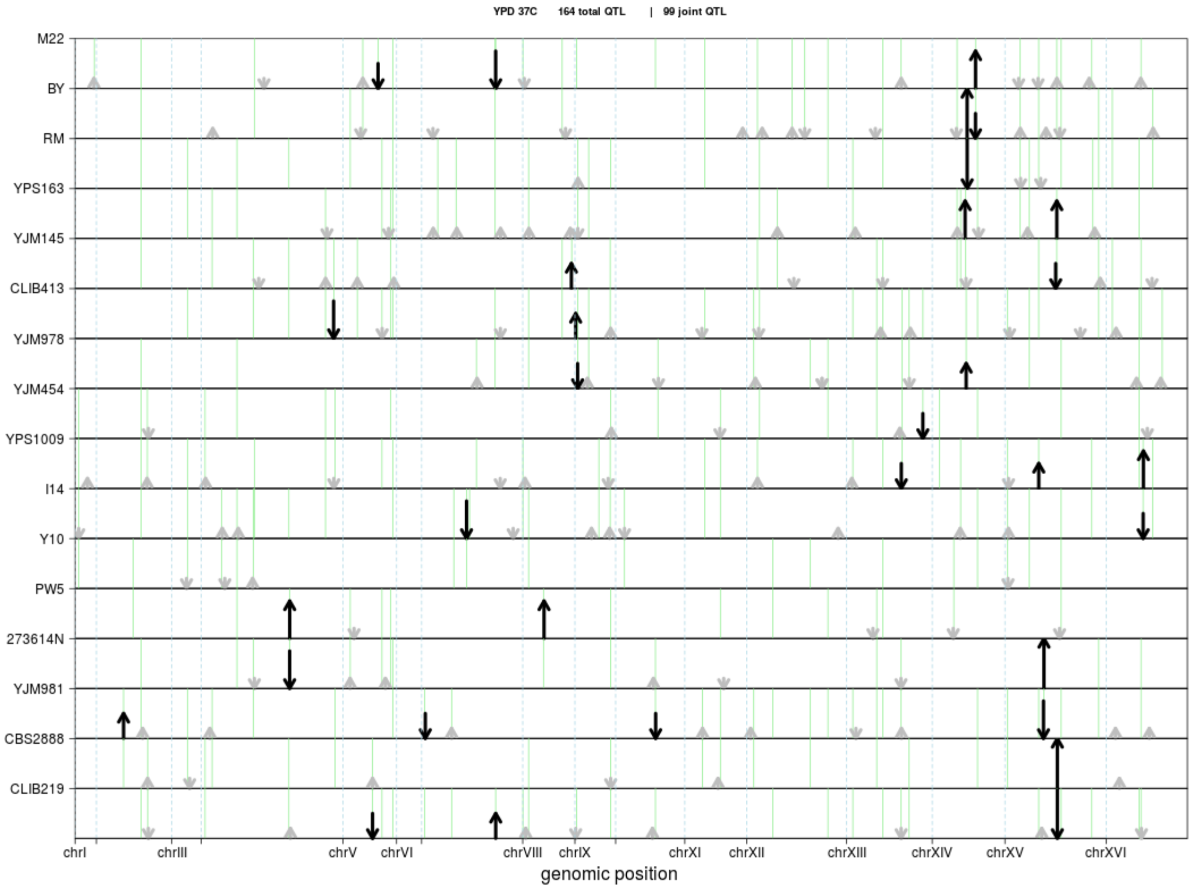


Supplementary Figure 2. Results from QTL mapping are shown for each trait. Each subpanel represents the results for a cross. Along the Y-axis the two parental strains for each cross are shown. Position of QTL along the genome is represented on the X-axis. The arrows represent QTL effects from the within-cross mapping. The arrows point toward the strain that increases growth. The size of the arrow reflects the magnitude of the QTL effect. Full length arrows represent QTL that explain more than 25% of phenotypic variance, ¾ length arrows represent QTL that explain between 8% and 25% of phenotypic variance, ½ length arrows represent QTL that explain between 4% and 8% of phenotypic variance, and short arrows represent QTL that explain less than 4% of phenotypic variance. Large effect QTL (explaining more than 4% of phenotypic variance) are colored black, and small effect QTL (less than 4% of phenotypic variance) are colored grey. The green vertical lines correspond to QTL detected from the joint QTL mapping analysis (Methods).

Supplementary Figure 3

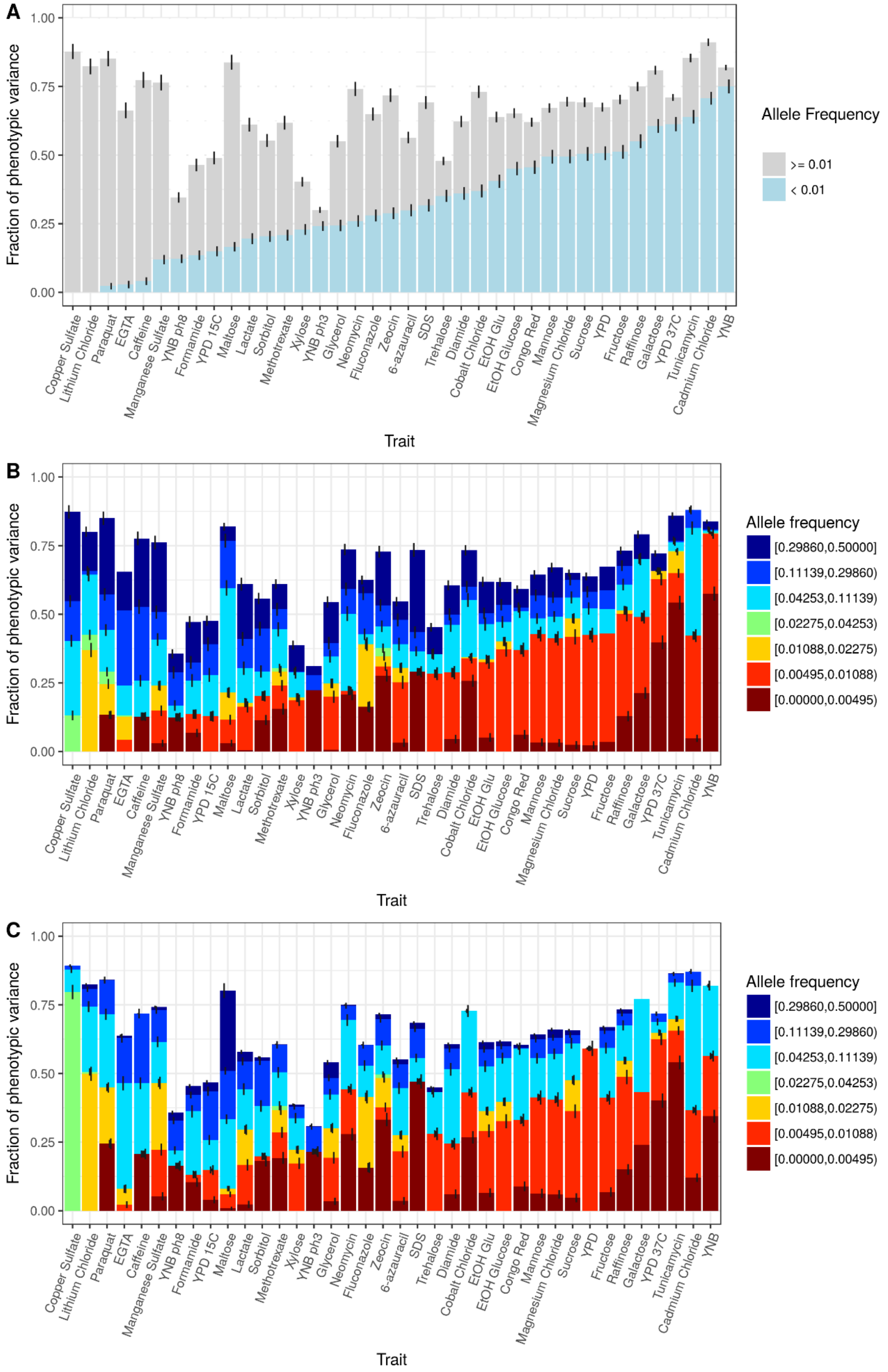

Supplementary Figure 3. (A) Stacked bar plots of a variance component analysis for each trait are shown. Results are the same as in Figure 3a except that the fraction of phenotypic variance, instead of the fraction of total additive variance, is indicated. The variance component model splits the additive genetic variance into the fraction coming from variants with MAF < 1% (blue) and MAF>1% (grey). Error bars show +/- s.e. (B) Same as in (A), except a 7 allele-frequency bin variance component model was fit per trait. (C) Same as in (B), except only private variants (segregating in two crosses) were used in the analysis. This model makes no assumption about the relationship between allele-frequency in the mapping panel and effect sizes.

Supplementary Figure 4.

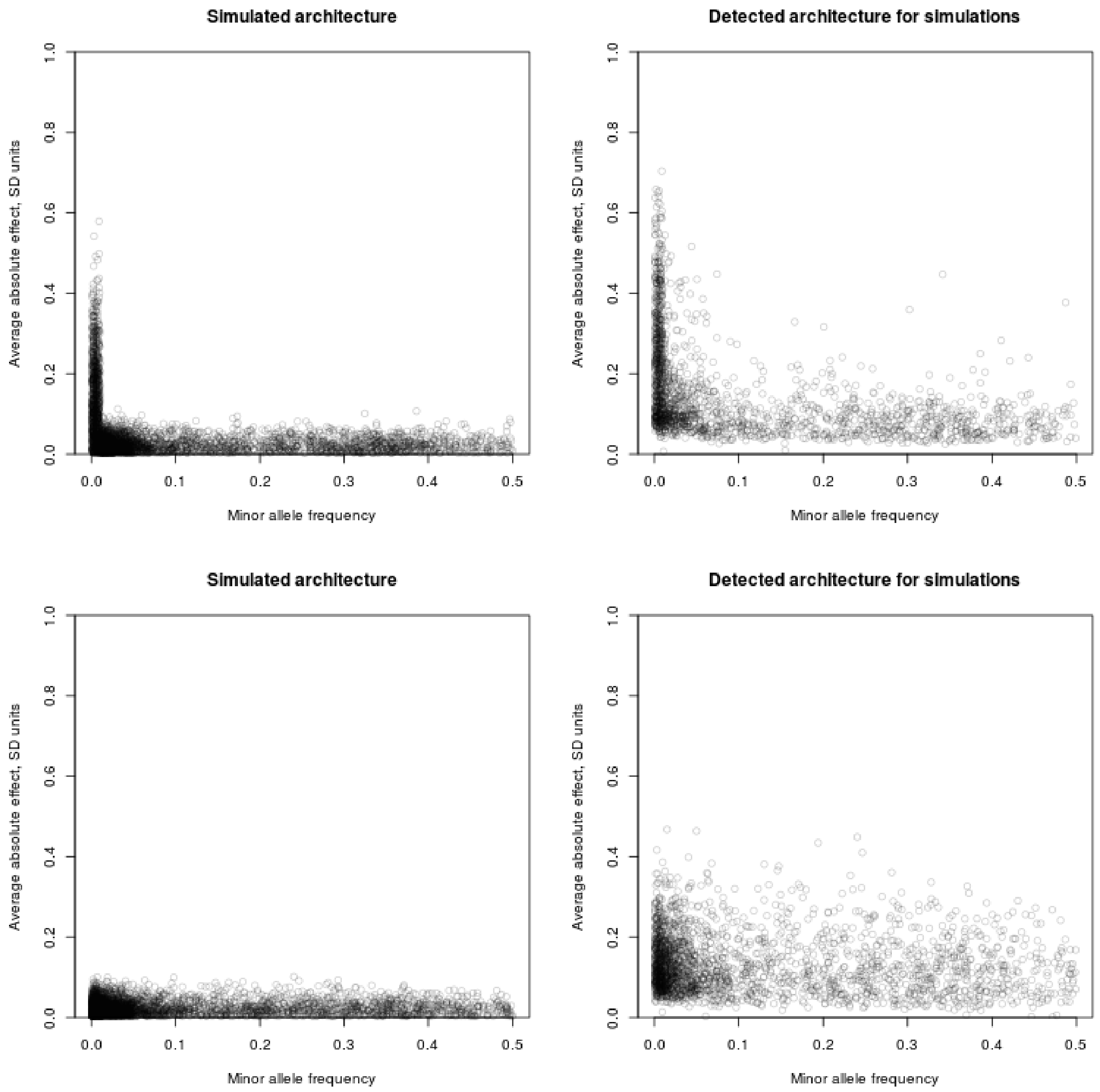

Supplementary Figure 4. Simulated architectures and resulting inference based on lead variants. In the top and bottom leftmost panels the X-axis is the MAF of simulated causal variants, the Y-axis is the simulated average absolute effect of a simulated causal variant. In the top and bottom rightmost panels the X-axis is the MAF of the detected lead variants, and the Y-axis is the estimate of the average absolute effect of the detected lead variants. The top two panels correspond to a simulated architectures of 20 traits, where for each trait 200 causal variants were drawn at random from all variants, effects were sampled from a normal distribution N(0, 0.05), and total additive heritability was 60%. The bottom two panels correspond to a simulated architectures of 20 traits, where for each trait 200 causal variants were drawn at random from all variants, effects were sampled from a normal distribution N(0, 0.05) for variants with MAF>=1% and N(0,0.3) for variants with MAF<1%, and total additive heritability was 60%.

Supplementary Figure 5.

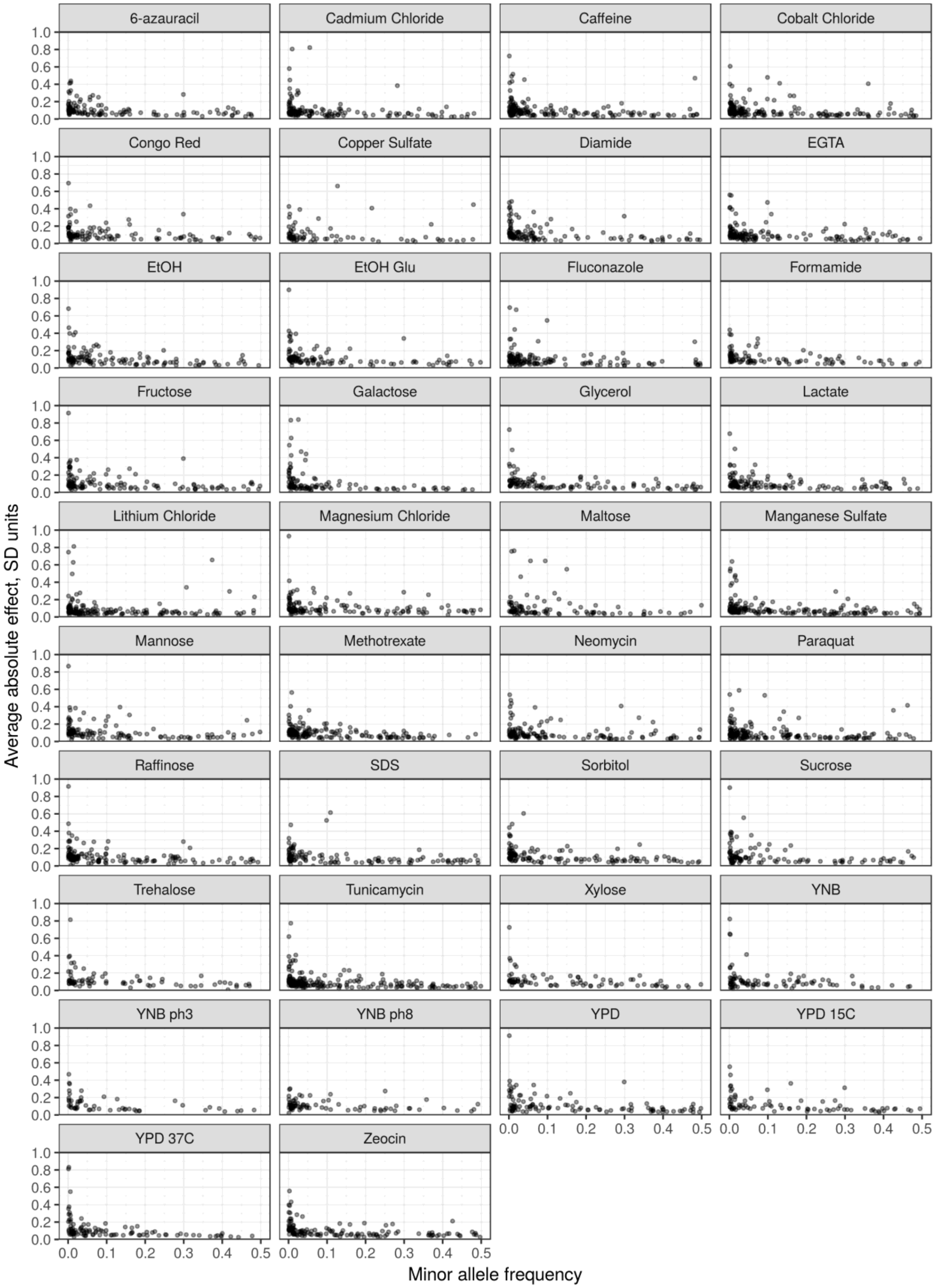

Supplementary Figure 5. The minor allele frequency (X-axis) of the lead variant at each QTL is plotted against QTL effect size as in Figure 3B. Here results are shown separately for each phenotype.

Supplementary Figure 6.

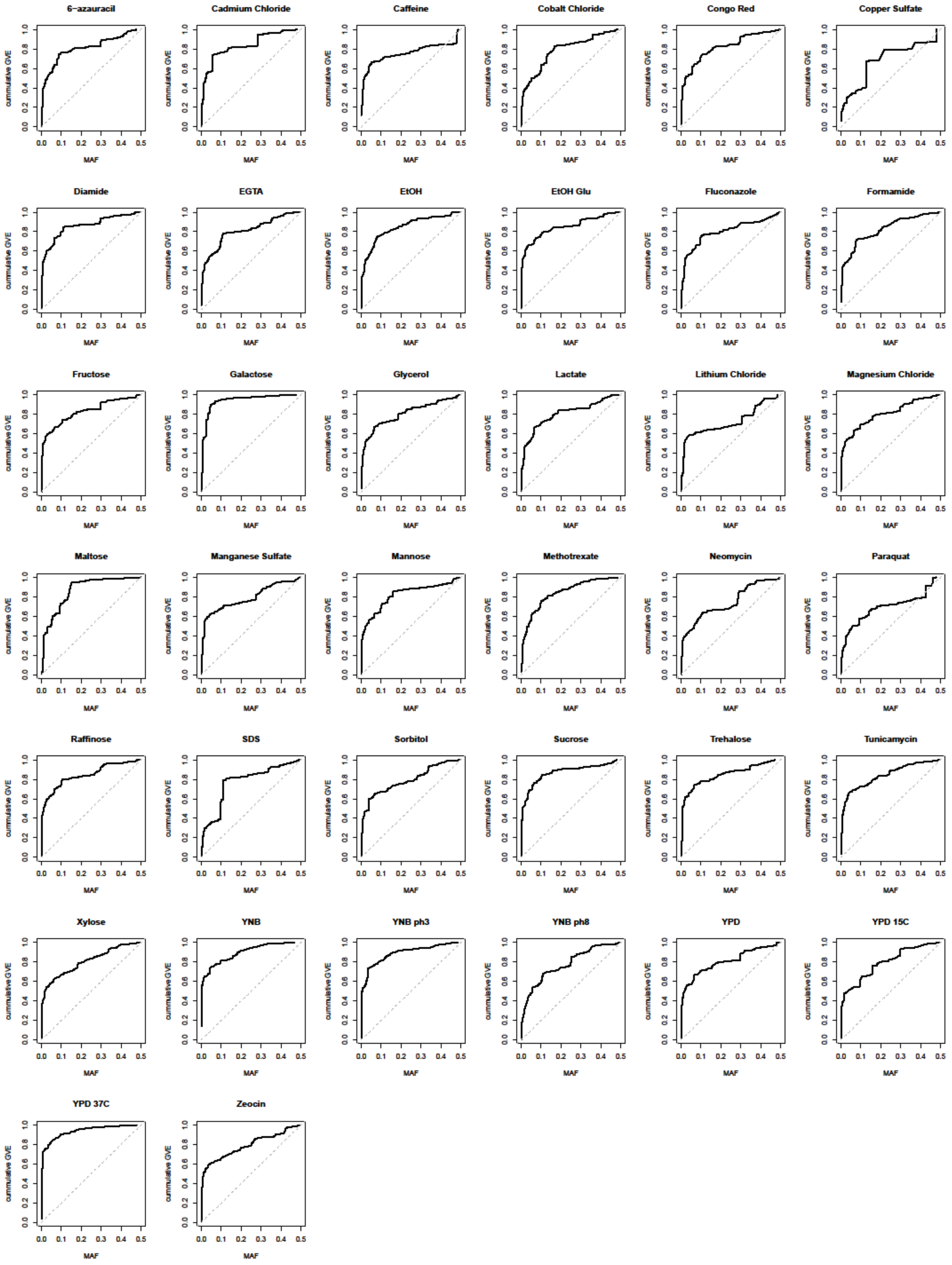

Supplementary Figure 6.

The cumulative genetic variance explained (GVE) in our mapping panel (Y-axis) is plotted against minor allele frequency (X-axis) of the lead variants. Under an evolutionarily neutral model, the cumulative genetic variance explained is linearly proportional to MAF (the identity line MAF=cumulative genetic variance explained). Deviations from the identity are indicative of the action of selective forces.

Supplementary Table 1.

Strain information for the 16 haploid parents and 16 F1 hybrids between them is shown. Additional information about the conditions tested is indicated.

Supplementary Table 2.

Results from within-cross variance components models and total variance explained by the QTL models are shown.

Supplementary Table 3.

QTL mapping results are shown for both the within-cross and the joint analysis.

Supplementary Table 4.

Results for the joint variance component models are given. This includes results for a model with two allele frequency bins (Fig. 3a, Supplementary Fig. 3a), seven allele frequency bins (Supplementary Fig. 3b), and seven allele frequency bins using only variants that are private to each of the 16 parents (Supplementary Fig. 3c).

Supplementary Table 5.

Candidate causal genes per QTL are shown. GO enrichments for causal genes are shown.

#### References


1. Li, H. Aligning sequence reads, clone sequences and assembly contigs with BWA-MEM. *arXiv:1303.3997 [q-bio]* (2013).
2. Engel, S. R. *et al.* The Reference Genome Sequence of Saccharomyces cerevisiae: Then and Now. *G3 (Bethesda)* **4**, 389–398 (2013).
3. Picard Tools - By Broad Institute. Available at: http://broadinstitute.github.io/picard/. (Accessed: 2nd April 2019)
4. Van der Auwera, G. A. *et al.* From FastQ data to high confidence variant calls: the Genome Analysis Toolkit best practices pipeline. *Curr Protoc Bioinformatics* **43**, 11.10.1-33 (2013).
5. Walker, B. J. *et al.* Pilon: an integrated tool for comprehensive microbial variant detection and genome assembly improvement. *PLoS ONE* **9**, e112963 (2014).
6. Layer, R. M., Chiang, C., Quinlan, A. R. & Hall, I. M. LUMPY: a probabilistic framework for structural variant discovery. *Genome Biol.* **15**, R84 (2014).
7. Ebler, J., Schönhuth, A. & Marschall, T. Genotyping inversions and tandem duplications. *Bioinformatics* **33**, 4015–4023 (2017).
8. Bloom, J. S., Ehrenreich, I. M., Loo, W. T., Lite, T.-L. V. & Kruglyak, L. Finding the sources of missing heritability in a yeast cross. *Nature* **494**, 234–237 (2013).
9. Treusch, S., Albert, F. W., Bloom, J. S., Kotenko, I. E. & Kruglyak, L. Genetic Mapping of MAPK-Mediated Complex Traits Across S. cerevisiae. *PLOS Genetics* **11**, e1004913 (2015).
10. Aronesty, E. Comparison of Sequencing Utility Programs. *The Open Bioinformatics Journal* **7**, (2013).
11. Bolger, A. M., Lohse, M. & Usadel, B. Trimmomatic: a flexible trimmer for Illumina sequence data. *Bioinformatics* **30**, 2114–2120 (2014).

12. Albert, F. W., Bloom, J. S., Siegel, J., Day, L. & Kruglyak, L. Genetics of trans-regulatory variation in gene expression. *Elife* **7**, (2018).

13. Arends, D., Prins, P., Jansen, R. C. & Broman, K. W. R/qtl: high-throughput multiple QTL mapping. *Bioinformatics* **26**, 2990–2992 (2010).

14. Pau, G., Fuchs, F., Sklyar, O., Boutros, M. & Huber, W. EBImage--an R package for image processing with applications to cellular phenotypes. *Bioinformatics* **26**, 979–981 (2010).

15. Bloom, J. S. *et al.* Genetic interactions contribute less than additive effects to quantitative trait variation in yeast. *Nature Communications* **6**, ncomms9712 (2015).

16. G'Sell, M. G., Wager, S., Chouldechova, A. & Tibshirani, R. Sequential Selection Procedures and False Discovery Rate Control. *arXiv:1309.5352 [math, stat]* (2013).

17. Churchill, G. A. & Doerge, R. W. Empirical threshold values for quantitative trait mapping. *Genetics* **138**, 963–971 (1994).

18. Clifford, D. & McCullagh, P. The regress package. (2014).

19. Li, H. *et al.* The Sequence Alignment/Map format and SAMtools. *Bioinformatics* **25**, 2078–2079 (2009).

20. Peter, J. *et al.* Genome evolution across 1,011 Saccharomyces cerevisiae isolates. *Nature* **556**, 339–344 (2018).

21. Danecek, P. *et al.* The variant call format and VCFtools. *Bioinformatics* **27**, 2156–2158 (2011).

22. Marçais, G. *et al.* MUMmer4: A fast and versatile genome alignment system. *PLoS Comput. Biol.* **14**, e1005944 (2018).

23. McArdle, B. H. & Anderson, M. J. Fitting Multivariate Models to Community Data: A Comment on Distance-Based Redundancy Analysis. *Ecology* **82**, 290–297 (2001).

24. Kang, H. M. *et al.* Variance component model to account for sample structure in genome-wide association studies. *Nature Genetics* **42**, ng.548 (2010).
25. Forni, S., Aguilar, I. & Misztal, I. Different genomic relationship matrices for single-step analysis using phenotypic, pedigree and genomic information. *Genetics Selection Evolution* **43**, 1 (2011).
26. Yang, J. *et al.* Common SNPs explain a large proportion of the heritability for human height. *Nature Genetics* **42**, 565–569 (2010).
27. Yang, J., Zaitlen, N. A., Goddard, M. E., Visscher, P. M. & Price, A. L. Advantages and pitfalls in the application of mixed-model association methods. *Nat. Genet.* **46**, 100–106 (2014).
28. McMullen, M. D. *et al.* Genetic Properties of the Maize Nested Association Mapping Population. *Science* **325**, 737–740 (2009).
29. Stich, B. Comparison of Mating Designs for Establishing Nested Association Mapping Populations in Maize and Arabidopsis thaliana. *Genetics* **183**, 1525–1534 (2009).
30. Farh, K. K.-H. *et al.* Genetic and epigenetic fine mapping of causal autoimmune disease variants. *Nature* **518**, 337–343 (2015).
31. Storey, J. D. & Tibshirani, R. Statistical significance for genomewide studies. *PNAS* **100**, 9440–9445 (2003).
32. Storey, J. D. The positive false discovery rate: a Bayesian interpretation and the q-value. *Ann. Statist.* **31**, 2013–2035 (2003).
33. Käll, L., Storey, J. D., MacCoss, M. J. & Noble, W. S. Posterior error probabilities and false discovery rates: two sides of the same coin. *J. Proteome Res.* **7**, 40–44 (2008).
34. Alexa, A. & Rahnenfuhrer, J. topGO: Enrichment Analysis for Gene Ontology. (2018).
